# Circadian modulation of neurons and astrocytes controls synaptic plasticity in hippocampal area CA1

**DOI:** 10.1101/666073

**Authors:** John P. McCauley, Maurice A. Petroccione, Lianna Y. D’Brant, Gabrielle C. Todd, Nurat Affinnih, Justin J. Wisnoski, Shergil Zahid, Swasti Shree, Alioscka A. Sousa, Rose M. De Guzman, Rosanna Migliore, Alexey Brazhe, Richard D. Leapman, Alexander Khmaladze, Alexey Semyanov, Damian G. Zuloaga, Michele Migliore, Annalisa Scimemi

**Affiliations:** SUNY Albany, Department of Biology, 1400 Washington Avenue, Albany (NY) 12222, USA; SUNY Albany, Department of Physics, 1400 Washington Avenue, Albany (NY) 12222, USA; Bethlehem Central High School, 700 Delaware Avenue, Delmar (NY) 12054, USA; Federal University of São Paulo, Department of Biochemistry, 100 Rua Tres de Maio, São Paulo (SP) 04044-020, Brazil; National Institute of Biomedical Imaging and Bioengineering, National Institutes of Health, 9000 Rockville Pike, Bethesda (MD) 20892, USA; SUNY Albany, Department of Psychology, 1400 Washington Avenue, Albany (NY) 12222, USA; Institute of Biophysics, National Research Council, 153 Via Ugo la Malfa, Palermo (PA) 90146, Italy; Lomonosov Moscow State University, Department of Biophysics, Leninskie Street 1/12, Moscow 119234, Russia; Department of Molecular Neurobiology, Institute of Bioorganic Chemistry, Miklukho-Maklaya Street 16/10, Moscow 117997, Russia; Sechenov First Moscow State Medical University, Bolshaya Pirogovskaya Street 19c1, Moscow 119146, Russia

**Keywords:** Hippocampus, astrocytes, circadian rhythms, synapses, glutamate, synaptic integration, corticosterone

## Abstract

Most animal species operate according to a 24-hour period set by the suprachiasmatic nucleus (SCN) of the hypothalamus. The rhythmic activity of the SCN is known to modulate hippocampal-dependent memory processes, but the molecular and cellular mechanisms that account for this effect remain largely unknown. Here, we show that there are cell-type specific structural and functional changes that occur with circadian rhythmicity in neurons and astrocytes in hippocampal area CA1. Pyramidal neurons change the surface expression of NMDA receptors, whereas astrocytes change their proximity to synapses. Together, these phenomena alter glutamate clearance, receptor activation and integration of temporally clustered excitatory synaptic inputs, ultimately shaping hippocampal-dependent learning *in vivo*. We identify corticosterone as a key contributor to changes in synaptic strength. These findings identify important mechanisms through which neurons and astrocytes modify the molecular composition and structure of the synaptic environment, contribute to the local storage of information in the hippocampus and alter the temporal dynamics of cognitive processing.

## Introduction

The circadian rhythmicity with which mammals sleep, feed, regulate their body temperature and engage in reproductive behaviors is controlled by the main circadian master clock, the SCN of the hypothalamus, and the coordinated activity of other semi-autonomous ancillary oscillators distributed in the brain and in peripheral tissues (Guilding and Piggins, 2007). The core molecular transcriptional and translational feedback loops that drive circadian rhythmicity in the SCN are fully established during embryonic development (Namihira et al., 1999; Reick et al., 2001; Sumova et al., 2012; Wakamatsu et al., 2001). The endogenous rhythmic activity of the SCN is powerfully entrained by environmental cues, or zeitgebers (ZT), the most potent of which is light. By activating direct retino-hypothalamic glutamatergic connections, light evokes a fast-onset, sustained increase in the firing rate of SCN neurons. The SCN controls the activity of subordinate circadian oscillators through different neuronal projections. Those that remain confined within the medial hypothalamus control the release of corticosteroids into the bloodstream (i.e. glucocorticoids and mineralocorticoids) (Kalsbeek and Buijs, 2002; Lehman et al., 1987; Meyer-Bernstein et al., 1999). A few projections from the SCN reach the inter-geniculate leaflet and paraventricular nucleus of the thalamus and control the rhythmic production of melatonin by the pineal gland (Moore, 1996). Therefore, in nocturnal animals like mice, the production of glucocorticoids, mineralocorticoids and melatonin peaks during the dark phase of the circadian cycle (ZT12-24) (Ishida et al., 2005; Meijer et al., 1998; Moore and Lenn, 1972; Stephan and Zucker, 1972). Corticosteroids and melatonin have important regulatory effects, which take place through the activation of receptors expressed broadly throughout the brain and in peripheral organs (Dubocovich, 2007; Dubocovich et al., 2003; Reppert et al., 1996).

The hippocampus is a subordinate circadian oscillator. Here, more than 10% of genes and proteins show circadian fluctuations (Debski et al., 2017) and are associated with changes in synaptic excitability (Barnes et al., 1977). Learning and memory formation, two functions that are conserved across species and that in mammals rely on the activity of the hippocampal formation, are also modulated by circadian rhythms (Gerstner et al., 2009; Rawashdeh et al., 2018; Ruby et al., 2008; Shimizu et al., 2016; Smarr et al., 2014; Snider et al., 2016). For example, mice trained to context fear conditioning protocols acquire the conditioning more slowly if trained during the dark (D) phase of the circadian cycle (ZT12-24) (Chaudhury and Colwell, 2002), a time when long-term plasticity at Schaffer collateral synapses is also reduced (Winson and Abzug, 1977, 1978). Interestingly, this circadian control of hippocampal function and plasticity is retained in reduced slice preparations, as evidenced by the reduced magnitude of long-term potentiation (LTP) at hippocampal Schaffer collateral synapses in slices prepared during the D-phase (Harris and Teyler, 1983; Raghavan et al., 1999). The persistence of these effects in slices raise the possibility that they are independent of the ongoing activity of extra-hippocampal inputs and might be due to the presence of one or more diffusible factors that are not easily washed out of slices, but whose identity remains enigmatic. Some studies show that increased glucocorticoid levels during the active phase of the circadian cycle facilitate memory formation and cognition (Hui et al., 2004; Whitehead et al., 2013; Yuen et al., 2011). However, other works suggest melatonin is both necessary and sufficient for poor memory formation and plasticity during the D-phase (Ozcan et al., 2006; Rawashdeh et al., 2007). The physiological glucocorticoid in rodents is corticosterone, which acts by binding to intracellular glucocorticoid and mineralocorticoid receptors. Glucocorticoid receptors are expressed at different levels throughout the mouse brain, with the exception of the SCN (Balsalobre et al., 2000; Rosenfeld et al., 1988). The expression of mineralocorticoid receptors is more restricted, but is abundant in the hippocampus, where melatonin receptors are also present (McEown and Treit, 2011; Musshoff et al., 2000; Qi et al., 2013; Venkova et al., 2009). Here we ask what accounts for time-of-day changes in LTP at Schaffer collateral synapses and how neurons, astrocytes and corticosterone contribute to this effect.

Neurons change their propensity to express LTP of excitatory transmission depending on the density, composition and time course of activation of their glutamate receptors (Migliore et al., 2015). What determines the time course for the activation of these receptors is, in part, passive diffusion of glutamate out of the synaptic cleft and, in part, glutamate uptake from astrocytes. Since astrocytes are competent circadian oscillators (Brancaccio et al., 2019; Brancaccio et al., 2017; Prolo et al., 2005), and impaired glutamate uptake degrades LTP (Katagiri et al., 2001; Scimemi et al., 2009), we hypothesize that these cells, in addition to neurons, are fundamental contributors to circadian changes in hippocampal plasticity. Our data show that NMDA receptor activation is significantly reduced during the D-phase, a time when astrocytes retract their fine processes from excitatory synapses and glutamate clearance is prolonged. These changes lead to decreased temporal summation of AMPA EPSPs and reduced LTP during the D-phase. Together, our data provide a new mechanistic understanding of the molecular and cellular mechanisms responsible for the circadian modulation of hippocampal plasticity and memory processing.

## Results

### The hippocampus is a competent circadian oscillator

Central and peripheral circadian clocks are genetically equipped to generate rhythms during embryonic development, but continue their maturation during the first postnatal week. At P10, when synaptogenesis in the SCN and other peripheral oscillators is complete, rats show regular changes in their body temperature that are controlled by the rhythmic activity of circadian oscillators (Sumova et al., 2008; Weinert, 2005). We confirmed that this holds true also in juvenile mice aged P14–21 kept under a 12H:12H L:D schedule, from which we measured the body temperature using micro-transponders implanted under the skin of the shoulder (**Supplementary Fig. 1A**). Their mean body temperature (~35.8°C) displayed consistent oscillations of ±0.5°C every 24.3±5.1 hours, with detectable peaks at the end of the light (L)-phase (ZT12) and troughs at the end of the dark (D)-phase (ZT24). This trend was also present in mice maintained in constant darkness for at least three cycles (12H:12H D:D in red dim light), whose mean body temperature (~36.0°C) showed ±0.8°C oscillations every 24.0±4.0 hours (**Supplementary Fig. 1B**). We confirmed that the expression of the clock genes *Arntl* and *Per2* is not limited to the SCN but also occurs in the CA1 region of the mouse hippocampus. Accordingly, using an RNAscope fluorescent *in situ* hybridization assay (**Fig. 1A-C**), we showed that *Per2* expression increased significantly in sections prepared at ZT15.5 (D-phase) compared to those prepared at ZT3.5 (L-phase) in the SCN (SCN_L_ 2.9e-3±0.6e-3 puncta/μm^2^ (n=28), SCN_D_: 14.8e-3±1.8e-3 puncta/μm^2^ (n=33), ***p=3.5e-7), the pyramidal cell layer of hippocampal area CA1 (CA1-PC_L_: 12.8e-3±1.5e-3 puncta/μm^2^ (n=31), CA1-PC_D_: 19.6e-3±1.4e-3 puncta/μm^2^ (n=36), **p=1.7e-3) and *stratum radiatum* (CA1-SR_L_: 3.1e-3±0.6e-3 puncta/μm^2^ (n=23), CA1-SR_D_: 7.5e-3±0.7e-3 puncta/μm^2^ (n=31) ***p=1.1e-5). By contrast, the expression of *Arntl* did not change significantly between these two time points in any of the brain regions mentioned above (SCN_L_ 4.2e-3±0.5e-3 puncta/μm^2^ (n=28), SCN_D_: 4.9e-3±1.1e-3 puncta/μm^2^ (n=33), p=0.54; CA1-PC_L_: 9.2e-3±1.0e-3 puncta/μm^2^ (n=28), CA1-PC_D_: 7.1e-3±1.0e-3 puncta/μm^2^ (n=31), p=0.16; CA1-SR_L_: 1.9e-3±0.7e-3 puncta/μm^2^ (n=16), CA1-SR_D_: 2.3±0.4 puncta/μm^2^ (n=17), p=0.57). Similar results were obtained in mice kept in constant darkness (**Supplementary Fig. 2A-B**). Here, *Per2* expression increased significantly in sections prepared at ZT15.5 (subjective D-phase) compared to those prepared at ZT3.5 (subjective L-phase) in the pyramidal cell layer of hippocampal area CA1 (CA1-PC_L_: 16.6e-3±1.2e-3 puncta/μm^2^ (n=35), CA1-PC_D_: 23.4e-3±1.8e-3 puncta/μm^2^ (n=32), ***p=9.5e-4) and *stratum radiatum* (CA1-SR_L_: 3.1e-3±0.3e-3 puncta/μm^2^ (n=50), CA1-SR_D_: 4.9e-3±0.4e-3 puncta/μm^2^ (n=33) **p=1.5e-3). By contrast, the expression of *Arntl* did not change significantly (CA1-PC_L_: 11.5e-3±1.0e-3 puncta/μm^2^ (n=35), CA1-PC_D_: 12.9e-3±1.5e-3 puncta/μm^2^ (n=32), p=0.44; CA1-SR_L_: 2.5e-3±0.2e-3 puncta/μm^2^ (n=50), CA1-SR_D_: 3.1±0.3 puncta/μm^2^ (n=33), p=0.10). These findings are consistent with previous works showing that clock gene expression can be detected in the hippocampus of 4-6 month old mice and in hippocampal slice cultures (Kwapis et al., 2018; Namihira et al., 1999; Wang et al., 2009). Our data expands these findings showing that the transcription-translation feedback loops responsible for the circadian regulation of clock gene expression are functional also in the hippocampus of juvenile mice.

**Figure 1.**
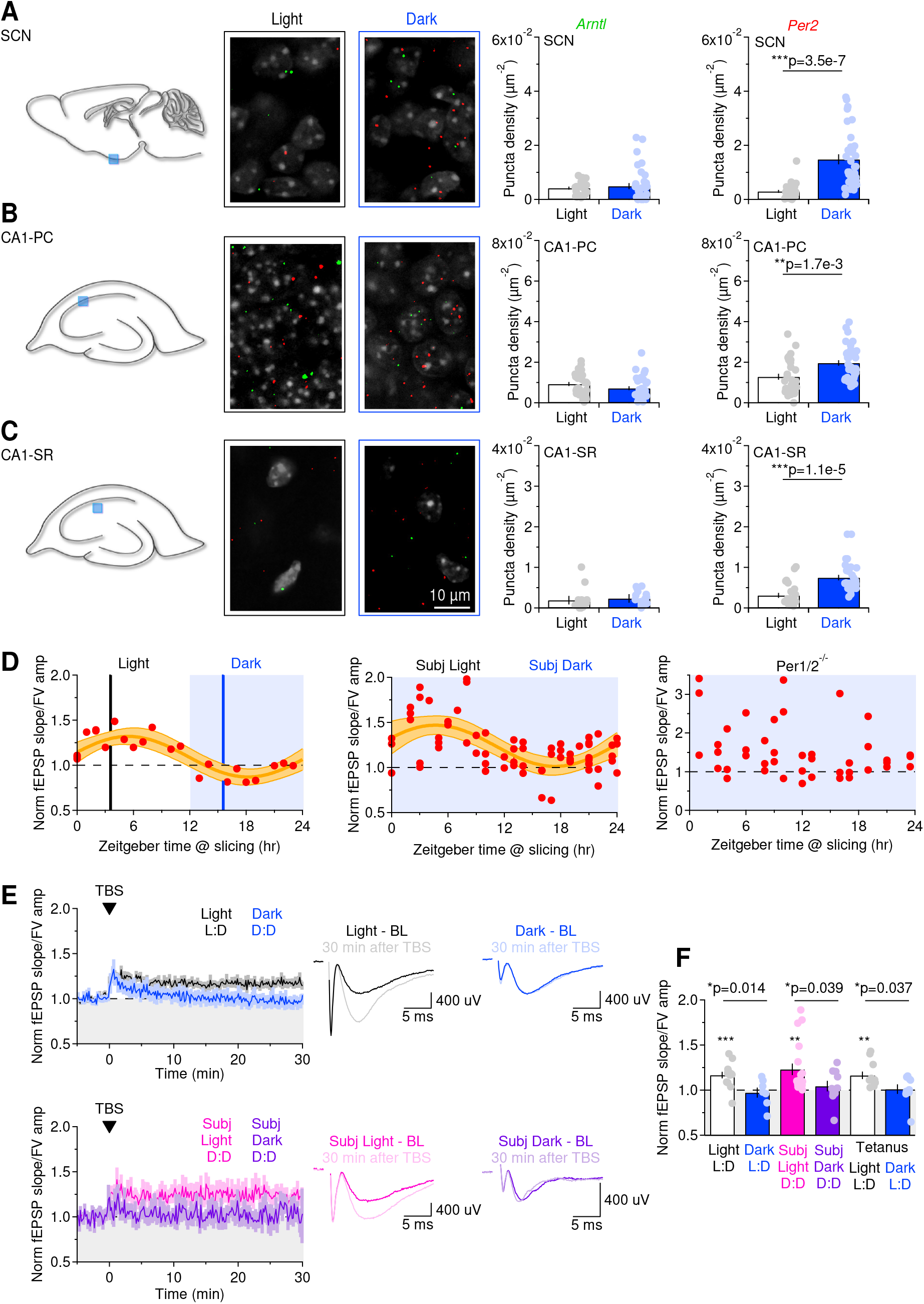
Circadian rhythmicity is present in juvenile mice and is associated with changes in hippocampal plasticity. **(A)** Schematic representation of a mid-sagittal section of the mouse brain. The light blue square represents the location of the SCN region from which we collected confocal images. The green and red puncta represent the fluorescence signal obtained with RNAscope *in situ* hybridization for transcripts of clock genes *Arntl* (green) and *Per2* (red) in slices prepared at ZT3.5 (Light) and ZT15.5 (Dark) in mice kept under 12H:12 L:D conditions. The bar charts on the right provide a summary of the *Arntl* and *Per2* puncta density. **(B)** As in (A), for the pyramidal cell layer of hippocampal area CA1. **(C)** As in (A), for measures obtained in *stratum radiatum*. **(D)** *Left*, Measurements of the baseline-normalized field EPSP slope/FV amplitude in slices prepared at different ZTs. Each red dot represents the result of one experiment performed in hippocampal slices from one mouse. The thick orange line represents the data fitting to a sinusoidal function. The light orange bands represent the 95% confidence bands for the fit. The black and dark blue vertical lines define ZT3.5 and ZT15.5, respectively. Unless otherwise stated, these are times at which we prepared slices representative of the L/D-phases. *Middle*, As on the left panel, for C57BL/6 mice kept under 12H:12H D:D conditions. *Right*, As on the left panel, for Per1/2^-/-^ mice kept under 12H:12H D:D conditions. **(E)** *Top left*, time course of baseline-normalized field EPSP slope/FV amplitude in slices prepared at ZT3.5 (Light, black) and ZT15.5 (Dark, blue). The induction protocol consisted of 10 bursts at 10 Hz (100 Hz and 0.05 s each). *Bottom left*, time course of baseline-normalized field EPSP slope/FV amplitude in slices prepared at ZT3.5 (Subjective Light, magenta) and ZT15.5 (Subjective Dark, purple) from mice kept under constant darkness. *Right*, average responses recorded in each condition 5 min before and 30 min after TBS. Each trace represents the average of 30 consecutive traces, recorded 10 s after each other. The mean fEPSP slope/FV ratio for the representative traces is 1.18 for Light and 0.86 for Dark. The fEPSP slope/FV ratio for the representative traces is 1.17 for subjective Light and 1.07 for subjective Dark. These correspond to the following values obtained by only analyzing the fEPSP slope: Light: 1.96, Dark: 0.95, subjective Light: 1.55, subjective Dark: 1.10). **(F)** Summarized effect of TBS in Light, Dark and constant darkness.

### Circadian rhythmicity shapes long-term plasticity at Schaffer collateral synapses

Previous work showed that the magnitude and incidence of LTP at Schaffer collateral synapses in acute hippocampal slices varies between the L- and D-phase of the circadian cycle (Harris and Teyler, 1983; Raghavan et al., 1999), but the exact time course of this effect was not determined. To do this, we induced LTP using a theta burst stimulation (TBS) protocol that mimics the endogenous electroencephalographic activity of the rodent hippocampus during exploratory learning tasks (Buzsaki, 2002). We applied this protocol to slices prepared at different times of the day (**Fig. 1D**, *left*). Consistent with previous work, the magnitude of LTP changed with circadian rhythmicity, with a period of 25.5 hours. Similar fluctuations in synaptic plasticity were obtained in mice kept under constant darkness, where LTP displayed a period of 25.4 hours, in phase with the one observed in hippocampal slices from mice kept under 12H:12H L:D conditions (**Fig. 1D**, *middle*). No change in the magnitude of LTP across a 24-hour period was detected in arrhythmic Per1/2^-/-^ mice kept in constant darkness (**Fig. 1D**, *right*) (Bae et al., 2001; Zheng et al., 2001). The results were not biased by differences in the age of the mice used for slice preparation (**Supplementary Fig. 3A**). Therefore, the persistence of LTP oscillations in C57BL/6 mice, not Per1/2^-/-^, in constant darkness indicates that they reflect a circadian phenomenon. We selected two time points that were representative of the L-phase (ZT3.5) and D-phase (ZT15.5) to perform the experiments included in the remainder of this work. For simplicity, we refer to slices prepared at ZT3.5 and ZT15.5 as slices prepared during the L- and D-phase, respectively. We detected LTP in slices prepared at ZT3.5 (L-phase) but not in those prepared at ZT15.5 (D-phase; LTP_L_: 1.17±0.03 (n=14), ***p=3.9e-4; LTP_D_: 0.97±0.06 (n=7), p=0.64; LTP_L_ vs LTP_D_: *p=0.014; **Fig. 1E-F**). Similar results were obtained when recording LTP from slices prepared from mice kept in constant darkness for at least three cycles. Here, LTP could be evoked in slices prepared during the subjective L-phase (i.e. ZT3.5; LTP_SL_: 1.23±0.00 (n=19), **p=2.0e-3), not in the subjective D-phase (i.e. ZT15.5; LTP_SD_: 1.04±0.06 (n=10), p=0.47; LTP_SL_ vs LTP_SD_: *p=0.039; **Fig. 1E-F**). There was no correlation between the magnitude of LTP and the stimulus intensity 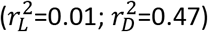 and the results were not influenced by the choice of LTP induction protocol. Accordingly, the magnitude of LTP induced by tetanic stimulation (100 Hz, 1 s) was also larger at ZT3.5 compared to ZT15.5 in mice kept in 12H:12H L:D conditions (LTP_L_: 1.16±0.04 (n=11), **p=3.2e-3; LTP_D_: 1.01±0.05 (n=9), p=0.84; LTP_L_ vs. LTP_D_: *p=0.037; **Fig. 1F**).

In all these experiments, we measured LTP using the ratio between the fEPSP slope and the fiber volley (FV) amplitude. This approach allows us to minimize potential errors in the estimates of LTP that can arise from trial-to-trial variability in the recruitment of pre-synaptic afferents, due to the stochastic nature of voltage-gated sodium and potassium channel gating (O’Donnell and van Rossum, 2014) and the highly non-linear effects this has on neurotransmitter release (Dodge and Rahamimoff, 1967). This approach also allows us to minimize any potential confounding factor due small fluctuations in the tip resistance of the recording electrode or to variability in the number of afferent fibers recruited in different slice preparations. To confirm that this approach did not introduce factual errors in our measure of LTP, we repeated the analysis of the experiments shown in **Fig. 1D-F** using the fEPSP slope as a measure of LTP (**Supplementary Fig. 3A-E**; LTP_L_: 1.47±0.08 (n=13), ***p=4.8e-5; LTP_D_: 0.94±0.06 (n=7), p=0.37; LTP_L_ vs. LTP_D_: ***p=4.4e-5; LTP_SL_: 1.52±0.08 (n=14), ***p=2.0e-5; LTP_SD_: 0.93±0.08 (n=9), p=0.42; LTP_SL_ vs. LTP_SD_: ***p=3.8e-5). The results of this analysis are in agreement with those obtained using the fEPSP slope/FV amplitude ratio. They show that the magnitude of LTP varies with a period of ~24.1 hours and LTP can be detected at ZT3.5 but not at ZT15.5 in mice kept under 12h:12h L:D and D:D conditions. For these reasons, we used the fEPSP slope/FV amplitude ratio to measure LTP in the remainder of this work.

We asked whether the circadian modulation of synaptic plasticity at Schaffer collateral synapses could be due to daily fluctuations in extracellular levels of D-Serine, a co-agonist of NMDA receptors (Papouin et al., 2017). To address this, we recorded NMDA EPSCs from CA1 pyramidal cells, evoked by Schaffer collateral stimulation, before and after application of a saturating concentration of D-Serine (100 μM). Consistent with our own previous works (Scimemi et al., 2004; Scimemi et al., 2009), D-Serine did not potentiate the NMDA EPSC amplitude (88±1% of baseline (n=26); **Supplementary Fig. 3F**). These data indicated that changes in the activation of the NMDA receptor glycine-binding sites have no effect on the circadian modulation of LTP in our experiments.

### The surface expression of NMDA receptors on CA1 pyramidal cells varies with circadian rhythmicity

We performed experiments to determine whether the loss of LTP during the D-phase could be accounted by changes in pre- or post-synaptic function (**Fig. 2**). We alternated single and paired electrical stimuli (10-20 V, 50 μs) to evoke glutamate release from Schaffer collaterals. We recorded AMPA EPSCs using a Cs^+^-based internal solution while keeping CA1 pyramidal cells at a holding potential of −65 mV, in the presence of the GABA_A_ receptor antagonist picrotoxin (100 μM). After recording a stable baseline of AMPA EPSCs, we blocked them by bath applying the AMPA receptor antagonist NBQX (10 μM) and switched the holding potential to +40 mV to record NMDA EPSCs. Our measures of AMPA and NMDA EPSC paired-pulse ratio (PPR) remained similar during the L/D-phases, suggesting that there was no change in pre-synaptic release probability (AMPA PPR_L_: 2.7±0.1 (n=19), AMPA PPR_D_: 2.7±0.1 (n=13), p=0.98; NMDA PPR_L_: 2.3±0.2, NMDA PPR_D_: 2.6±0.3, p=0.62; **Fig. 2A**). In these experiments, the stimulus strength was set to evoke AMPA EPSCs of similar amplitude during the L/D-phases (AMPA EPSC amp_L_: 72±7 pA, AMPA EPSC amp_D_: 93±12 pA, p=0.15). Their rise time (AMPA EPSC rise_L_: 1.9±0.1 ms, AMPA EPSC rise_D_: 2.2±0.2 ms, p=0.15) and half decay time (AMPA EPSC t_50L_: 10.0±0.4 ms, AMPA EPSC t_50D_: 10.7±0.8 ms, p=0.47) were also similar, suggesting no change in the time course of AMPA receptor activation when delivering single stimuli to Schaffer collaterals (**Fig. 2C**). The amplitude of the NMDA EPSCs, however, was smaller in the D-phase (NMDA EPSC amp_L_: 94±15 pA, NMDA EPSC amp_D_: 47±7 pA, **p=8.2e-3), with no change in their rise and half decay time (NMDA EPSC rise_L_: 5.6±0.2 ms, NMDA EPSC rise_D_: 5.8±0.4 ms, p=0.75; NMDA EPSC t_50L_: 99±7 ms, NMDA EPSC t_50D_: 101±7 ms, p=0.85; **Fig. 2D**). The reduced NMDA EPSC amplitude led to reduced NMDA/AMPA ratio in the D-phase, an effect that had not been previously described (NMDA/AMPA_L_: 2.2±0.3, NMDA/AMPA_D_: 1.0±0.2, **p=3.7e-3; **Fig.2E**). We confirmed these results in separate experiments in which RuBi-glutamate (100 μM) was superfused through the recording chamber and uncaged using a 5 ms duration light pulse generated by a white LED light source connected to the epifluorescence port of the microscope (2.7 mW at the specimen plane). We recorded electrically-evoked and flash-evoked AMPA and NMDA EPSCs in the same cells (**Fig. 2F-G**). Once again, only the amplitude of electrically-evoked NMDA EPSCs was smaller during the D-phase (AMPA EPSC amp_L_: 110±17 pA (n=5), AMPA EPSC amp_D_: 73±19 pA (n=5), p=0.18; NMDA EPSC amp_L_: 40±7 pA, NMDA EPSC amp_D_: 13±5 pA, *p=0.017; **Fig. 2F**) as was that of flash-evoked NMDA EPSCs, not AMPA EPSCs (AMPA Flash-EPSC amp_L_: 80±16 pA, AMPA Flash-EPSC amp_D_: 112±25 pA, p=0.31; NMDA Flash-EPSC amp_L_: 254±42 pA, NMDA Flash-EPSC amp_D_: 69±28 pA, ***p=7.8e-3; **Fig. 2G**). These effects were not associated with gross changes in the density of excitatory synaptic contacts onto CA1 pyramidal cells, as indicated by the presence of a similar spine density in dendrites from biocytin-filled neurons in slices prepared during the L/D-phases (L: 0.64±0.03 (n=85), D: 0.65±0.02 (n=105), p=0.90; **Fig. 2B**).

**Figure 2.**
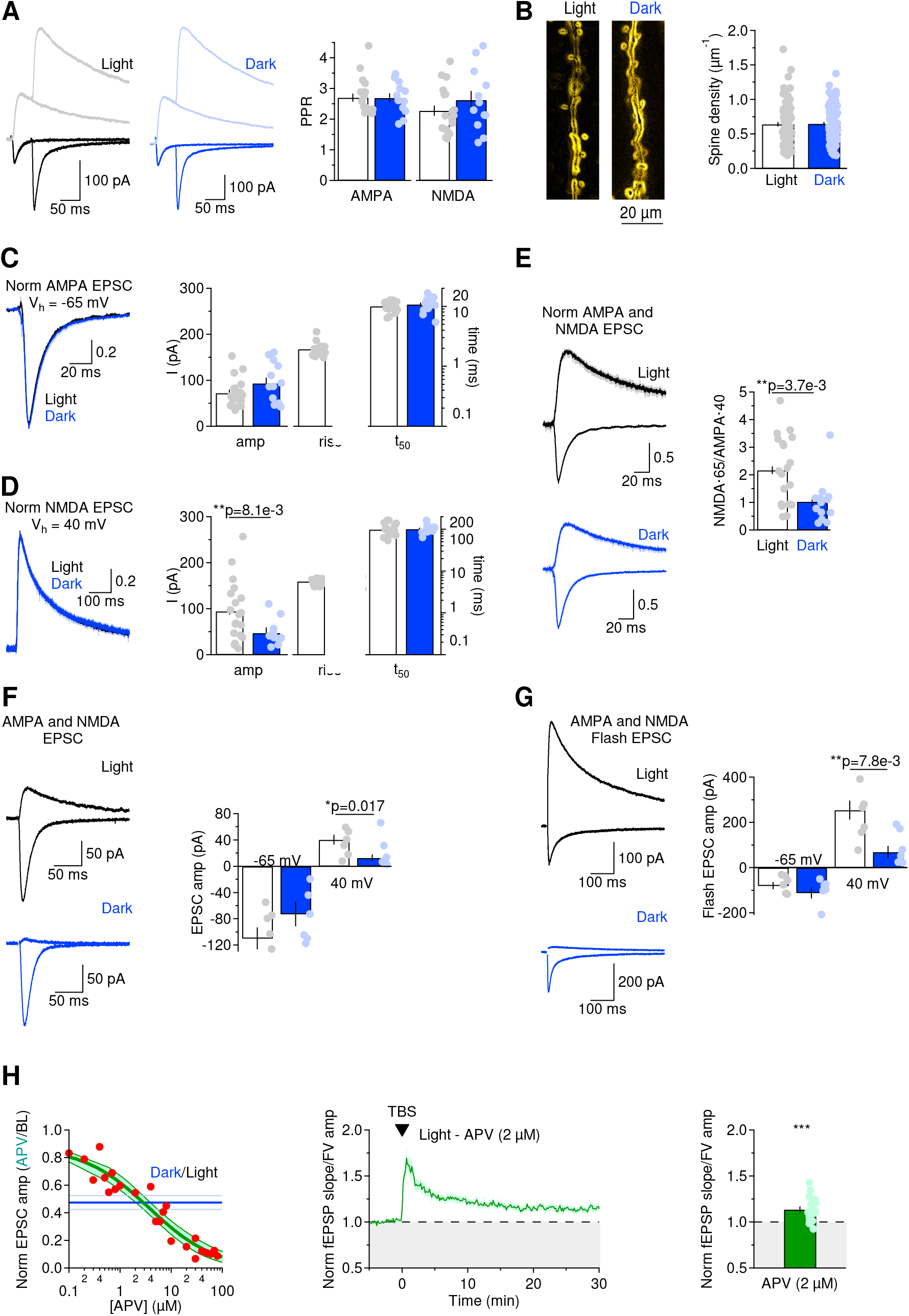
NMDA receptor activation in CA1 pyramidal cells is reduced during the dark phase. **(A)** Example of single and paired AMPA and NMDA EPSCs, recorded at a holding potential of −65 mV and 40 mV, respectively. Each trace represents the average of 20 stimulations, delivered with a 10 s inter-stimulus interval. Summary graph of the paired-pulse ratio (PPR) for AMPA and NMDA EPSCs. **(B)** Biocytin-filled dendritic branches of CA1 pyramidal cells in slices prepared at ZT3.5 (light) and ZT15.5 (dark). The confocal images were acquired as confocal z-stacks using a 63X/1.4 NA oil objective. They were subsequently converted into maximum intensity projections and processed with a Canny-Deriche filtering for edge detection using Fiji. The summary of spine linear density shows no significant difference between samples fixed during the light and dark phase. **(C)** Peak-normalized AMPA EPSCs recorded during the L/D-phases, with summary of the amplitude and kinetics analysis of AMPA EPSCs. No significant difference in the amplitude, rise and half-decay time was detected during the L/D-phases. **(D)** Peak-normalized NMDA EPSCs recorded during the L/D-phases. The summary of the amplitude and kinetics analysis of NMDA EPSCs shows that the amplitude of NMDA EPSCs was smaller during the D-phase. **(E)** Average AMPA and NMDA EPSCs, normalized by the peak of the AMPA current in each cell. The NMDA/AMPA ratio, scaled by their corresponding driving force, was reduced in the D-phase. **(F)** Average electrically-evoked AMPA and NMDA EPSCs recorded from CA1 pyramidal cells in the presence of RuBi-glutamate (100 μM). The amplitude of electrically-evoked NMDA, not AMPA EPSCs, was smaller in slices prepared at ZT15.5. **(G)** Average flash-evoked AMPA and NMDA EPSCs recorded from CA1 pyramidal cells in response to RuBi-glutamate uncaging (100 μM). The amplitude of uncaging-evoked NMDA, not AMPA EPSCs, was smaller in slices prepared at ZT15.5. **(H)** *Left*, Titration curve showing the effect of increasing concentrations of APV on the amplitude of NMDA EPSCs recorded in slices prepared in the L-phase. Each red dot represents the result of one experiment. The thick green line and shaded areas represent the Hill fit of the data and 95% confidence bands, respectively. The horizontal blue and light blue lines represent the reduction of NMDA receptor activation detected in the D-phase (see panel E). APV (2 μM) reduces the NMDA EPSC amplitude by 53%. *Middle*, Time course of baseline-subtracted ratio between the field EPSC slope and the fiber volley amplitude in slices prepared at ZT3.5 (light), in the presence of APV (2 μM). *Right*, Statistical summary, showing that LTP in the L-phase is not occluded by APV (2 μM).

If the loss of LTP during the D-phase was due to reduced NMDA receptor expression (~53%), we should block LTP in the L-phase by blocking NMDA receptors to the same extent. To do this, we recorded NMDA EPSCs from slices prepared at ZT3.5, in the presence of increasing concentrations of the competitive, high-affinity NMDA receptor antagonist APV (**Fig. 2H**, *left*). APV (2 μM) blocked the NMDA EPSC amplitude by ~53%. We then applied the TBS protocol in the presence of APV (2 μM), to activate Schaffer collateral synapses in hippocampal slices prepared at ZT3.5 (L-phase). Despite the partial block of NMDA receptors, we could still induce LTP (1.14±0.03 (n=19), ***p=1.6e-4; **Fig. 2H**, *right*), suggesting that other mechanisms, in addition to loss of NMDA receptors, contribute to the circadian modulation of synaptic plasticity at Schaffer collateral synapses.

### Circadian changes in astrocyte glutamate clearance affect the temporal summation of AMPA EPSCs

The results presented so far point to the existence of regulatory mechanisms that alter NMDA receptor activation and LTP expression in CA1 pyramidal cells. This circadian reduction in the synaptic weight of NMDA currents on its own is not expected to change the summation of AMPA or NMDA EPSCs evoked by temporally clustered stimuli, like those that compose each burst of the LTP-inducing TBS stimulation. Surprisingly, however, our experiments showed that the summation of AMPA EPSCs was reduced in the D-phase (**Fig. 3A**, *top*) whereas the peak NMDA EPSC amplitude recorded when delivering trains of five pulses at 100 Hz to CA1-PCs was similar during the L/D-phases (**Fig. 3A**, *bottom*). Using a kinetic model of AMPA and NMDA receptors, we showed that these effects could be recapitulated by a prolongation in the lifetime of glutamate in the extracellular space, which would prolong the recovery time of AMPA receptors from the open to the closed state (**Fig. 3B**). This hypothesis was supported by the results obtained using a multi-compartmental NEURON model, which takes into account the complex morphology of CA1 pyramidal cells and the random spatial distribution of excitatory inputs through the apical dendritic tree in *stratum radiatum* (**Fig. 3C,D**). The model confirmed that the smaller AMPA EPSC summation in the D-phase could be accounted for by a 3-fold increase in the recovery rate of the peak conductance of AMPA receptors in the D-phase (from 10 to 30 ms). The small reduction in NMDA EPSC summation in the D-phase (**Fig. 3A**) could be accounted for by a small, 15% reduction in the peak conductance of NMDA receptors, which does not lead to a significant change in the temporal summation of consecutive NMDA EPSPs (**Fig. 3D**).

**Figure 3.**
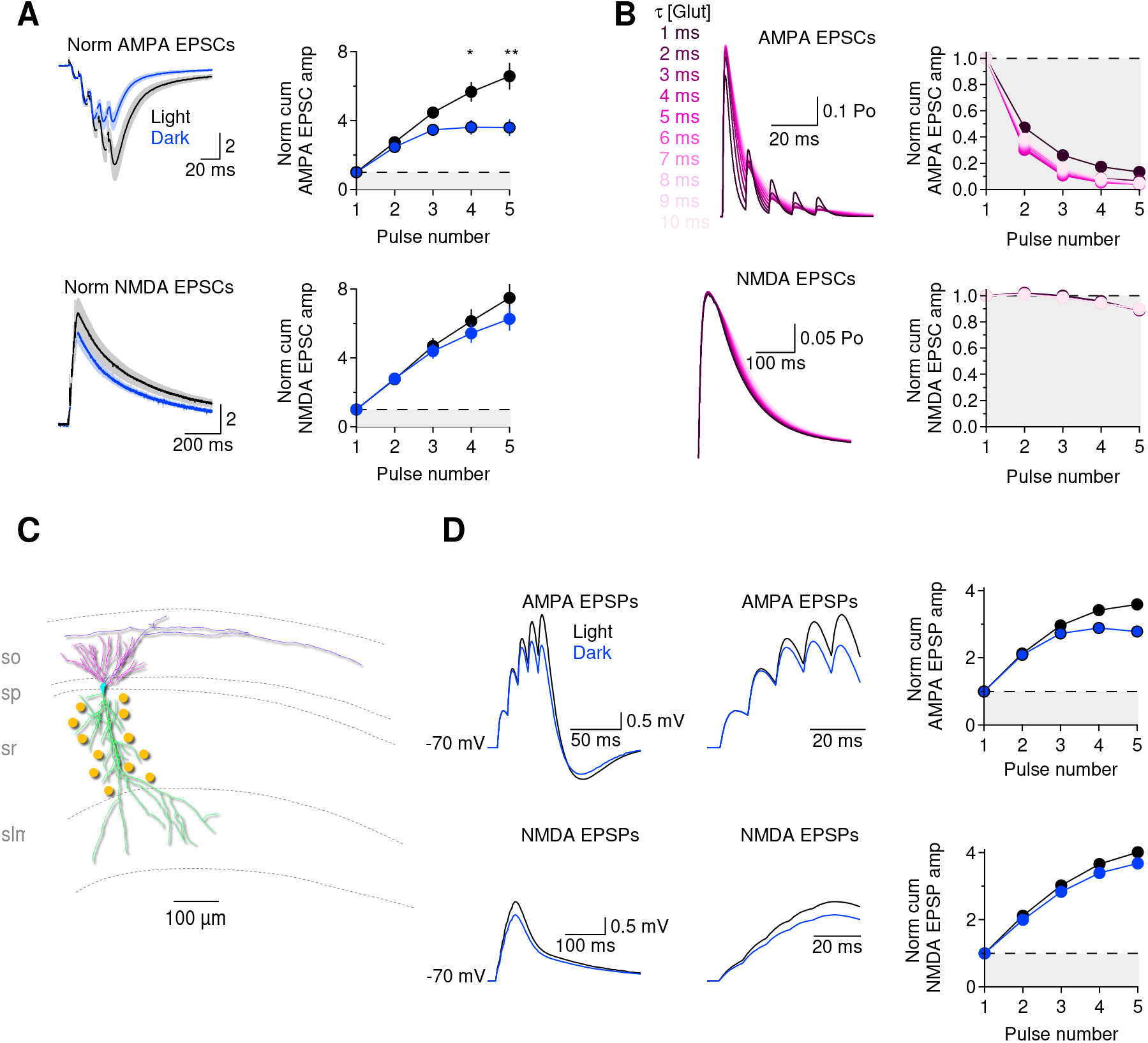
Temporal summation of AMPA EPSCs is reduced during the D-phase. **(A)** *Top*, train of AMPA EPSCs recorded in voltage clamp mode while holding pyramidal cells at a holding potential of −65 mV. Each train stimulation was composed of five pulses, with an inter-pulse interval of 10 ms. The AMPA EPSC summation was reduced in the D-phase. *Bottom*, as the panels described on the top, for NMDA EPSCs recorded at a holding potential of 40 mV. NMDA EPSC summation is similar during the L/D-phases. **(B)** *Top*, ChanneLab simulations of AMPA EPSCs evoked by five consecutive glutamate release events, with a peak glutamate concentration of 1 mM and an inter-event interval of 10 ms. The kinetic scheme of AMPA receptors was taken from (Jonas et al., 1993). The decay time of each glutamate transient (τ) was color-coded according to the scheme on the left hand side of the figure. Longer glutamate transients lead to reduced AMPA EPSC summation. *Bottom*, as the top panels, for NMDA EPSCs. The kinetic scheme for NMDA receptor activation was taken from (Lester and Jahr, 1992). Longer glutamate transients did not lead to significant changes in NMDA EPSC summation. **(C)** Morphology of the biocytin-filled CA1 pyramidal cell neuron used to run the NEURON simulations (soma: cyan; basal dendrites: magenta; apical dendrites: green; axon: purple; synaptic inputs in *stratum radiatum*: yellow). **(D)** Results of NEURON multicompartmental model. *Top*, the traces represent somatic AMPA EPSP evoked by activating a cohort of 100 synapses randomly distributed through the dendritic tree of CA1 pyramidal cells. Changes in the recovery rate of the AMPA receptors, mimicking those induced by prolonged glutamate clearance, reduced the temporal summation of consecutive EPSPs. The second panel of this figure shows the evoked EPSPs on an expanded time scale. *Bottom*, somatic change in potential evoked by activating the same cohort of excitatory synapses that in this case only contained NMDA receptors. To mimic the experimental results obtained during the D-phase, we reduced the NMDA conductance at each synaptic contact. The summary graphs on the right hand side provide an overview of the normalized cumulative AMPA and NMDA EPSP amplitude recorded in modeling conditions that mimic experimental data collected during the L/D-phases.

We performed a number of simulations, in which we changed the conductance for a variety of ion channels, as a further test to determine if these could explain the changes in the temporal summation of EPSPs between the L- and D-phases. We examined the role of high and low voltage activated calcium channels (HVA and LVA, respectively), hyperpolarization-activated cation channels (I_h_) and A-type potassium channels (I_KA_; **Supplementary Fig. 4**). According to the model, a reduction in the temporal summation of EPSPs could occur if there was a 250-fold increase in the current density of HVA and LVA channels (**Supplementary Fig. 4A,C**). However, no significant change in the amplitude and voltage dependence of HVA and LVA calcium currents was detected experimentally in CA1-PCs (**Supplementary Fig. 4B,D**). Likewise, a 3.3-fold increase in I_h_ or a 5-fold increase in I_KA_ could decrease EPSP summation (**Supplementary Fig. 4E,G**), but these changes were not detected experimentally (**Supplementary Fig. 4F,H**). In CA1-PCs from slices prepared during the L- and D-phase 4-aminopyridine (4AP, 50 μM), an antagonist of I_KA_ channels, increased the action potential duration without altering the action potential peak and latency. Because these effects were similar in slices prepared during the L- and D-phase, we concluded that there are no detectable circadian changes in the expression of I_KA_ channels in CA1-PCs (**Supplementary Fig. 8I-L**). A 35-fold increase in the magnitude of delayed rectifier potassium currents (I_KDR_) or a 25-fold increase in calcium-activated potassium currents (I_KCa_) could also reduce the EPSP summation, but these effects are also expected to change the action potential waveform between the Land D-phase. Since this was not the case, based on the results of our experiments, it is unlikely that any of these effects accounts for the changes in the temporal summation of excitatory synaptic inputs observed in the D-phase. Therefore, circadian changes in the temporal summation of excitatory inputs in CA1 pyramidal cells cannot be attributed to changes in active ion channel conductances, leading to the possibility that these might result from changes in glutamate clearance and receptor activation.

If there is a prolongation in the lifetime of synaptically-released glutamate in the extracellular space during the D-phase, then the time course of synaptically-activated glutamate transporter currents (STCs) in astrocytes should be slower. As previously shown, stimulation of excitatory Schaffer collaterals evokes glutamate release in *stratum radiatum* and causes the onset of a complex current waveform in astrocytes (**Fig. 4A**) (Diamond, 2005; Scimemi and Diamond, 2013; Scimemi et al., 2009; Sweeney et al., 2017). Because of the low resistance of the astrocytes’ cell membrane, in whole-cell patch clamp recordings, the first transient outward current corresponds to the extracellular fiber volley, a readout of action potentials travelling along Schaffer collaterals. This fiber volley is not followed by a field current generated by glutamate receptor activation, because all recordings were performed in the presence of AMPA and GABA_A_ receptor blockers (NBQX 10 μM, picrotoxin 100 μM). What follows the fiber volley is an inward current representing the overlay of a sustained potassium current, through which astrocytes remove potassium ions accumulated in the extracellular space from action potential firing in neurons, and a glutamate transporter-mediated current (i.e. the STC), generated by the coordinated movement of glutamate with Na^+^, K^+^ and H^+^ across the membrane (Zerangue and Kavanaugh, 1996). As expected, small changes in the stimulation strength across cells led to proportionate changes in the amplitude of the fiber volley and of the potassium current (**Fig. 4B**). The range of stimulus intensity used in our experiments evoked STCs of similar amplitude and rise time, across the L/D-phases (STC amp_L_: 25.0±3.1 pA (n=7), STC amp_D_: 23.8±3.1 pA (n=7), p=0.79; STC rise_L_: 1.5±0.2 ms (n=7), STC rise_D_: 1.7±0.2 ms (n=7), p=0.34; **Fig. 4C**). However, STCs decayed more slowly in the D-phase, as evidenced by measuring the half decay time (t_50L_: 8.0±0.3 ms (n=7), t_50D_: 11.9±0.8 (n=7), **p=2.2e-3) and the centroid (<*t*>_L_: 8.5±0.4 ms (n=7), <*t*>_D_: 14.3±1.3 ms (n=7), **p=3.6e-3; **Fig. 4C**). The kinetics of the STC are similar to those of the facilitated portion of the STC (fSTC), obtained by subtracting STCs evoked by single and paired stimulation (100 ms inter-pulse interval) (Diamond, 2005; Scimemi and Diamond, 2013; Scimemi et al., 2013; Scimemi et al., 2009; Sweeney et al., 2017). A low concentration of the broad spectrum glutamate transporter antagonist TFB-TBOA (1 μM) reduced the fSTC amplitude (norm fSTC amp_L_: 0.43±0.05 (n=7) ***p=1.9e-5, norm fSTC amp_D_: 0.49±0.06 (n=7), ***p=9.5e-5, L vs D: p=0.47) and prolonged its kinetics similarly during the L/D-phases (norm fSTC rise_L_: 1.3±0.1 **p=2.3e-3, norm fSTC rise_D_: 1.1±0.1, ***p=8.7e-4, L vs D: p=0.066; norm fSTC t_50L_: 1.2±0.1, **p=7.8e-3, norm fSTC t_50D_: 1.1±0.1, *p=0.017, L vs D: p=0.070; norm fSTC <*t*>_L_ 1.2±0.1, *p=0.02, norm fSTC <*t*>_D_: 1.1±0.1, *p=0.029, L vs D: p=0.097; **Fig. 4D**). The fSTCs are useful because they can be used to derive the time course of glutamate clearance from astrocytes (Diamond, 2005; Scimemi and Diamond, 2013; Sweeney et al., 2017). This analysis indicated that the time of glutamate clearance from astrocytic membranes is slower during the D-phase (<*t*>_clearanceL_: 7.4±0.4 ms (n=7), <*t*>_clearanceD_: 9.0±0.5 ms (n=7), L vs D: *p=0.035; **Fig. 4E,F**), suggesting that the glutamate uptake capacity of these cells varies physiologically at different times of the day.

**Figure 4.**
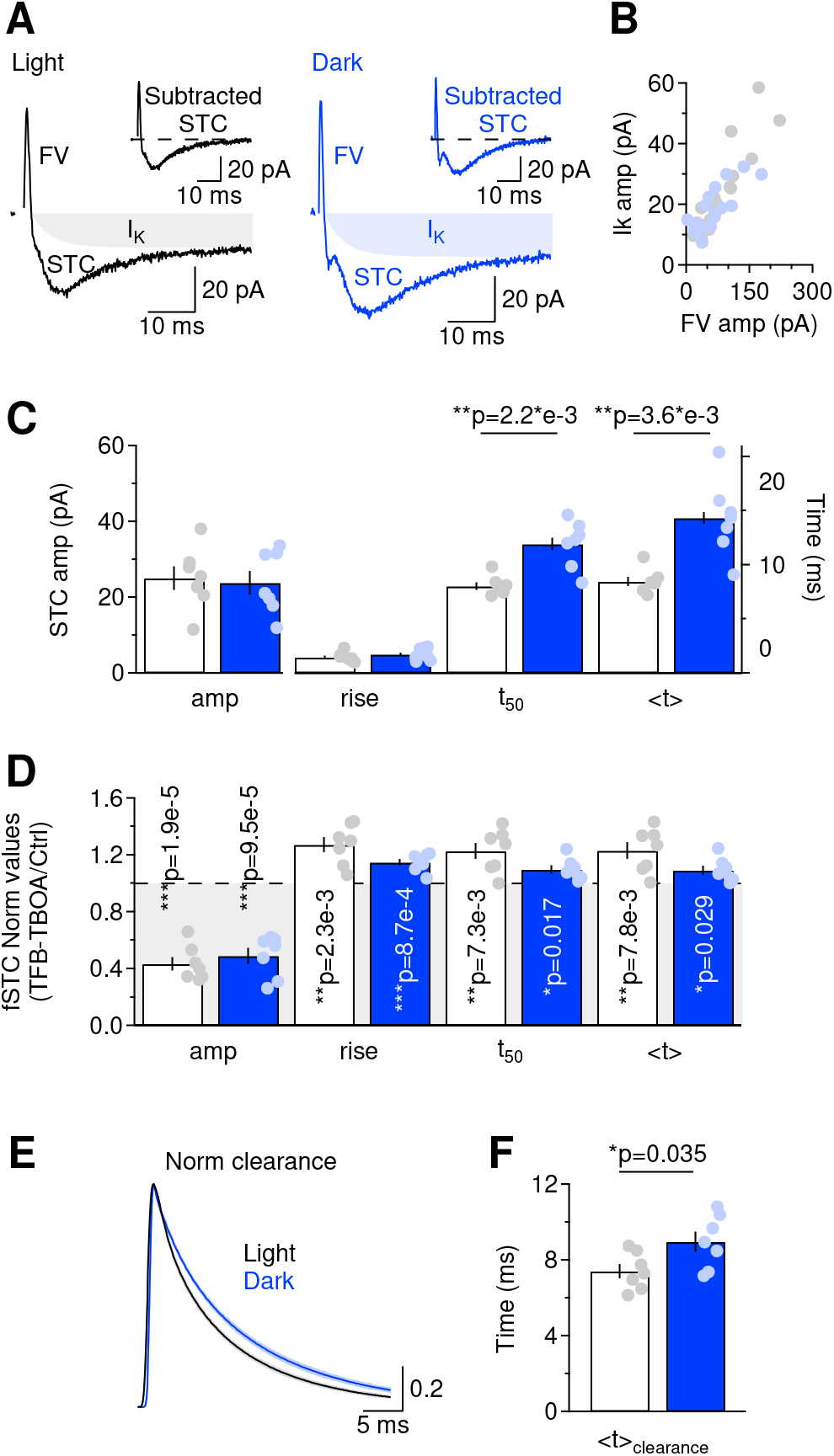
Glutamate clearance from the extracellular space is slower during the D-phase. **(A)** Example of whole-cell recordings from astrocytes obtained in voltage-clamp mode, in the presence of the GABA_A_ and AMPA receptor antagonists picrotoxin (100 μM) and NBQX (10 μM). Extracellular stimuli to Schaffer collaterals evoked a brief fiber volley (FV) followed by a fast-decaying STC and a slow-decaying potassium current (I_K_). **(B)** The amplitude of the sustained potassium current was proportional to the amplitude of the evoked fiber volley and this relationship remained unaltered across the L/D-phases. **(C)** Summary graph of the amplitude and kinetics of STCs evoked in slices prepared during the L/D-phases. We used extracellular stimulus strengths that evoked STCs of similar amplitude during the L/D-phases. The evoked STCs displayed similar 20-80% rise time but their half-decay time (t_50_) and centroid (<*t*>) were prolonged during the D-phase. **(D)** The facilitated portion of the STC (fSTC) was isolated by subtracting single from paired STCs and was used to estimate the time course of glutamate clearance from astrocytes (Scimemi and Diamond, 2013). The high-affinity, broad-spectrum glutamate transporter antagonist TFB-TBOA (1 μM) reduced the fSTC amplitude significantly and to a similar extent during the L/D-phases. The fSTC rise, half-decay time and centroid were larger in the presence of TFB-TBOA. The similar effects of TFB-TBOA on slices prepared during the L/D-phases was an important pre-requisite for the deconvolution analysis of fSTCs. **(E)** Average astrocyte glutamate clearance waveforms during the L/D-phases (mean±SEM). **(F)** Summary graph showing that glutamate clearance by astrocytes is slower during the D-phase.

These effects were not confounded by changes in the passive membrane properties of astrocytes across the L/D-phases (**Supplementary Fig. 5**). The I/V plots, obtained by measuring the steady-state current response to 100 ms long voltage steps (**Supplementary Fig. 5A**), showed a similar profile during the L/D-phases (**Supplementary Fig. 5B**). The membrane resistance (R_mL_: 4.3±0.3 MOhm (n=35), R_mD_: 5.2±0.3 MOhm (n=29), p=0.07; **Supplementary Fig. 5C,D**) and capacitance were also similar during the L/D-phases (C_mL_: 13.3±1.2 pF (n=35), C_mD_: 12.7±1.5 pF (n=29), p=0.74; **Supplementary Fig. 5C,D**). These values were significantly lower than the membrane resistance and capacitance of CA1 pyramidal cells (R_mL_: 202.2±20.0 MOhm (n=15), ***p=3.1e-7, R_mD_: 176.3±15.6 MOhm (n=13), ***p=1.3e-7; C_mL_: 60.0±4.9 pF (n=15), ***p=8.9e-8, C_mD_: 57.7±6.5 pF (n=13), ***p=1.3e-5), which also did not differ significantly during the L/D-phases (R_mL_ vs R_mD_ p=0.32; C_mL_ vs C_mD_ p=0.78; **Supplementary Fig. 5E**). The astrocytes’ resting membrane potentials remained similar during the L/D-phases (V_rL_: −82±4 mV (n=9), V_rD_: −81±4 mV (n=6), p=0.88; **Supplementary Fig. 5F**), as did their level of dye coupling, measured via confocal imaging of biocytin-filled cells (coupling_L_: 12±2 (n=11), coupling_D_: 12±1 (n=7), p=0.98; **Supplementary Fig. 5F**).

The slower glutamate clearance in the D-phase was not due to reduced expression of the astrocyte glutamate transporters GLAST and GLT-1, which we measured in protein extracts from mouse hippocampi dissected at ZT3.5 or ZT15.5 using Western blotting (**Fig. 5A,B**). Glutamate transporters assemble as trimers that can only be partially broken into monomers and dimers during SDS-PAGE. Thus, the total band intensity of GLAST monomers and dimers, normalized to the band intensity of our loading control β-actin, was 1.49±0.13 (n=20) during the L-phase and 1.33±0.17 (n=20) during the D-phase (p=0.47), representing a non-significant 11±8% reduction of GLAST expression during the D-phase (p=0.30; **Fig. 5A**). Similarly, the total normalized band intensity of GLT-1 was 0.77±0.08 (n=16) during the L-phase and 0.78±0.06 (n=16) during the D-phase (p=0.90), leading to a non-significant 3±6% change of expression during the D-phase (p=0.78; **Fig. 5B**). A potential limitation of these experiments lies in the fact that the Western blot analysis is performed on all membrane-associated proteins, including those in the membrane of intracellular organelles. In reality, only glutamate transporters incorporated into the plasma membrane contribute to extracellular glutamate clearance. To address this concern, we used an *in vitro* surface biotinylation assay to measure the relative expression of GLAST and GLT-1 in slices prepared in the L- and D-phase, according to previously established methods (Thomas-Crusells et al., 2003) (**Supplementary Fig. 6**). This approach provided a reliable tool to separate transporters expressed in the plasma membrane from those trafficking through intracellular organelles, as indicated by the absence of biotinylated GLAST/GLT-1 in the cytoplasmic fraction of our protein extracts (GLAST_L_: 0.01±0.02 (n=7), GLAST_D_: −0.01±0.02 (n=7), p=0.48; GLT-1_L_: 0.10±0.07 (n=5), GLT-1_D_: 0.02±0.01 (n=5), p=0.33; **Supplementary Fig. 6B,D**). The data showed that the total normalized band intensity of GLAST was 1.05±0.17 (n=21) during the L-phase and 0.80±0.12 (n=19) during the D-phase (p=0.23), which accounted for a non-significant 7±13% change in GLAST plasma membrane expression during the D-phase (p=0.66; **Supplementary Fig. 6A**). The total normalized band intensity of GLT-1 was 1.16±0.23 (n=15) during the L-phase and 0.96±0.20 (n=13) during the D-phase (p=0.52), which accounted for a non-significant 20±12% change in GLT-1 plasma membrane expression during the D-phase (p=0.23; **Supplementary Fig. 6C**). The results of these experiments confirmed our previous findings, showing that the relative expression of GLAST and GLT-1 on the plasma membrane does not change between the L- and D-phases of the circadian cycle. Therefore, the changes in the time course of glutamate clearance from astrocytes between the Land D-phase are not due to changes in the total amount of glutamate transporters expressed on the plasma membrane of hippocampal astrocytes.

**Figure 5.**
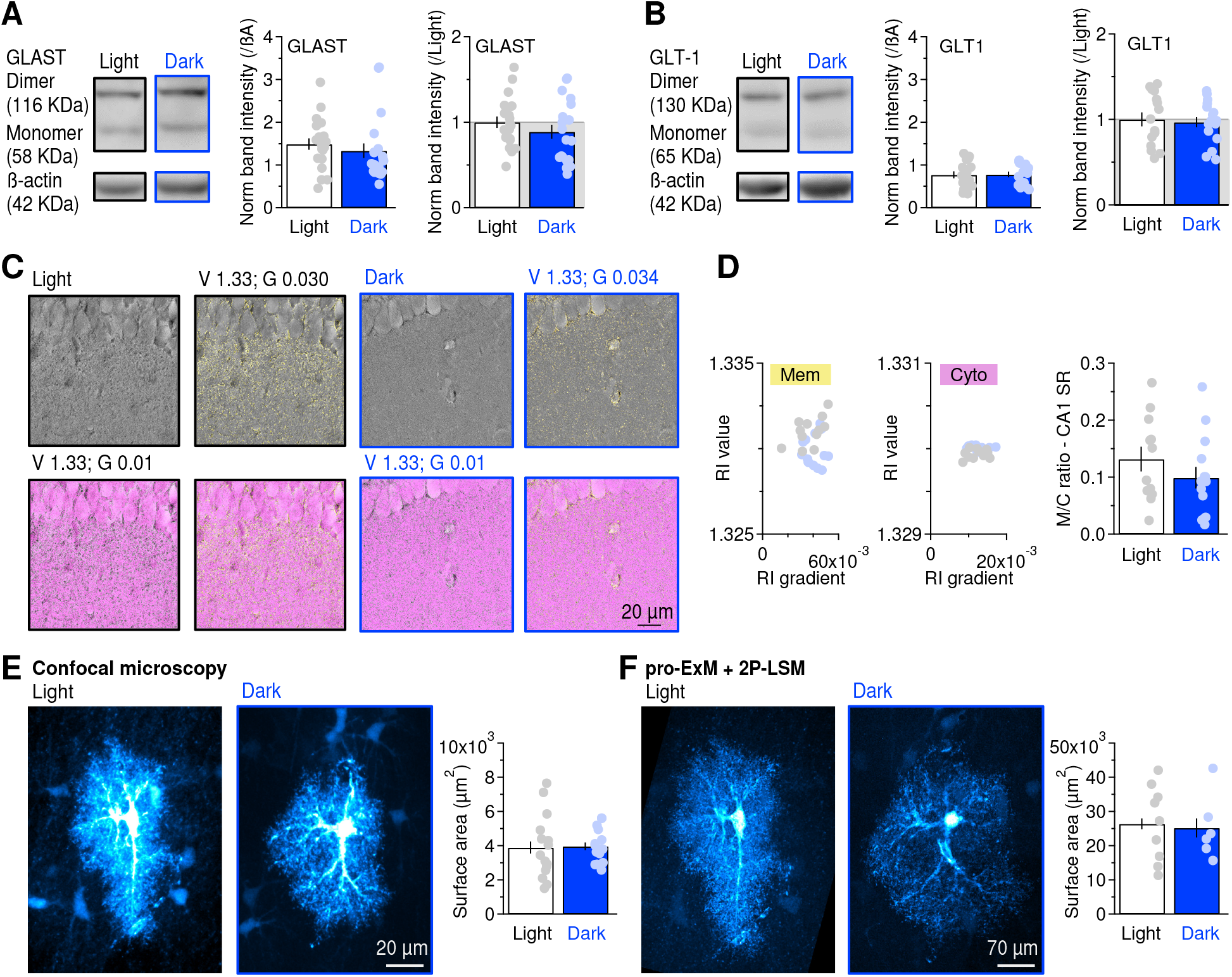
Glutamate transporter expression and the overall astrocyte neuropil coverage are not altered during the L/D-phases. **(A)** Western blot analysis of hippocampal GLAST expression. The two upper bands correspond to GLAST monomers and dimers (*left*). The summary graphs show the quantification of GLAST band intensity, normalized to the corresponding intensity of β-actin (*middle*), or divided by the normalized band intensity in the L-phase (*right*). **(B)** As in (A), for GLT-1. **(C)** Example images collected from slices prepared during the L/D-phases. Each image was digitally stained using refractive index (RI) value (V) and gradient (G) combinations that allowed labeling cell membranes (yellow) and cell cytoplasm (magenta). **(D)** The scatter plots show the specific combinations of RI value and gradient used to label cell membranes (*left*) and cytoplasm (*middle*) across slices. The bar graph (*right*) shows the membrane to cytoplasm (M/C) ratio measured in *stratum radiatum*, with similar M/C ratio values measured during the L/D-phases. **(E)** Maximum intensity projection of biocytin-filled astrocytes collected using confocal microscopy. The summary graph of astrocyte surface area shows no difference between the astrocyte coverage in slices prepared during the L/D-phases. **(F)** Maximum intensity projection of the same astrocytes after proExM collected using 2P-LSM. The summary graph of astrocyte surface area shows no difference between the astrocyte coverage in slices prepared during the L/D-phases.

### The gross structure of the hippocampal neuropil does not change during the circadian cycle

Other potential mechanisms that could account for the slower glutamate clearance during the D-phase are widespread changes in the structure of the neuropil caused by cell swelling or shrinkage. Alternatively, there could be more subtle, local rearrangements of the synaptic environment (Sweeney et al., 2017). We analyzed the overall cell membrane and cytoplasm composition of hippocampal slices prepared at ZT3.5 and ZT15.5 using a marker-free holo-tomographic approach that distinguishes plasma membranes from cytoplasmic compartments based on differences in their refractive-index gradient (Cotte et al., 2013). This approach can detect widespread changes in the plasma membrane/cytoplasm (M/C) ratio caused by storing hippocampal slices for one hour in saline solutions of different osmolality (**Supplementary Fig. 7**). After this incubation time, we fixed the slices, mounted them on slides and collected z-stacks using the 3D Cell Explorer digital holo-tomographic microscope (Nanolive, Ecublens, Switzerland). Plasma membranes and cytoplasmic compartments were detected and digitally stained using the refractive index values and gradients shown in **Supplementary Fig. 7B**. The M/C ratio, calculated as the ratio between the number of voxels that labelled plasma membranes (yellow) and cytoplasm (magenta), provides a readout of the surface area-to-volume ratio of cell processes in hippocampal area CA1. As expected, the M/C ratio increased with increasing osmolality and induced cell shrinkage (**Supplementary Fig. 7C,D**). There was a tight correlation between changes in the M/C ratio and changes in osmolality in both the pyramidal cell layer (**Supplementary Fig. 7C**) and *stratum radiatum*, where the M/C ratio was larger due to the absence of cell bodies (**Supplementary Fig. 7D**). Having confirmed the sensitivity of this imaging technique, we used it to measure the M/C ratio in slices prepared during the L/D-phases. The parameters used to label the plasma membrane and cytoplasm during the L/D-phases were similar among each other and with respect to those used in the osmolality experiments (**Fig. 5C,D**). We used them to measure the M/C ratio in *stratum radiatum*, the domain of area CA1 that contains most astrocytes and excitatory synapses (**Fig. 5D**). The lack of difference between the measures obtained during the L/D-phases suggests that no major cell swelling/shrinking occurs in *stratum radiatum* during the L/D-phases. (M/C ratio_L_: 0.13±0.02 (n=12), M/C ratio_D_: 0.10±0.02 (n=14), p=0.27).

Though valuable, this analysis cannot identify cell-specific structural changes that might occur only in astrocytes. To test whether changes in the overall structure of astrocytes occur during the L/D-phases, we acquired confocal images of biocytin-filled astrocytes and analyzed their maximal intensity projections before and after protein-retention expansion microscopy (proExM) (Tillberg et al., 2016). Using proExM, we can isotropically expand biocytin-filled astrocytes and overcome a caveat of confocal microscopy applied to astrocyte imaging: the spatial resolution of confocal microscopy can be larger than the size of astrocytes’ distal processes (res_x,y_=λ/2·NA=488/2·1.4=174 nm). We subjected hippocampal slices collected at ZT3.5 and ZT15.5 to three, 15 min rounds of expansion (**Supplementary Fig. 8A**), which lead to a roughly 3.5-fold increase in slice area (norm area_L_: 3.4±0.3 (n=6), norm area_D_: 3.5±0.1 (n=6), p=0.84; **Supplementary Fig. 8B-D**) and a 2-fold increase in slice perimeter (norm perimeter_L_: 1.8±0.1 (n=6), norm perimeter_D_: 1.9±0.1 (n=6), p=0.57; **Supplementary Fig. 8E-G**). Before expansion, the biocytin-filled astrocytes imaged by confocal microscopy showed similar surface area during the L/D-phases (area_L_: 3,891±478 μm^2^ (n=15), area_D_: 3,961±224 μm^2^ (n=17), p=0.90; **Fig. 5E,F; Supplementary Fig. 9A**). We imaged astrocytes processed using proExM using two-photon laser scanning microscopy. Even in this case, we found no difference in the average surface area of astrocytes during the L/D-phases (area_L_: 26,400±3,680 μm^2^ (n=9), areaD: 25,200±3,888 μm^2^ (n=6), p=0.83; **Fig. 5E,F; Supplementary Fig. 9B**), suggesting that the gross morphology of these cells does not change during the circadian cycle.

### Astrocytes reduce the number of their fine processes during the dark phase of the circadian cycle

The data outlined above do not rule out the occurrence of more subtle changes in astrocyte morphology, for example in the density of its smallest branches and/or in their proximity to excitatory synapses, which can be resolved using 3D axial STEM tomography (Sousa et al., 2011; Sweeney et al., 2017). We identified astrocytic processes in 5 μm wide, 1-1.5 μm thick tissue samples because of their lack of spines and clear cytoplasm, which distinguishes them from neurons. There were fewer astrocytic processes during the D-phase (2 μm^−3^) than during the L-phase (4 μm^−3^; **Fig. 6A,B**). Their volume and surface area did not change during the L/D-phases (vol_L_: 0.019±0.001 μm^3^ (n=73), vol_D_: 0.023±0.002 μm^3^ (n=37), p=0.11; area_L_: 0.78±0.04 μm^2^ (n=73), area_D_: 0.89±0.05 μm^2^ (n=37) p=0.080; **Fig. 6C,D**). Because of this decrease in the number of astrocytic processes, the nearest neighbor distance between each post-synaptic density and their closest astrocyte process increased more than 2-fold from 155±18 nm (n=59) to 342±63 nm (n=28) during the L- and D-phases, respectively (**p=7.6e-3; **Fig. 6E**). These changes were not associated with changes in the size of the PSD regions (PSD area_L_: 0.024±0.002 μm^2^ (n=59), PSD area_D_: 0.025±0.004 μm^2^ (n=28), p=0.93; **Fig. 6F**).

**Figure 6.**
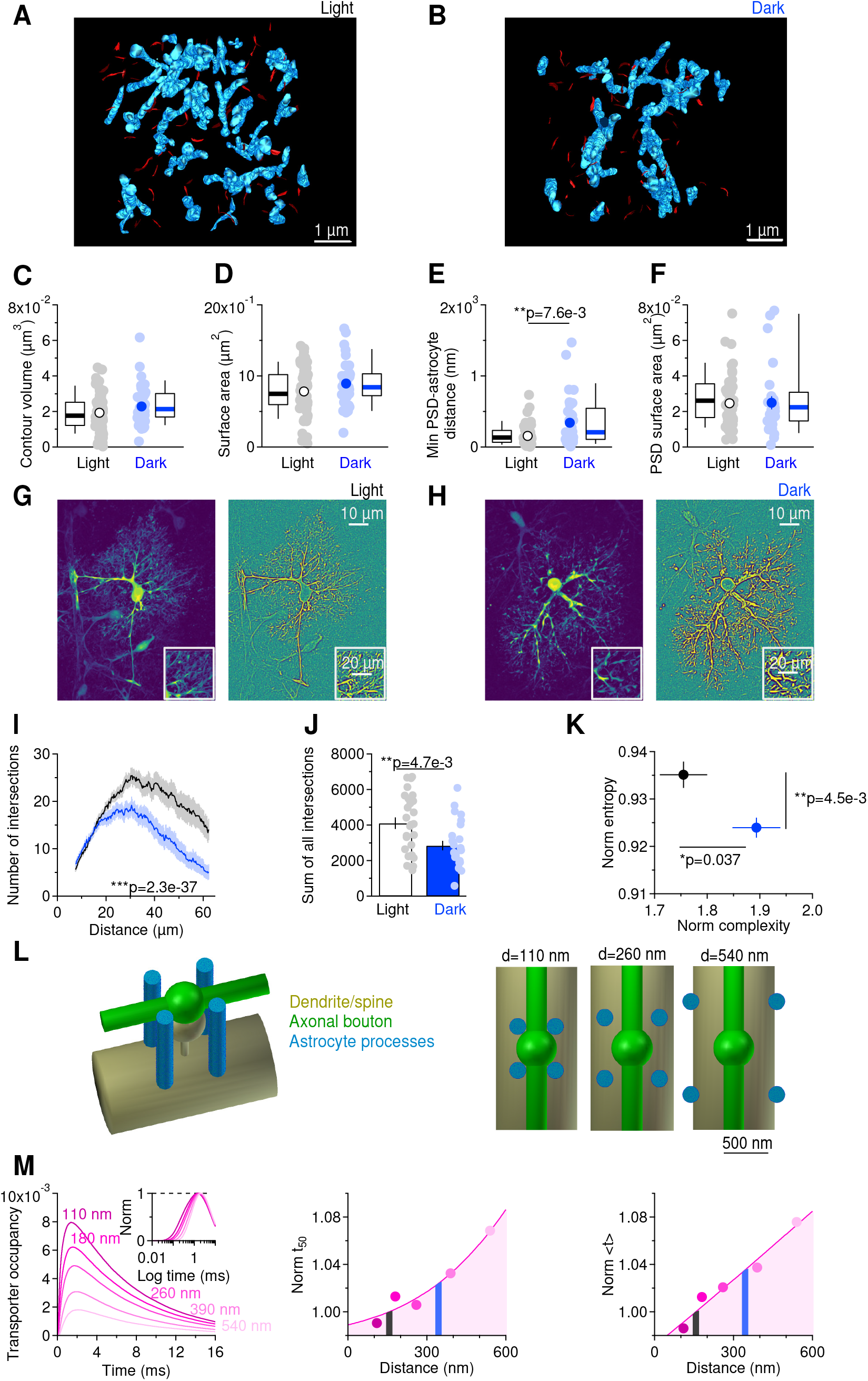
Astrocytes reduce the number of their fine processes during the D-phase, leading to slower glutamate clearance. **(A)** Axial STEM tomography reconstruction of hippocampal processes (blue) and PSDs (red) in *stratum radiatum* of a hippocampal slice prepared during the L-phase. **(B)** As in (A), for the D-phase. **(C)** Astrocytic processes have similar volume in the L/D-phases. The box and whisker plots show the median (thick central line), 25^th^-75^th^ percentile (box top and bottom) and 10^th^-90^th^ percentile (whisker top and bottom). **(D)** Astrocytic processes have similar surface area in the L/D-phases. **(E)** The distance between PSDs and their nearest neighboring astrocyte was increased during the D-phase. **(F)** The PSD surface area did not change during the L/D-phases. **(G)** *Left*, maximum intensity projection of preprocessed confocal z-stacks. *Right*, segmentation of the projection for Sholl analysis with probabilistic thresholding using anisotropic filters. The probabilistic thresholding procedure relied on the following steps. *First*, the input image was mixed with additive Gaussian noise then convolved by a filter bank of isotropic and anisotropic high-pass filters. *Second*, responses to isotropic and highest-amplitude responses to anisotropic filters were thresholded. This procedure was repeated 20 times, and the probabilities of the pixel to be “on” in filter responses are shown as color maps. **(H)** As in G, for biocytin-filled neurons in slices prepared at ZT15.5 (D-phase). **(I)** The Sholl analysis of astrocytes showed that astrocytes have fewer distal processes during the D-phase. **(J)** Summary of the total number of intersections measured by the Sholl analysis using concentric circles centered at the soma, with radii that increased progressively in 1 μm steps. The total number of intersections was reduced during the D-phase. **(K)** Entropy and complexity values for each cell, normalized by the corresponding entropy values in the neuropil around them. The normalized entropy was smaller during the D-phase. In contrast, the normalized complexity was increased during the D-phase. **(L)** *Left*, 3D representation of the synaptic environment, including an axonal bouton (green), a spine with its parent dendrite (gold) and four adjacent astrocytic processes (blue). *Right*, Top view of the synaptic environment with astrocyte processes positioned at increasing distance from the post-synaptic density. **(M)** *Left*, Glutamate transporter occupancy following the release of 2,000 glutamate molecules from the synaptic cleft. The inset shows the peak normalized traces. *Middle*, Normalized half decay time of glutamate transporter occupancy obtained at varying astrocyte-PSD distances. The black and blue lines highlight the values of astrocyte-PSD distance measured in the 3D axial STEM tomography analysis (see panel E). *Right*, As in the middle panel, for centroid measures.

A common way to analyze the number of dendritic branches formed by neurons relies on the use of Sholl analysis. This analysis is prone to errors when used on non-processed confocal images of astrocytes, because the multitude of fine processes of these cells confers them a fuzzy look. Recent image analysis approaches allow information about these fine, filamentous processes to be extracted using coherence-enhancing diffusion filtering (Weickert and Scharr, 2002) and an iterative convolution with oriented filters (see Methods). This procedure allowed a detailed visualization of astrocytic processes in confocal images (**Fig. 6G,H**). The Sholl analysis, applied to the processed confocal images, showed that the number of astrocytic intersections with circles of increasing radii centered on the soma decreased during the D-phase (intersections_L_: 4,108±326 (n=26), intersections_D_: 2,844±272 (n=20), **p=4.7e-3; **Fig. 6I,J**). An alternative method, not requiring image binarization, relies on the use of a shearlet multiscale framework (Guo and Labate, 2007) and provides an optimal approximation of anisotropic features like astrocytic process in confocal images (Brazhe, 2018). This method uses spatial entropy (H) and complexity (C) measures to describe the orderliness and feature-preferred orientation of an image, respectively (Brazhe, 2018; Gavrilov et al., 2018). In our analysis, we normalized the values of spatial entropy and complexity measured in the area covered by the biocytin-filled neuron by the values obtained in the adjacent neuropil. Consistent with the Sholl analysis, the normalized entropy was smaller during the D-phase (L: 0.935±0.003 (n=29), D: 0.924±0.002 (n=16), *p=0.037; **Fig. 6K**). These results support the 3D axial STEM data and indicate that astrocytes remodel their processes during the circadian cycle. To further test whether changes in the location of astrocytic processes with respect to excitatory synapses could lead to changes in the time course of astrocyte transporter currents, we developed a 3D representation of the synaptic environment and used it to run 3D Monte Carlo reaction-diffusion simulations (see Methods). We performed different sets of simulations, in which we varied the astrocyte-PSD distance from 110 nm to 540 nm. The results show that changing the average distance of the astrocytic processes, within the range detected in the 3D axial STEM tomography reconstructions in samples collected during the L- and D-phases, leads to a prolongation of the decay time of the glutamate transporter occupancy (**Fig. 6L,M**), which in turn can slow down the time course of STCs and of glutamate clearance from the extracellular space.

### Neurons and astrocytes cooperate to mediate circadian changes in excitatory synaptic transmission

The experiments describe so far show that the structural and functional properties of hippocampal astrocytes change over the course of the circadian cycle, in ways that might affect LTP expression. What is not clear is whether this effect can, on its own, account for the loss of LTP in the D-phase. To test this hypothesis, we first performed titration experiments with the positive allosteric modulator of AMPA receptors LY404187 on AMPA EPSCs (**Fig. 7A**) (Quirk and Nisenbaum, 2002). A maximal potentiation of amplitude and decay of single AMPA EPSCs was obtained with LY404187 concentrations greater than 0.02 μM (**Fig. 7A**, *top*). This concentration of LY404187 led to reduced AMPA EPSC summation in slices prepared during the L-phase, similar to the one detected in the D-phase (**Fig. 2A**). Is this pharmacological manipulation capable, on its own, to occlude LTP in the L-phase? To answer this question, we applied the TBS LTP-induction protocol in the presence of LY404187 (0.02 μM), in slices prepared during the L-phase. However, we found that LY404187 did not alter LTP (1.14±0.02 (n=9), ***p=3.7e-5; **Fig. 7B**, *top*). This suggests that the prolonged activation of AMPA receptors caused by astrocyte retraction from synapses and the reduced temporal summation of AMPA EPSCs are not the sole mechanisms to account for loss of LTP during the D-phase. There is, however, the possibility that together, the reduced NMDA receptor expression and altered time course of AMPA receptor activation might degrade LTP at Schaffer collaterals in the D-phase. We tested this hypothesis by applying the TBS LTP-induction protocol in the presence of both APV (2 μM) and LY404187 (0.02 μM), in slices prepared during the L-phase (**Fig. 7B**, *bottom*). This manipulation led to a complete occlusion of LTP in slices prepared during the L-phase (0.98±0.03 (n=8), p=0.61).

**Figure 7.**
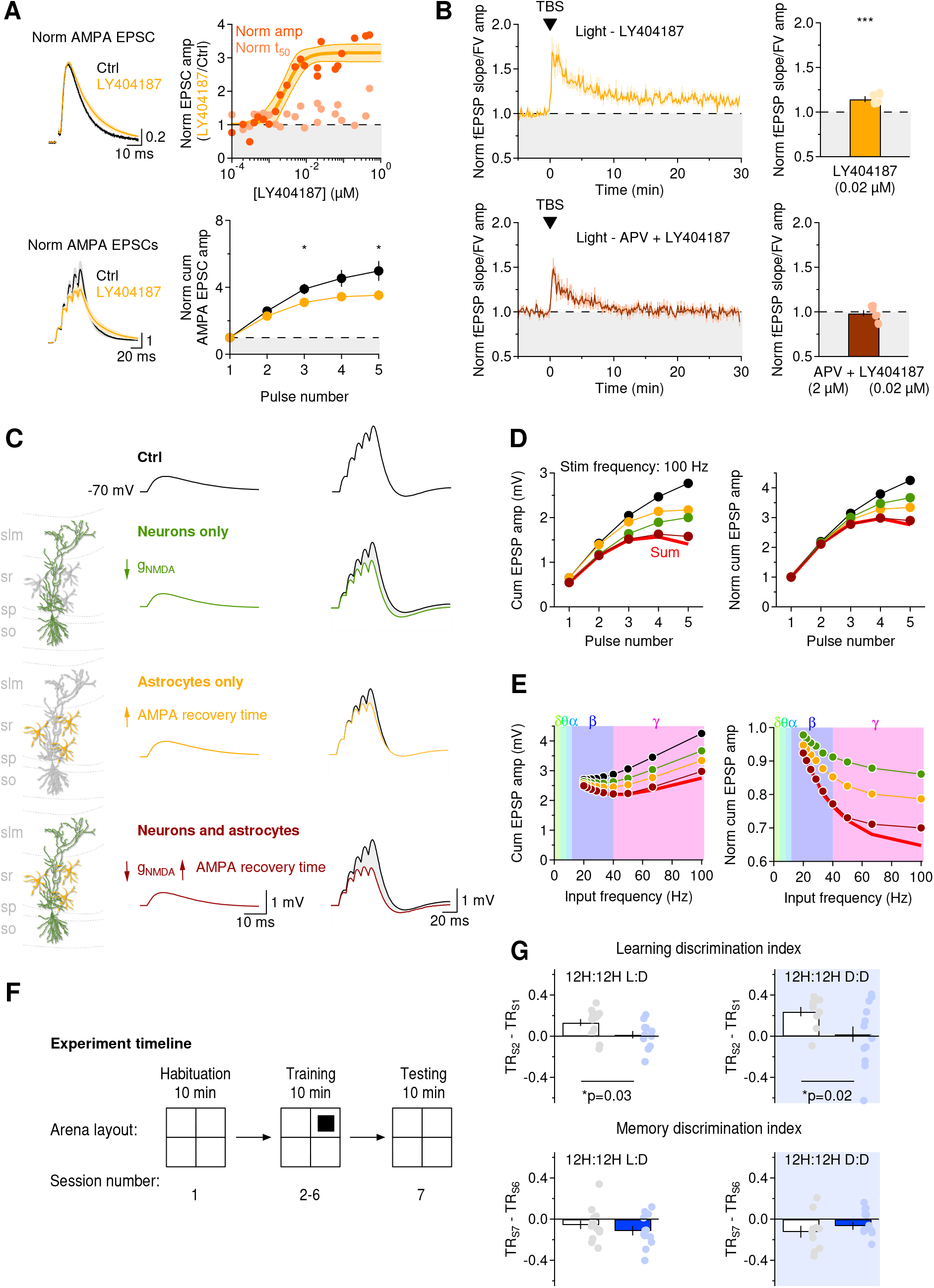
Astrocytes and neurons cooperate to mediate circadian changes in synaptic integration over specific frequency ranges. **(A)** *Top left*, Peak normalized AMPA EPSCs in control conditions and in the presence of LY404187 (0.02 μM). Each trace represents the average of recordings collected in n=15 cells. *Top right*, Titration curve showing the effect of increasing concentrations of LY404187 on the amplitude (red dots) and half decay time (t_50_, yellow dots) of AMPA EPSCs recorded in slices prepared in the L-phase. The thick orange line and shaded areas represent the Hill fit of the amplitude data and 95% confidence bands, respectively. *Bottom left*, Train of AMPA EPSCs recorded in voltage clamp mode while holding pyramidal cells at a holding potential of −65 mV. Each train stimulation was composed of five pulses, with an inter-pulse interval of 10 ms. *Bottom right*, The AMPA EPSC summation was reduced in the presence of LY404187 (0.02 μM). **(B)** *Top left*, Time course of baseline-subtracted ratio between the field EPSC slope and the fiber volley amplitude in slices prepared at ZT3.5 (light), in the presence of LY404187 (0.02 μM). *Top right*, Statistical summary, showing that LTP in the L-phase is not occluded by LY404187 (0.02 μM). *Bottom left*, Time course of baseline-subtracted ratio between the field EPSC slope and the fiber volley amplitude in slices prepared at ZT3.5 (light), in the presence of LY404187 (0.02 μM) and APV (2 μM). *Top right*, Statistical summary, showing that LTP in the L-phase is occluded by the concurrent application of LY404187 (0.02 μM) and APV (2 μM). **(C)** NEURON simulations of composite AMPA an NMDA EPSPs evoked using the realistic model of a CA1 pyramidal cell shown in Fig. 4D. The simulation was run to mimic glutamatergic EPSPs during the L-phase (*black*) and in conditions that mimic the reduced NMDA receptor expression during the D-phase (*green*), the reduced AMPA EPSP summation during the D-phase due to retraction of astrocytic processes (*yellow*) or both effects at the same time (*brown*). The simulations were run using either single pulses or five pulses delivered at varying frequencies. **(D)** *Left*, Cumulative EPSP amplitude recorded when delivering five pulses at 100 Hz. The red line represents the arithmetic sum of the effect introduced by reducing the NMDA receptor conductance (*green*) and by increasing the recovery time of AMPA receptors (*yellow*). Simulations run when combining these effects (*brown*) show that the combined effect of the circadian changes on AMPA/NMDA EPSP summation is sublinear (i.e. smaller than the arithmetic sum shown in red). *Right*, As in (A), after normalizing the data by the peak of the first EPSP. **(E)** Results of NEURON simulations obtained in response to stimulations with five pulses at different frequencies. The graph on the left hand side represents the cumulative glutamatergic EPSP amplitude. The graph right hand side shows the cumulative EPSP amplitude, obtained by normalizing the peak depolarization after the fifth stimulus by the amplitude of the first EPSP. The results show that the combined effect of the detected circadian changes in neuron and astrocyte function are sub-linear for stimulation frequencies greater than 50 Hz. **(F)** Schematic of the experimental timeline for the novel object recognition behavioral test. **(G)** Summary results for the learning and memory discrimination index measured in mice maintained on 12H:12H L:D (*left*) and D:D cycles (*right*). The mice were trained and tested either in the L- or D-phase (*left*), or in the subjective L- and D-phase (*right*). The proportion of time spent in the top right corner of the arena, where the novel object is positioned, is abbreviated as TR.

### Novel object recognition, not memory of familiar environments, is susceptible to circadian modulation

We used the realistic biophysical model of a CA1 pyramidal neuron described in **Fig. 3** to determine how decreased NMDA receptor expression and increased recovery from activation of AMPA receptors alter the summation of composite glutamatergic EPSPs at different stimulation frequencies (**Fig. 7C-E**). We simultaneously activated 20 excitatory synapses randomly distributed in the apical dendrites between 50 and 300 μm from the soma using a train of 5 pulses delivered at 10-100 Hz. The results show that changes in NMDA receptor activation or AMPA receptor recovery time reduce EPSP summation at β-γ frequencies. A further reduction in temporal summation was observed when both manipulations were performed at the same time. Their effects summate linearly in the β and slow γ frequency range (<50 Hz), and sub-linearly for inputs in the fast (65-100 Hz) γ range. These results are relevant in the context of hippocampal-dependent cognition and exploratory behaviors.

In hippocampal area CA1, slow γ oscillations synchronize with slow γ activity in area CA3 during exploration of familiar environments and memory recall (Colgin et al., 2009). By contrast, fast γ activity synchronizes with fast γ activity in entorhinal cortex during novel object recognition and learning (Zheng et al., 2016), an effect that is mediated by activation of ionic conductances non-uniformly distributed in the proximal and apical dendrites of CA1 pyramidal cells (Combe et al., 2018). The results of this model suggest that the circadian modulation of the molecular composition of excitatory synapses and of astrocyte morphology exert a profound modulation of hippocampal activity in the high γ range, whereas integration of inputs at lower frequency ranges are less affected. If this were true, we would expect behaviors that rely on high frequency hippocampal activity, for example learning ad exploration of novel environments, to vary with the time-of-day. By contrast, memory recall and exploration of familiar environments, which rely on hippocampal activity at lower frequencies, should be less susceptible to circadian modulation. We tested this hypothesis *in vivo*, using a novel object recognition behavioral test (**Fig. 7F,G**) (Antunes and Biala, 2012; Broadbent et al., 2010; Ennaceur and Delacour, 1988). We used four cohorts of mice for this test: two cohorts were kept under 12H:12H L:D conditions and were trained and tested either in the L-phase or in the D-phase. Two other cohorts of mice were kept in constant darkness and were trained and tested either in the subjective L-phase or in the subjective D-phase. During the first habituation session, we positioned a mouse in an open field arena and allowed it to freely explore it for 10 min. During the following five training sessions, we re-introduced the mouse in the same arena, after positioning a non-familiar object in its top right corner, which we labelled with a visual cue. Then, on the last session (i.e. session 7), we removed the object and allowed the mouse to explore the open field arena (**Fig. 7F**). We quantified object exploration by computing the learning discrimination index from scores of the proportion of time that each mouse spent exploring the new object in the first training session (**Fig7G**, *top*). Object memory was quantified by computing the discrimination index from scores of the proportion of time that the mice spent in the corner of the arena that used to contain the novel object during the last testing session (**Fig. 7G**, *bottom*). The results showed a significant decrease in the learning discrimination index in mice trained and tested during the D-phase (LDI_L_: 0.13±0.03 (n=14), LDI_D_: 0.02±0.04 (n=12), *p=0.03). By contrast, the memory discrimination index remained similar between mice trained and tested either in the L- or D-phase (MDI_L_: −0.05±0.04 (n=14), MDI_D_: −0.11±0.04 (n=12), p=0.34; **Fig. 7G**, *left*). Similar results were obtained using mice kept in constant darkness (**Fig. 7G**, *right*). Here, the learning discrimination index was reduced during the subjective D-phase (LDI_SL_: 0.24±0.05 (n=10), LDI_SD_: 0.02±0.08 (n=16), *p=0.02), whereas the memory discrimination index did not change between mice trained and tested during the subjective L- and D-phase (MDI_SL_: −0.12±0.06 (n=10), MDI_SD_: −0.07±0.04 (n=16), p=0.42; **Fig. 7G**, *right*). The consistence of the results collected from mice kept under L:D and D:D conditions indicates that they are due to circadian modulation. The main finding here is that, learning – not memory recall – shows the highest sensitivity to time-of-day.

### The role of corticosterone

As a last step in our analysis, we aimed to gain insights into the mechanisms that could account for the circadian modulation of the receptor composition and activation at Schaffer collateral synapses. We reasoned that the circadian changes in LTP might be triggered by an increase in the circulating levels of corticosterone, a glucocorticoid hormone whose production is known to increase during the active phase in adult mice (i.e. the D-phase). By using radioimmunoassay, we confirmed that changes in the blood levels of corticosterone occur in juvenile mice aged P14-21 (**Fig. 8A**, *left*). A similar trend was detected in mice kept in constant darkness, where the plasma levels of corticosterone peaked at the end of the subjective D-phase (**Fig. 8A**, *right*). If the circadian changes in LTP were due to higher activation of corticosteroid receptors by increased circulating corticosterone levels in the D-phase, blocking these receptors pharmacologically should rescue LTP during the D-phase without altering LTP in the L-phase. Consistent with our hypothesis, blocking NR3C1 mineralocorticoid receptors with spironolactone (10 μM; LTP_L_: 1.19±0.03 (n=7), **p=1.4e-3) and blocking NR3C2 glucocorticoid receptors with mifepristone (1 μM; LTP_L_: 1.17±0.02 (n=8), ***p=1.9e-4) did not occlude LTP in slices prepared during the L-phase (**Fig. 8B,C**). By contrast, blocking NR3C1 (LTP_D_: 1.13±0.02 (n=8), ***p=5.9e-4) or NR3C2 receptors (1.23±0.04 (n=8), ***p=9.1e-4) in slices prepared during the D-phase rescued LTP (**Fig. 8D,E**). We tested whether these effects could be mimicked by blocking melatonin receptors, a hormone produced by the pineal gland during the D-phase (Korf and von Gall, 2006). C57BL/6 mice carry two mutations in biosynthetic enzymes for melatonin (Ebihara et al., 1986; Kasahara et al., 2010; Peirson et al., 2018) and MT1/2 melatonin receptor gene expression is low in the hippocampus (Saunders et al., 2018). Consistent with these findings, blocking MT1/2 melatonin receptors with luzindole (50 μM) in slices prepared during the L-phase did not occlude LTP (LTP_L_: 1.16±0.04 (n=11), **p=1.4e-3; **Fig. 8B,C**). No LTP was detected in the D-phase in the presence of luzindole (LTP_D_: 1.05±0.04 (n=7), p=0.22; **Fig. 8D,E**). Is the rescue of LTP detected by blocking NR3C1/2 receptors in the D-phase due to a rescue of NMDA receptor expression and AMPA receptor activation? We tested this by repeating the RuBi-glutamate uncaging and the train stimulation experiments. We found that the NMDA/AMPA ratio of EPSCs evoked by flash uncaging RuBi-glutamate (100 μM) was larger in slices treated with spironolactone (5.3±1.0 (n=5), **p=4.4.e-3) or mifepristone (5.0±0.9 (n=5), **p=9.3e-3), not luzindole (1.4±0.5 (n=5), p=0.69; **Fig. 8F**). The temporal summation of AMPA EPSCs in slices treated with NR3C1/2, not MT1/2 antagonists, was similar to that recorded in slices prepared during the L-phase (**Fig. 8G**). These data identify corticosterone and corticosteroid receptors as key players in the circadian modulation of synaptic strength at Schaffer collateral synapses.

**Figure 8.**
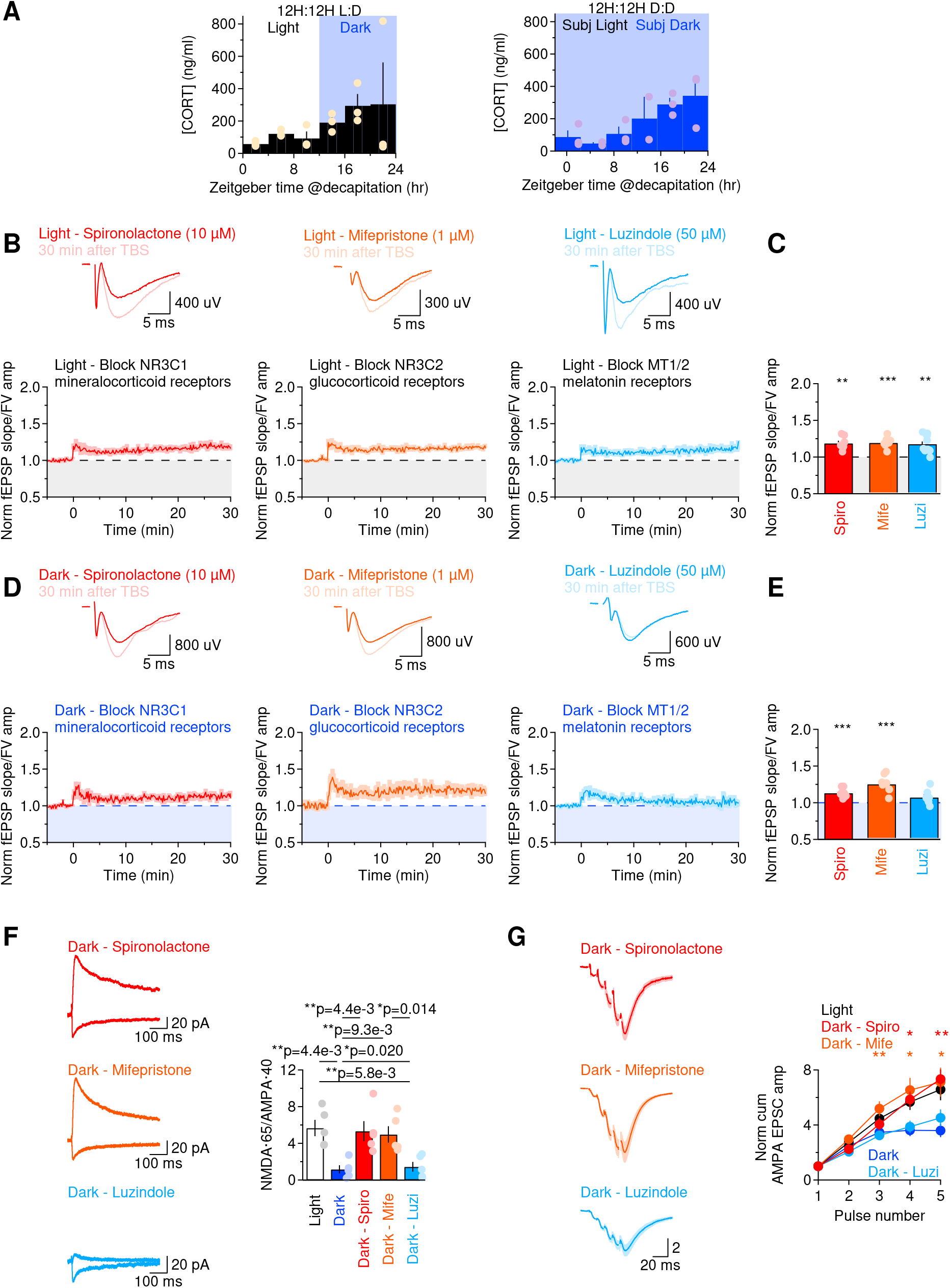
Blocking corticosteroid receptors rescues LTP, NMDA receptor expression and AMPA receptor activation in the dark phase. **(A)** Plasma corticosterone measured at baseline, showing a progressive increase during the D-phase, which in mice represents the active phase (*left*). A progressive increase in baseline levels of plasma corticosterone is also detected during the subjective D-phase in mice kept on a 12H:12H D:D schedule (*right*). **(B)** *Top*, average responses recorded 5 min before and 30 min after TBS. *Bottom*, time course of baseline-subtracted ratio between the field EPSC slope and the fiber volley amplitude in slices prepared at ZT3.5 (light), in the presence of the NR3C1 mineralocorticoid receptor antagonist spironolactone (10 μM), the NR3C2 glucocorticoid receptor antagonist mifepristone (1 μM) and the melatonin receptor antagonist luzindole (50 μM). **(C)** Statistical summary, showing that LTP in the L-phase was not affected by NR3C1/2 and MT1/2 antagonists. **(D)** As in (A), for slices prepared at ZT15.5 (dark). **(E)** Statistical summary, showing that LTP in the D-phase can be rescued by blocking NR3C1/2 receptors. **(F)** Average flash-evoked AMPA NMDA EPSCs recorded from CA1 pyramidal cells in response to RuBi-glutamate uncaging (100 μM). The slices were pre-incubated for >30 min with spironolactone (10 μM), mifepristone (1 μM), or luzindole (50 μM). The amplitude of flash-evoked NMDA EPSCs is rescued by blocking NR3C1/2, not MT1/2, receptors. **(G)** *Left*, train of AMPA EPSCs recorded in voltage clamp mode while holding pyramidal cells at a holding potential of −65 mV. Each train stimulation was composed of five pulses, with an inter-pulse interval of 10 ms. The AMPA EPSC summation in the D-phase was rescued by blocking NR3C1/2, not MT1/2, receptors. *Right*, Summary graph.

## Discussion

The results presented in this manuscript provide direct evidence for the existence of a circadian modulation of excitatory synaptic transmission within the juvenile hippocampus (**Fig. 1**), due to structural and functional remodeling of the synaptic environment. Neurons and astrocytes both contribute to this phenomenon, though in different ways. Neurons alter the size of the functional pool of glutamate NMDA receptors at the synapse (**Fig. 2**). Astrocytes change the number and proximity of their fine processes with respect to excitatory synapses (**Fig. 6**). The remodeling of astrocytes alters the time course of glutamate clearance from the extracellular space (**Fig. 4**). Although this does not have notable consequences on glutamate receptor activation during sparse synaptic stimulation, it profoundly alters the temporal summation of recurring stimuli, long-term plasticity and hippocampal-dependent learning (**Fig. 7**). We identify corticosterone and corticosteroid receptors as candidate triggers for these effects (**Fig. 8**).

### Astrocytes, neurons and the values of an ever-changing brain

Modifications of the synaptic environment are critical for information coding in the brain, and provide an important substrate for circuit plasticity. The shape, size and turnover of dendritic spines can be controlled in an activity-dependent manner over time scales that range from seconds to hours, during development (Holtmaat et al., 2005; Kwon and Sabatini, 2011; Rakic et al., 1986; Richards et al., 2005), in response to LTP-inducing protocols (Engert and Bonhoeffer, 1999) and in the adult brain (Gilbert, 1998). Analogous activity-dependent phenomena of structural plasticity also occur in small protoplasmic astrocytic processes that are commonly found around synapses (Lavialle et al., 2011; Reichenbach et al., 2010). Accordingly, different groups have reported that these fine processes can extend, retract, bud and glide through the hippocampal and brainstem neuropil (Haber et al., 2006; Hirrlinger et al., 2004). In the SCN, similar phenomena occur with circadian rhythmicity (Becquet et al., 2008). The distance over which these movements take place are comparable to the distance that spines can grow or shrink (i.e. hundreds of nm) and are therefore physiologically relevant. One of the reasons these movements are functionally important is that small changes in the relative proximity of astrocytic processes and spines have important consequences on the lifetime of glutamate in the extracellular space, its uptake, and ultimately for the regulation of the strength and timing of information transfer across neurons. Accordingly, suppressing astrocyte motility has been shown to impair the stabilization and maturation of synapses (Nishida and Okabe, 2007).

At excitatory glutamatergic synapses, multiple molecular mechanisms can lead to changes in the proximity of astrocytic processes to spines, which in turn contributes to modify the time course of glutamate receptor activation. For example, we previously showed that activating PAR1 G-protein coupled receptors leads to proliferation and closer apposition of astrocytic processes to excitatory synapses, leading to reduced AMPA and NMDA receptor activation (Sweeney et al., 2017). Since PAR1 receptors are typically activated by serine proteases in the bloodstream, we concluded that this movement of astrocytes towards synapses might be indicative of similar phenomena of structural plasticity of the synaptic environment occurring during small hemorrhagic stroke. The experimental data described here indicate that astrocyte remodeling is not a rare phenomenon that only happens in experimental conditions mimicking pathological states, but can also occur daily under physiological conditions in the hippocampus. These recurring, circadian changes in the relative proximity of astrocytes and spines occur in concert with changes in glutamate receptor activation in pyramidal neurons, which in our experiments are not due to concurrent changes in the release of the NMDA receptor co-agonist D-Serine from astrocytes. Together, these phenomena allow the brain to regularly sculpt the fundamental units of its neuronal circuits and provide a new mechanistic understanding of the molecular events that control the circadian modulation of synaptic plasticity in the hippocampus.

We can only speculate on why the brain invests its energies to constantly turn up and down the strength of its synaptic connections. One possibility is that this may be a necessary step in a long-term investment plan, whereby tuning down synaptic efficacy during the D-phase may bring the hippocampus into a low energy-demanding state, to parse energy consumption over longer periods of time (Attwell and Laughlin, 2001). Because of these fluctuations, hippocampal networks may be able to switch between functioning modes that best process different types of information at specific time points during the day. Our data may provide a mechanistic basis for Marr’s theory of the archicortex (Marr, 1971), in which the hippocampus alternates between times during which it stores information about patterns of neural activity representing events as they happen, and times during which this information is transferred to other regions of the brain, including the neocortex (Willshaw et al., 2015). Based on our experiments, for nocturnal animals like mice, the time to store information locally would be the L-phase, whereas the time to relay information to the neocortex would be the active D-phase.

The frequency-dependent effect of these phenomena on synaptic integration also suggests that different types of hippocampal activity exhibit different susceptibility to circadian modulation. Specifically, hippocampal activity in the high γ-range, associated with learning and memory encoding, is highly susceptible to circadian modulation. In contrast, hippocampal activity that encodes episodic, semantic, and working memory information (θ, α, β-ranges) is much less affected by time-of-day. Our behavioral experiments support this hypothesis, showing that learning and memory display different susceptibilities to circadian modulation. According to our findings, time of day affects our ability to learn due to fluctuations in circulating corticosterone levels. By contrast, longer-term memories, hard-wired in the brain, remain relatively stable at different times of day.

### LTP, melatonin and glucocorticoids

The first observation that LTP expression at hippocampal Schaffer collateral synapses varies at different times of the day dates back to 1983, when Harris and Teyler first showed that LTP at Schaffer collateral synapses in juvenile and adult rats was more robust in slices prepared during the L-phase (Harris and Teyler, 1983). Based on the fact that these results were obtained in reduced slice preparations, it was concluded that they did not depend on the activity of extra-hippocampal afferents but rather on the enduring effects of modulators that are lost very slowly from slice preparations, for example hormones. Similar conclusions were later obtained in hamsters (Raghavan et al., 1999), highlighting the need to determine the underlying causes.

One of the first candidate molecules that was thought to mediate these effects was the hormone melatonin, mainly synthesized during the D-phase in the pineal gland from the precursor serotonin (Vanecek, 1998). In acute hippocampal slices, exogenously applied melatonin blocks LTP induction without altering low-frequency synaptic transmission (Collins and Davies, 1997; Wang et al., 2005). These effects are unlikely to be mediated by melatonin binding directly to melatonin receptors, because few receptors are expressed in the hippocampus. Given that melatonin has close structural similarity to various antagonists of the glycine-binding site of NMDA receptors (Huettner, 1989; Salituro et al., 1992; Salituro et al., 1990), it has been suggested that melatonin could reduce LTP by antagonizing NMDA receptors. This hypothesis has been tested and ultimately disputed by Collins and Davies (Collins and Davies, 1997). If melatonin does not have any known membrane receptor target, how does it exert its biological actions? Because of its hydrophobicity, melatonin released from the pineal gland into the bloodstream can easily pass through cell membranes. This allows melatonin to modulate a variety of intracellular molecular pathways and even bind to nuclear receptors (Benitez-King et al., 2001; BenitezKing et al., 1996; Carlberg and Wiesenberg, 1995; Rafii-El-Idrissi et al., 1998). Naturally, some of these interactions could have a deleterious effect on LTP. The problem is that many inbred strains of laboratory mice, like C57BL/6J, have no detectable levels of melatonin in the pineal gland due to two independent mutations in the genes encoding the biosynthetic enzymes for melatonin: N-acetylserotonin and hydroxyindole-o-methyltransferase (Ebihara et al., 1986; Kasahara et al., 2010; Peirson et al., 2018). It is therefore unlikely that our circadian effects on astrocyte morphology and AMPA EPSC summation are mediated by melatonin acting on molecular targets aside from MT1/2 receptors and our results do not support this hypothesis.

Other hormones known to be produced with circadian rhythmicity are adrenal corticosteroids, like corticosterone, which act by activating glucocorticoid and mineralocorticoid receptors (Chung et al., 2011). High-affinity (Type I) mineralocorticoid receptors and low-affinity (Type II) glucocorticoid receptors are abundantly expressed in the hippocampus and both affect synaptic plasticity and memory formation (Kim and Yoon, 1998). Because of their different steady-state affinity, Type I receptors are fully occupied at low glucocorticoid plasma levels, whereas Type II receptors become fully occupied only with higher glucocorticoid plasma levels (Chung et al., 2011; Pavlides et al., 1995). Physiological increase in alertness and arousal during the D-phase of the circadian cycle can activate both types of receptors. Although the exclusive activation of Type I receptors promotes LTP, the concomitant activation of Type II receptors suppresses it (Pavlides et al., 1995). Consistent with these findings, our experiments show that the detected loss of LTP in the D-phase is associated with activation of both Type I and Type II receptors. To the best of our knowledge, the molecular and cellular mechanisms leading Type I-II receptor activation to loss of LTP had not been previously determined.

### Generalizability and functional implications

Mice have evolved as nocturnal animals because they have fewer predators at night, making this the prime time to search for food. This nocturnal pattern of activity is still detected in mice kept in captivity. Although most animal facilities offer *ad libitum* access to food, caretakers are generally not around at night, likely contributing to make this the time of least perceived danger. In both nocturnal and diurnal species, the production of melatonin peaks during the D-phase, due to the fact that light inhibits its production through the activation of retino-hypothalamic afferent projections to the SCN. In contrast, the timing of glucocorticoid production coincides with the active phase, which is inverted between nocturnal and diurnal animals. This is because light exerts different effects on SCN neurons receiving light-sensitive inputs in the SCN of diurnal and nocturnal species. In diurnal species, light reduces the firing rate of photosensitive neurons. By contrast, in nocturnal species, light increases the firing rate of these cells. Given that activation of the SCN inhibits the HPA axis through the release of vasopressin, light stimulates the release of corticosteroids in diurnal but not in nocturnal animals (Jiao et al., 1999; Smale et al., 2003). Therefore, to a first approximation, the production of corticoisteroids in humans and mice is locked to their activity phase (Dickmeis, 2009). Because in our experiments activation of corticosteroid receptors disrupts plasticity at Schaffer collateral synapses, it is tempting to speculate that synaptic plasticity might also be degraded in the human hippocampus through mechanisms similar to those described here but with opposite periodicity. However, since there are numerous molecular and behavioral differences and similarities between humans and mice, one needs to be cautious before jumping to the conclusion that L-phase of humans is analogous to the D-phase of a mouse. That said, our findings indicate that there are circadian changes in the molecular landscape of the hippocampus that enable this structure of the brain to switch to different functional modes. As a substantial reprogramming of oscillating transcripts occurs in temporal lobe epilepsy (Debski et al., 2017), our findings may have important implications to advance not only our understanding of circadian rhythmicity during physiological conditions but also to understand the pathogenesis and treatment of this and other chronic disease states.

## Acknowledgements

We would like to thank Drs. Pablo Valdes and Ed Boyden for support with the proExM protocol, Tyler A. Mitchell for support with biotynilation and behavior experiments, and Dr. Jennifer M. Hurley for her generous gift of Per1/2^-/-^ mice. Thanks to Drs. Juan Burrone and Christophe Bernard for valuable discussions and Dr. Daniela Tropea for comments on the manuscript.

## Author Contributions

Conceptualization: A.Sc.

Methodology: A.B., A.Sc.

Formal Analysis: J.P.M., L.Y.D., N.A., J.J.W., S.Z., S.S., A.A.S, D.G.Z., R.M., A.Sc.

Investigation: J.P.M., M.A.P., L.Y.D., G.C.T., N.A., R.M.D., R.M., A.Sc.

Resources: R.D.L., A.K., A.Se., M.M., A.Sc.

Writing – Original Draft Preparation: A.Sc.

Writing – Review & Editing: A.Sc.

Visualization: A.Sc.

Supervision: A.Sc.

Project Administration: A.Sc.

Funding Acquisition: This work was supported by the Presidential Award for Undergraduate Research and Initiative for Women Award to N.A.; RFBR COMFI (Grant Number 17-00-00412K) for joint research to A.Se. (Grant Number 17-00-00409) and A.B. (Grant Number 17-00-00407); NIH/NIBIB Intramural Research Program (Grant Number ZIAEB000046) to R.D.L.; NIH/NIDA (R01DA04741001) to A.K.; NIH/NIMH (Grant Number R15MH118692), SUNY Albany and SUNY Albany Research Foundation to D.G.Z; EU Horizon 2020 Framework Program for Research And Innovation (Grant Number 785907 and 945539, Human Brain Project SGA2 and SGA3) to M.M. and R.M.; NSF (Grant number IOS2011998) and NIH/NINDS (Grant Number R03NS102822), SUNY Albany and SUNY Albany Research Foundation to A.Sc.

---

**Supplementary Figure 1.**
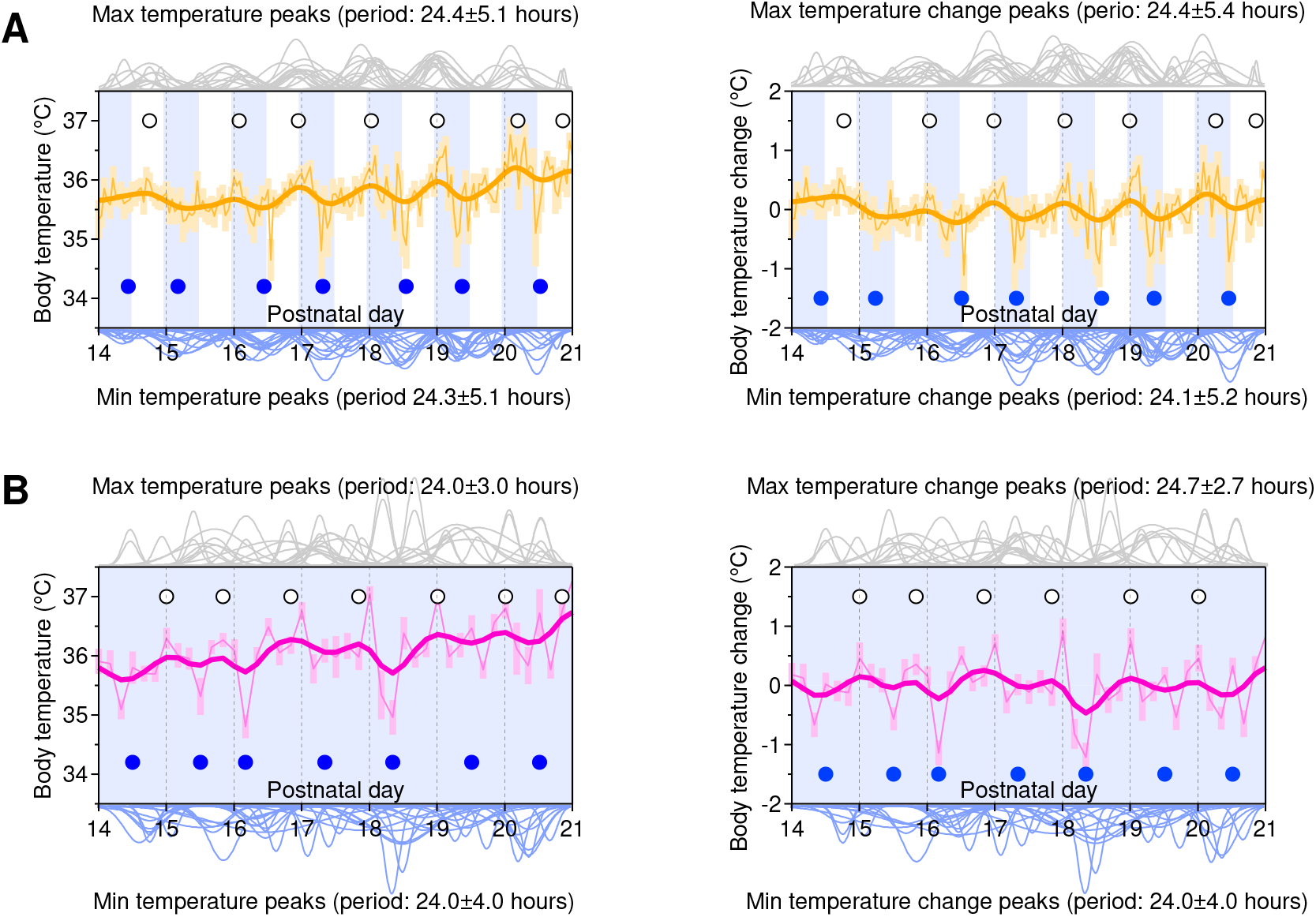
Circadian rhythmicity is present in juvenile mice. **(A)** *Left*, Average hourly body temperature values measured using subcutaneous transponders on P14-21 mice. *Right*, Average changes in hourly body temperature values measured using subcutaneous transponders on P14-21 mice, after subtraction of the mean body temperature of each mouse averaged over the entire duration of the experiment. The white and blue shaded areas in the graph represent the times of light exposure (L-phase, ZT0-12) and darkness (D-phase, ZT12-24), respectively. The white and blue dots represent the position of the maximum and minimum temperature peaks during each 24 hour period, respectively. The thin orange line and yellow shaded areas represent the mean±SEM of the temperature readouts across mice. The thick orange line represents the smoothed mean, obtained using a boxcar smoothing algorithm with 10 repetitions and a 5-point boxcar width. The grey and blue curves at the top and bottom of the figure represent a multi-peak fit of the data collected from each mouse, to determine the position of the maximum (top) and minimum body temperature for each mouse (bottom). Regularly spaced peaks for maximum and minimum body temperature occur with circadian and alternate rhythmicity in P14-21 mice. **(B)** As in (A), for mice kept in 12H:12H D:D conditions.

**Supplementary Figure 2.**
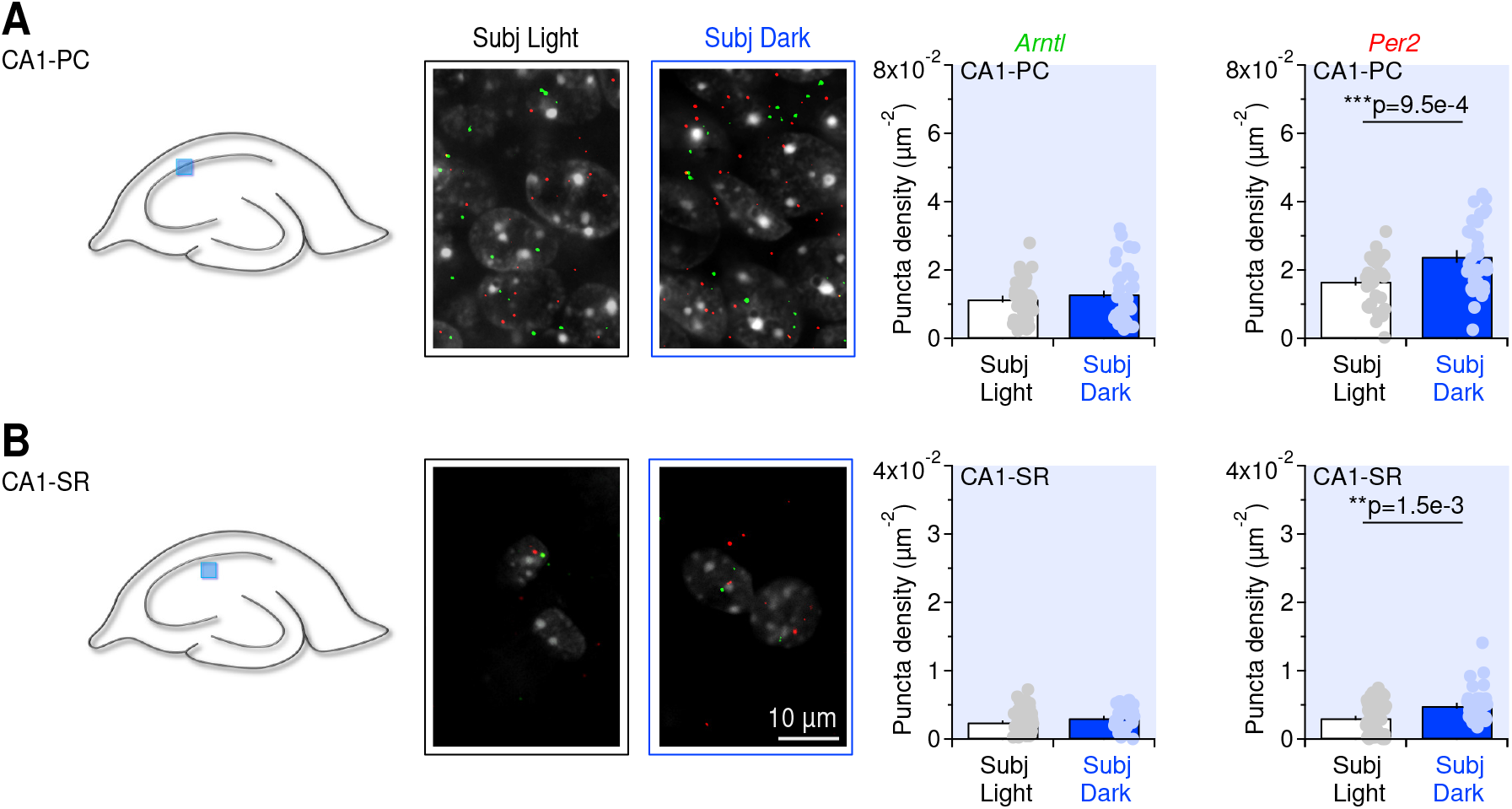
Hippocampal clock gene expression is preserved in mice maintained in constant darkness. **(A)** Schematic representation of a mid-sagittal section of the mouse brain. The light blue square represents the location of the region from which we collected confocal images. The green and red puncta represent the fluorescence signal obtained with RNAscope *in situ* hybridization for transcripts of clock genes *Arntl* (green) and *Per2* (red) in the CA1 pyramidal cell layer, slices prepared at ZT3.5 (Subjective light) and ZT15.5 (Subjective dark) from mice kept under 12H:12 D:D conditions. The bar charts on the right provide a summary of the *Arntl* and *Per2* puncta density. **(B)** As in (A), for *stratum radiatum*.

**Supplementary Figure 3.**
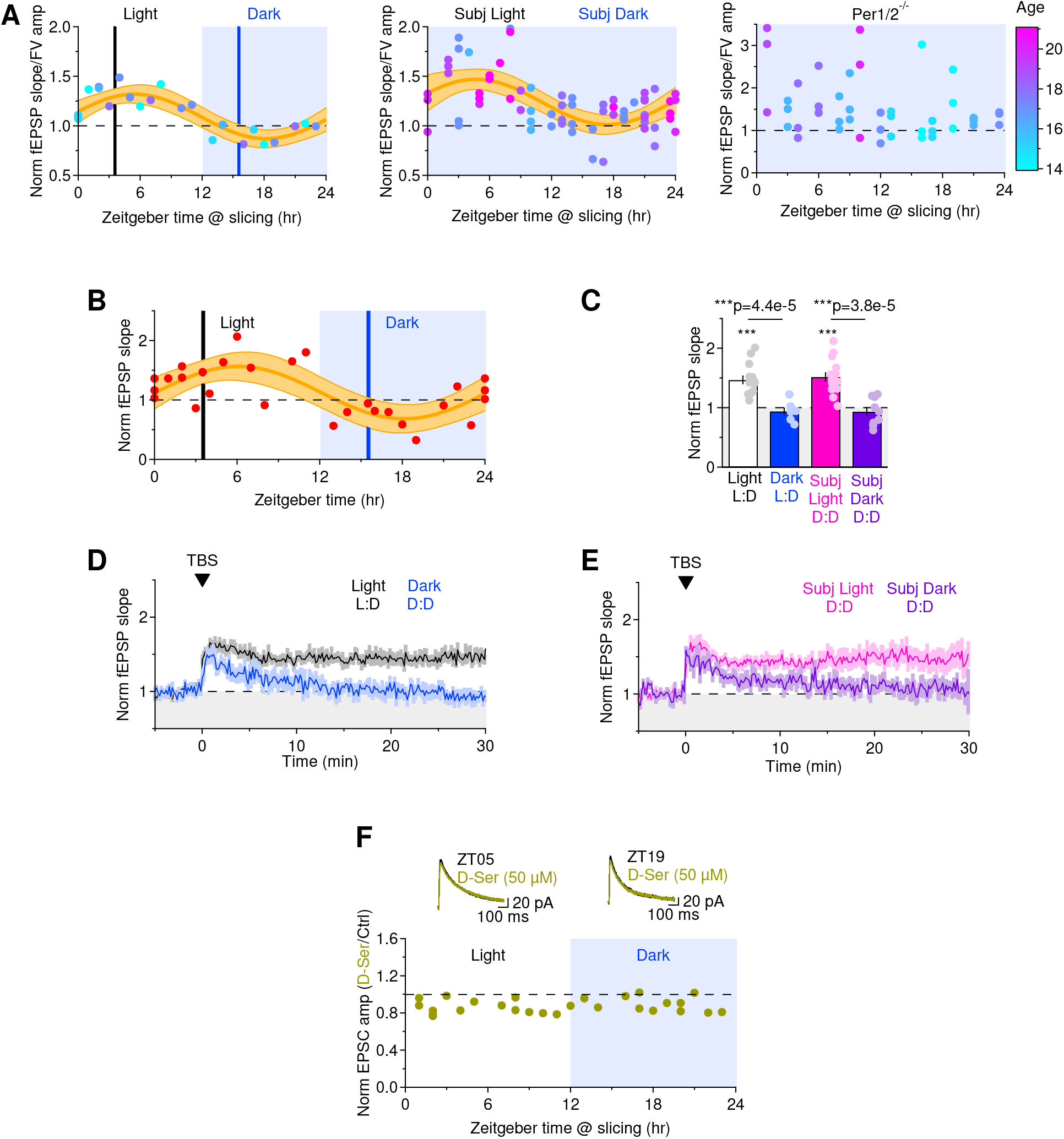
Validation of the circadian dependence of Schaffer collateral LTP. **(A)** In these graphs, thecolor of each dot represents the age of the mouse used to perform the experiments. NO correlation between the magnitude of LTP and age was detected in C57BL/6 mice kept in 12H:12H L:D (*left*) and D:D conditions (*middle*), or in Per1/2^-/-^ mice kept in constant darkness (*right*). **(B)** Circadian oscillations in Schaffer collateral LTP occur regardless of the method used to quantify LTP. The graph provides a measure of change in fEPSP slope in response to TBS, for the experiments described in Fig. 1D, *left*. **(C)** Summary of the effect of TBS on the fEPSP slope, in slices prepared in the L/D-phase of C57BL/6 mice maintained in 12H:12H L:D conditions, and in the subjective L/D-phase of C57BL/6 mice maintained in 12H:12H D:D conditions. **(D)** Time course of baseline-normalized field EPSP slope in slices prepared at ZT3.5 (Light, black) and ZT15.5 (Dark, blue), also shows in Fig. 1E, *top*. **(E)** Time course of baseline-normalized field EPSP slope in slices prepared at ZT3.5 (subjective light, magenta) and ZT15.5 (subjective dark, purple) from mice kept under constant darkness. **(F)** Effect of D-Ser (50 μM) on NMDA EPSCs recorded at a holding potential of 40 mV. Each data point represents the results obtained in slices from a single mouse. The traces represent the averages of 20 consecutive sweeps acquired under baseline conditions (black) or in the presence of D-Ser, at ZT5 (left) and ZT19 (right). We did not detect circadian oscillations in the effect of D-Ser on the NMDA EPSC amplitude.

**Supplementary Figure 4.**
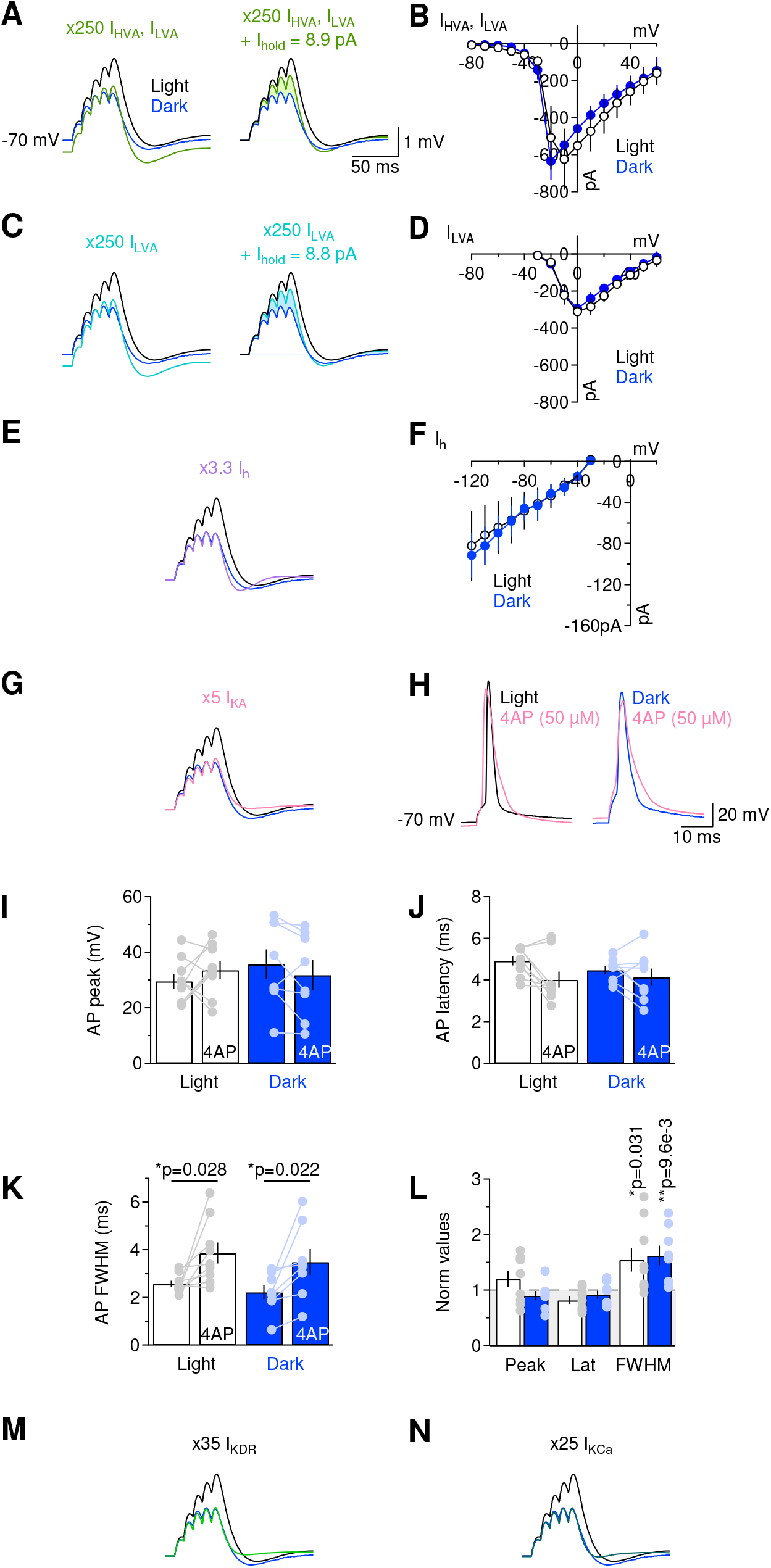
Comparison of modeling and experimental data on the contribution of different ionic conductances to the reduced temporal summation of excitatory inputs in the D-phase. **(A)** Results of NEURON multi-compartmental model mimicking changes in somatic AMPA and NMDA EPSP summation in the L/D-phases, evoked by activating a cohort of 100 synapses randomly distributed through the dendritic tree of CA1 pyramidal cells. The trace in green was obtained by imposing a 250-fold increase in the conductance of high- and low-voltage activated calcium channels (HVA and LVA, respectively) in the model used to mimic the results obtained in the L-phase condition. The traces on the right were obtained through the model by also injecting 8.9 pA positive holding current, to ensure the resting membrane potential was similar to that of L/D-phase simulations. **(B)** I/V plot obtained by recording HVA and LVA currents from CA1 pyramidal cells, evoked by 100 ms voltage step depolarization of increasing amplitude, from −80 mV to 60 mV. **(C)** As in (A), in response to a 250-fold increase in the conductance of LVA calcium channels (*left*) and after injection of 8.8 pA holding current (*right*). **(D)** I/V plot obtained by recording LVA currents from CA1 pyramidal cells, evoked by 100 ms voltage step depolarization of increasing amplitude, from −40 mV to 60 mV. **(E)** As in (A, *left*), in response to a 3.3-fold increase in the conductance of I_h_ channels. **(F)** I/V plot obtained by recording I_h_ currents from CA1 pyramidal cells, evoked by 100 ms voltage step depolarization of increasing amplitude, from −120 mV to 20 mV. **(G)** As in (A, *left*), in response to a 5-fold increase in the conductance of I_A_ potassium channels. **(H)** Example of action potentials recorded under control conditions, in slices prepared in the L/D-phase, and following application of the IA antagonist 4AP (50 μM). The action potentials were evoked by applying a 5-ms long, supra-threshold current step injection to CA1 pyramidal cells. **(I)** Summary graph and in-cell comparison of the effect of 4AP on the action potential peak. **(J)** Summary graph and in-cell comparison of the effect of 4AP on the action potential latency. **(K)** Summary graph and in-cell comparison of the effect of 4AP on the action potential full width at half maximum (FWHM). 4AP broadened the action potential to the same extent in slices prepared during the L/D-phases. **(L)** Summary graph of baseline-normalized action potential peak, latency and FWHM in the presence of 4AP. 4AP increases the action potential FWHM to the same extent in slices prepared during the L/D-phases. **(M)** As in (A, *left*), in response to a 35-fold increase in the conductance of delayed rectifier (I_DR_) potassium channels. **(N)** As in (A, *left*), in response to a 25-fold increase in the conductance of calcium-activated potassium channels (I_KCa_).

**Supplementary Figure 5.**
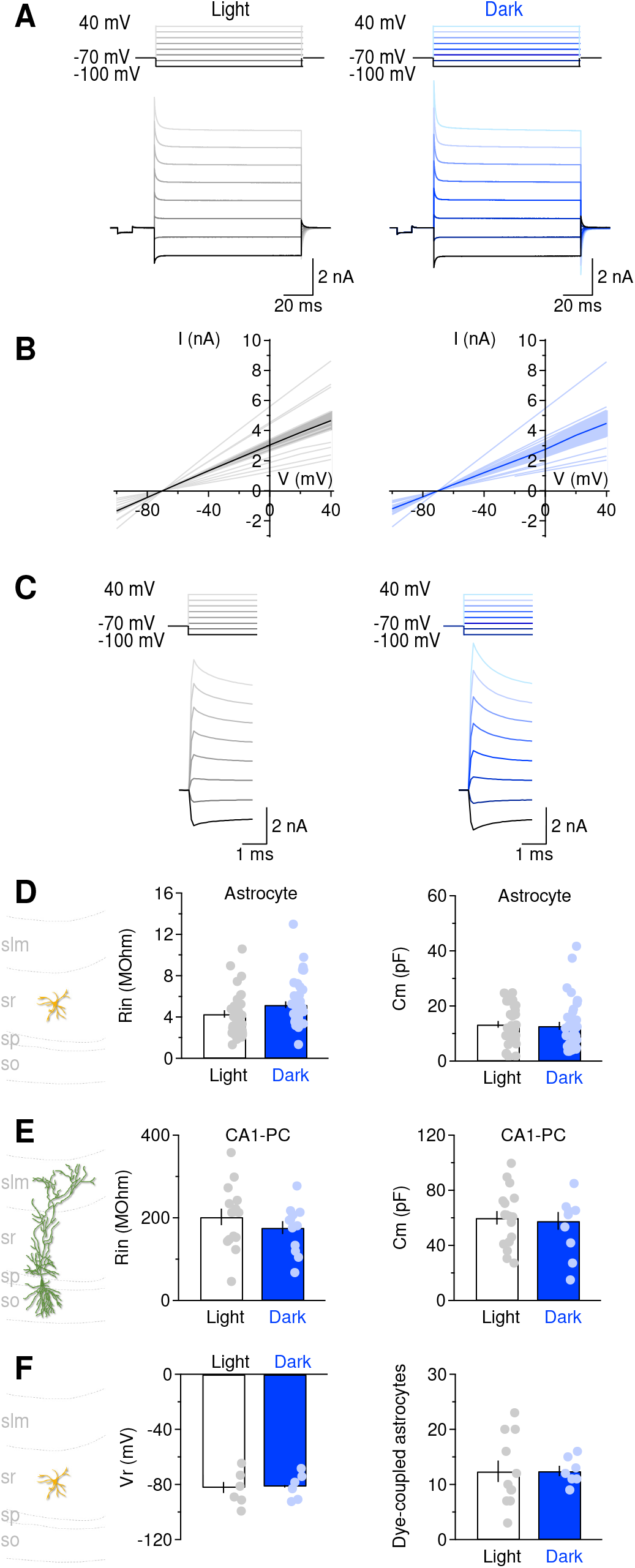
The passive membrane properties of astrocytes and neurons do not change during the L/D-phases. **(A)** Example of voltage step protocol and current responses recorded from astrocytes patched in *stratum radiatum* of acute hippocampal slices prepared during the L/D-phases. Each voltage step was 100 ms long. The holding membrane potential (−80 mV) was stepped from −100 mV up to 40 mV, in 20 mV voltage steps. Each current response represents the average of 20 responses from one astrocyte. A 10 ms, 5 mV hyperpolarizing voltage step was used to monitor changes in the series resistance of our recordings. Any recording in which the series resistance changed more than 20% compared to its initial value was discarded from the analysis. **(B)** I/V relationship derived from the experiment described in (A), during the L/D-phases. Thick lines and shaded areas represent mean±SEM. **(C)** Close-up view on the voltage step protocol and current responses. The decaying phase of the current responses provided information on the astrocyte capacitance. **(D)** *Left*, the astrocyte input resistance (R_in_) was similar during the L-phase (white bar and grey dots) and D-phase (blue bar and light blue dots). *Right*, the astrocyte membrane capacitance (C_m_) was similar during the L/D-phases. **(E)** *Left*, CA1 pyramidal cells had higher input resistance and membrane capacitance compared to astrocytes. The input resistance was similar during the L/D-phases. *Right*, the membrane capacitance was also similar during the L/D-phases. **(F)** *Left*, the resting membrane potential of astrocytes did not change significantly in the L/D-phases. *Right*, the number of dye-coupled astrocytes was measured in confocal images of biocytin-filled, streptavidin-AF488-labeled cells, and was similar during the L/D-phases.

**Supplementary Figure 6.**
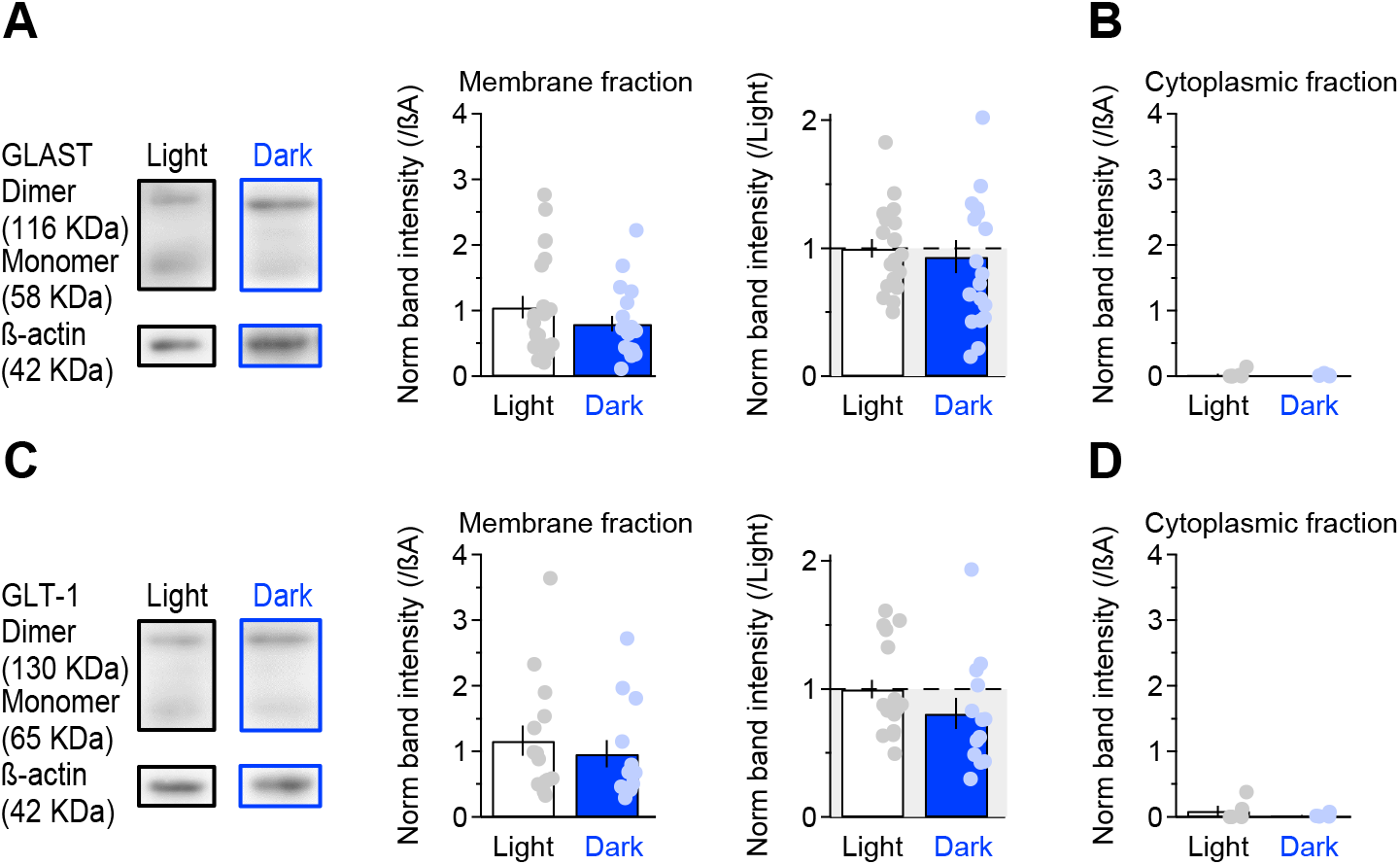
The plasma membrane expression of glutamate transporters GLAST and GLT-1 does not change with time-of-day. **(A)** Western blot analysis of biotinylated hippocampal GLAST expression. The two upper bands correspond to GLAST monomers and dimers (*left*). The summary graphs show the quantification of GLAST band intensity in the membrane fraction, normalized to the corresponding intensity of β-actin (*middle*), or divided by the normalized band intensity in the L-phase (*right*). **(B)** Quantification of GLAST band intensity in the cytoplasmic fraction, divided by the β-actin normalized band intensity in the L-phase (*right*). **(C,D)** As in (A,B), for GLT-1.

**Supplementary Figure 7.**
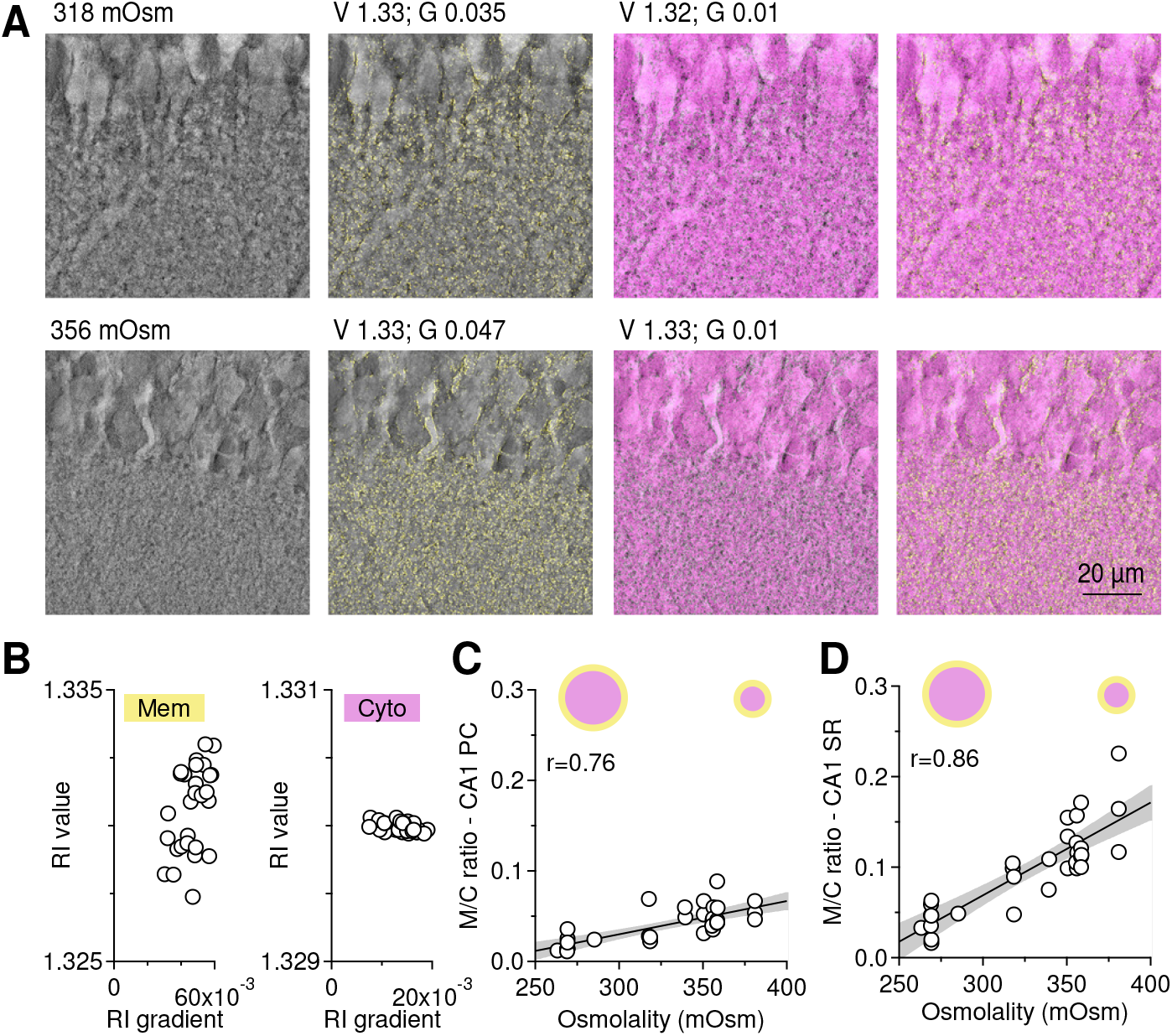
Validation of holotomography approach to measure cell shrinkage and swelling. **(A)** Example images of hippocampal area CA1, collected using holotomography in slices stored in extracellular solutions of different osmolality. Each image was digitally stained using refractive index (RI) value (V) and gradient (G) combinations that allowed labeling cell membranes (yellow) and cell cytoplasm (magenta). Cell shrinkage was detected at higher extracellular osmolality. **(B)** Scatter plot of RI value and gradient combinations use to label cell membranes (left, yellow) and cell cytoplasm (right, magenta). **(C)** Summary graph of M/C ratio measures in the CA1 pyramidal cell later. **(D)** As in (C), for measures collected from *stratum radiatum*.

**Supplementary Figure 8.**
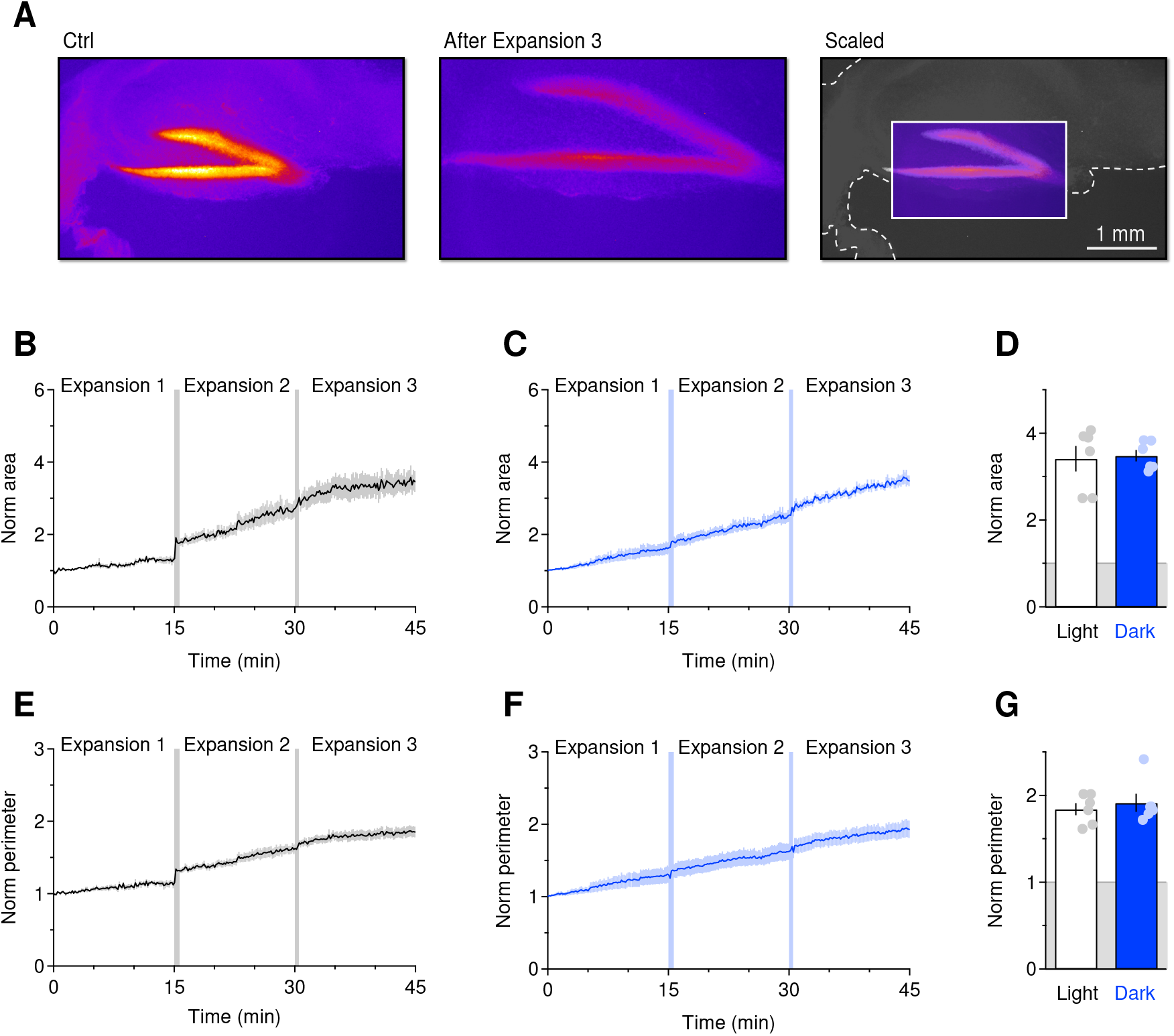
proExM provides a similar expansion of hippocampal slices prepared during the L/D-phases. **(A)** *Left*, fluorescence microscopy image of a hippocampal slice labelled with DAPI. The high cell density, bright area is the dentate gyrus. *Middle*, fluorescence microscopy image of the same hippocampal slice after three consecutive, 15 min long rounds of expansion. *Right*, overlaid image of the expanded dentate gyrus scaled down to the size of the hippocampal slice before expansion. This comparison shows that proExM generates an isotropic expansion of hippocampal slices. **(B)** Time course of hippocampal slice area increase during the proExM protocol, in slices prepared during the L-phase. **(C)** As in (B), in slices prepared during the D-phase. **(D)** Quantification of the proExM-induced surface area increase in hippocampal slices. **(E)** Time course of hippocampal slice perimeter increase during the proExM protocol, in slices prepared during the L-phase. **(F)** As in (E), in slices prepared during the D-phase. **(G)** Quantification of the proExM-induced perimeter increase in hippocampal slices.

**Supplementary Figure 9.**
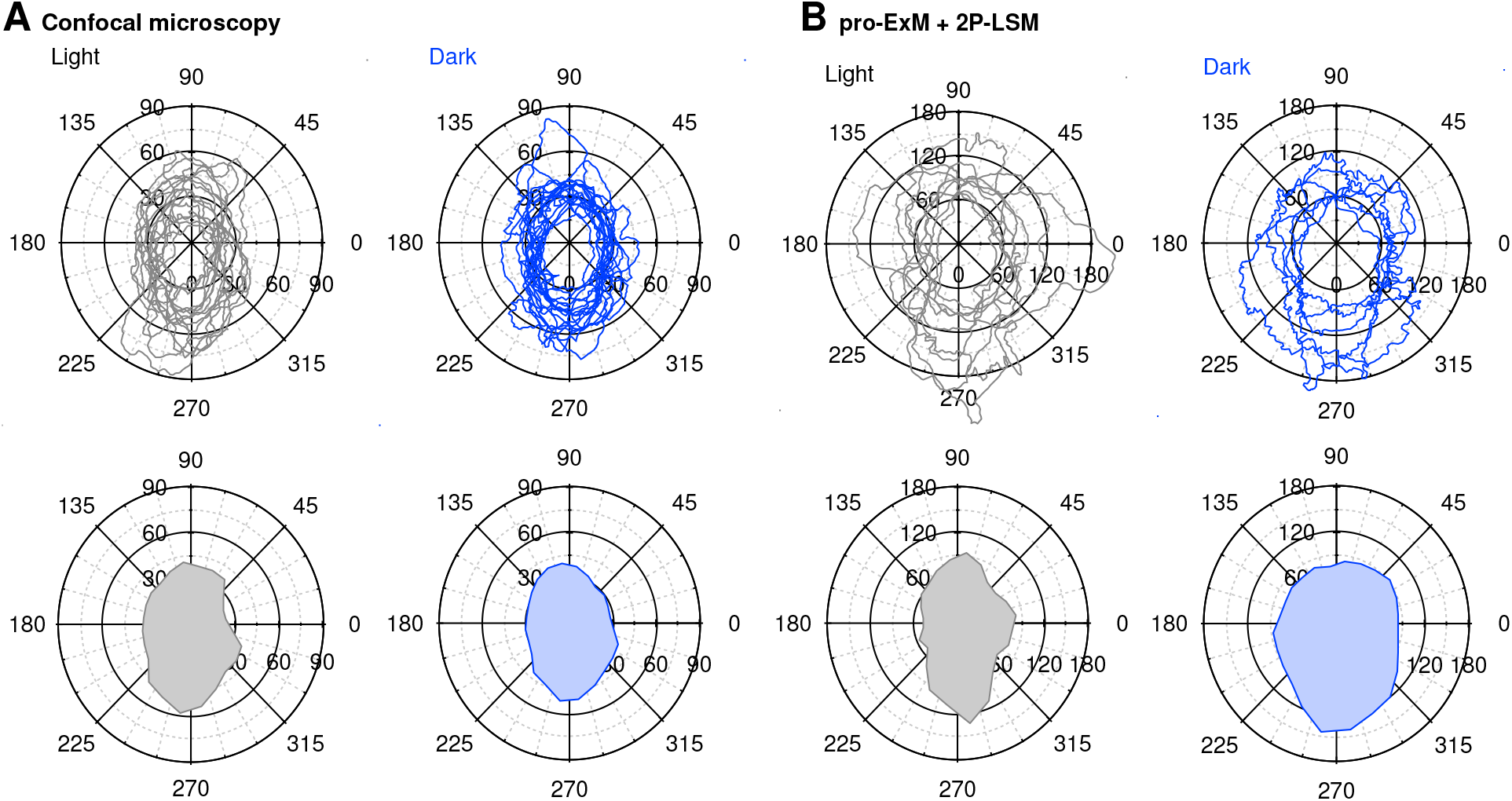
Polar plots used to estimate the astrocyte coverage area. **(A)** Polar plots of astrocyte contours after confocal imaging (*top*) and their corresponding averages (*bottom*). **(B)** Polar plots of astrocyte contours after proExM and 2P-LSM (*top*) with their corresponding averages (*bottom*).

## STAR Methods

### Ethics statement

All experimental procedures were performed in accordance with protocols approved by the Institutional Animal Care and Use Committee at the State University of New York (SUNY) Albany and guidelines described in the National Institutes of Health’s Guide for the Care and Use of Laboratory Animals.

### Mice

Unless otherwise stated, all C57BL/6 mice were group housed and kept under 12H:12H L:D conditions (lights on at 7 AM, ZT0; lights off at 7 PM, ZT12). We refer to the ZT0-12 time interval as the L-phase, and the ZT12-24 time interval as the D-phase. The ZT time sets the origin of the 24 hour period to the beginning of the L-phase, facilitating cross-study comparisons independently of the actual clock-time settings of different animal facilities. A subset of experiments were performed using mice kept under constant darkness (i.e. 12H:12H D:D conditions under red dim light illumination for at least three cycles). The dim red light intensity in constant darkness matched that of the D-phase for mice kept under 12H:12H L:D conditions. For mice maintained in constant darkness, the ZT0-12 time interval was referred to as the subjective L-phase, and the ZT12-24 time interval as the subjective D-phase. Mice with targeted disruption of *Per1* and *Per2* alleles (Per1^Brdm1/Brdm1^: Per2^Brdm1/Brdm1^), here referred to as Per1/2^-/-^ mice, were a generous gift from Dr. Jennifer Hurley (RPI) and have been characterized in previous works (Bae et al., 2001; Shiromani et al., 2004; Zheng et al., 2001). Food and water were available *ad libitum* to all mice throughout the 24 hour period. Unless otherwise stated, all dissections for the data collected in the L- and subjective L-phase were performed at ZT3.5. All dissections for the D- and subjective D-phase data were performed at ZT15.5.

Genotyping was performed on toe tissue samples of P7–P10 mice. Briefly, tissue samples were digested at 55°C overnight with shaking at 330 rpm in a lysis buffer containing the following (in mM): 100 Tris base, pH 8, 5 EDTA, and 200 NaCl, along with 0.2% SDS and 50 μg/ml proteinase K. Following heat inactivation of proteinase K at 97°C for 10 min, DNA samples were diluted 1:1 with nuclease-free water. The PCR primers used for Per1 and Per2 were purchased from Eurofins Genomics and their nucleotide sequences are listed in **Table 1**. The forward and reverse primers for *Per1* are located in exons 3 and 4, respectively. The reverse primer for *Per1^-/-^* mice is located in the mPGK promoter for the *Hprt* cassette that replaces exons 4 through 18 of *Per1^-/-^* mice. The forward and reverse primers for *Per2* are located within intron 11-12. The reverse primer for *Per2^-/-^* mice is located in the Neomycin cassette that replaces the highly conserved PAS domain in *Per2^-/-^* mice. PCR was carried out using the KAPA HiFi Hot Start Ready Mix PCR Kit (KAPA Biosystems, KK2602). Briefly, 12.5 μl of 2X KAPA HiFi Hot Start Ready Mix was added to 11.5 μl of a diluted primer mix (0.5-0.75 μM final concentration for each primer) and 1 μl of diluted DNA. The PCR cycling protocols for *Per1* and *Per2* are described in **Tables 2 and 3**, respectively.

**Table 1.**
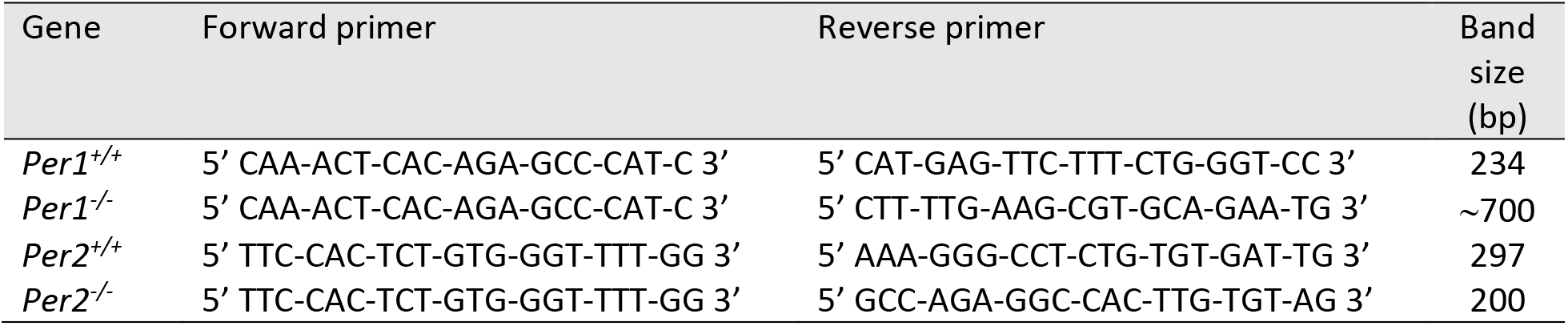
Sequence of genotyping primers

**Table 2.** PCR protocol for *Per1*

|  | Initiation/<br>melting | Denaturation | Annealing | Elongation | Amplification | Hold |
| --- | --- | --- | --- | --- | --- | --- |
| Temperature (°C) | 95 | 95 | 60 | 72 | 72 | 4 |
| Duration (min) | 3 | 0.25 | 0.25 | 0.5 | 1 | ∞ |
| Cycles | 1 | 35 |  |  | 1 |  |

**Table 3.**
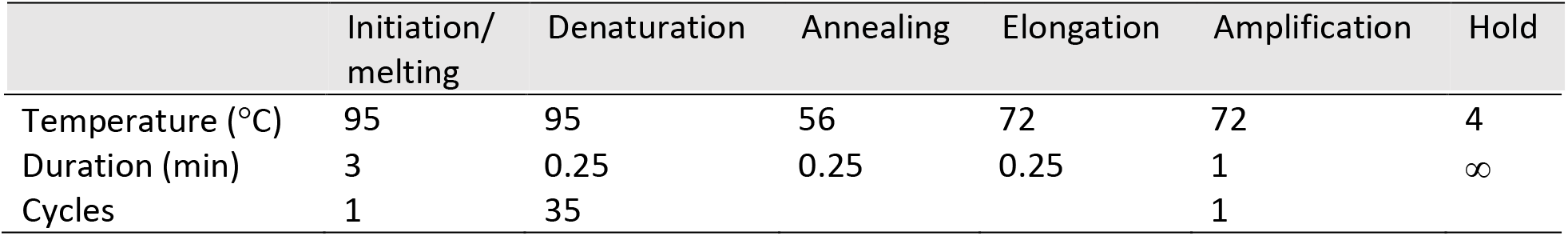
PCR protocol for *Per2*

### In vivo chronic temperature recordings

To record the mouse body temperature, we implanted programmable micro-transponders (Cat# IPTT300; BDMS) under the skin of the neck of P12 mice. We took temperature measures hourly from P14–21 using a wireless hand-held reader for IPTT micro-transponders (Cat# DAS-8007-IUS, BDMS). Data analysis was performed in IgorPro 6.37 (Wavemetrics, Lake Oswego, OR) using custom-made software (A.Sc.).

### Fluorescent in situ hybridization of clock gene transcripts using RNAscope

We dissected the brain of P14-21 C57BL/6 mice at ZT3.5 or ZT15.5, removed the olfactory bulbs, cerebellum and temporal lobes and fixed it with 4% PFA/PBS overnight at 4°C. The brain was then cryoprotected in 30% sucrose PBS at 4°C for 48 hr and stored in PBS for no more than a week. To prepare slices for RNAscope, we separated the two hemispheres, embedded them in agar and prepared 40 μm thick slices using a vibrating blade microtome (VT1200S, Leica Microsystems, Buffalo Grove, IL). The slices were post-fixed in 4% PFA/PBS for 30 min at room temperature (RT) and mounted onto Superfrost plus microscope slides. Mounted slices were used for fluorescence in situ hybridization (FISH) using an RNAscope multiplex fluorescent assay (Advanced Cell Diagnostics, Newark, CA) according to manufacturer instructions, using Mm-Per2-C1 and Mm-Arntl-C2 RNA probes and Opal 520 and Opal 570 dyes (Akoya Biosciences, Menlo Park, CA). DAPI Fluoromount G was used as the mounting medium (Cat# 0100-02; SouthernBiotech, Birmingham, AL). The presence of *Per2* and *Arntl* transcripts was assessed using a confocal microscope (Zeiss LSM710) equipped with a Plan-Apochromat 63X/1.4NA oil objective. Image size was set to 1024×1024 pixels and represented the average of 8 consecutive frames.

### Acute slice preparation and electrophysiology recordings

Acute coronal slices of the mouse hippocampus were obtained from C57BL/6 mice of either sex (P14–21), deeply anesthetized with isoflurane and decapitated in accordance with SUNY Albany Animal Care and Use Committee guidelines. The brain was rapidly removed and placed in ice-cold slicing solution bubbled with 95% O_2_/5% CO_2_ containing the following (in mM): 119 NaCl, 2.5 KCl, 0.5 CaCl_2_, 1.3 MgSO_4_·H_2_O, 4 MgCl_2_, 26.2 NaHCO_3_, 1 NaH_2_PO_4_, and 22 glucose, 320 mOsm, pH 7.4. The slices (250 μm thick) were prepared using a vibrating blade microtome (VT1200S; Leica Microsystems, Buffalo Grove, IL). Once prepared, the slices were stored in slicing solution in a submersion chamber at 36°C for 30 min and at RT for at least 30 min and up to 5 hr. Unless otherwise stated, the recording solution contained the following (in mM): 119 NaCl, 2.5 KCl, 2.5 CaCl_2_, 1 MgCl_2_, 26.2 NaHCO_3_, and 1 NaH_2_PO_4_, 22 glucose, 300 mOsm, pH 7.4. We identified the hippocampus under bright field illumination using an upright fixed-stage microscope (BX51 WI; Olympus Corporation, Center Valley, PA). To record evoked currents, we delivered constant voltage stimuli (50–100 μs) to a bipolar stainless steel electrode (Cat# MX21AES(JD3); Frederick Haer Corporation, Bowdoin, ME) positioned in *stratum radiatum*, ~100 μm away from the recorded cell. Whole-cell, voltage-clamp patch-clamp recordings were made using patch pipettes containing (in mM): 120 CsCH_3_SO_3_, 10 EGTA, 20 HEPES, 2 MgATP, 0.2 NaGTP, 5 QX-314Br, 290 mOsm, pH 7.2. All recordings were obtained using a Multiclamp 700B amplifier (Molecular Devices, San Jose, CA) and filtered at 10 KHz, converted with an 18-bit 200 kHz A/D board (HEKA Instrument, Holliston, MA), digitized at 10 KHz, and analyzed offline with custom-made software (A.Sc.) written in IgorPro 6.37 (Wavemetrics, Lake Oswego, OR). Patch electrodes (#0010 glass; Harvard Apparatus, Holliston, MA) had tip resistances of ~5 MΩ. Series resistance (~20 MΩ) was not compensated, but was continuously monitored and experiments were discarded if this changed by >20%. The series resistance was subtracted from the total resistance to estimate the cell membrane resistance of neurons and astrocytes. Extracellular recordings were performed using electrodes with a tip resistance of 2-5 MΩ, filled with the recording solution described above. The electrode resistance was continuously monitored and experiments were discarded if this changed by >20%. Slices were incubated with NR3C1/2 receptor antagonists for 30 min −2 hr before being used for electrophysiology recordings. Tetrodotoxin (TTX), 2,3-Dioxo-6-nitro-1,2,3,4-tetrahydrobenzo[f]quinoxaline-7-sulfonamide disodium salt (NBQX) and D,L-2-Amino-5-phosphonopentanoic acid (APV) were purchased from Hello Bio (Princeton, NJ; Cat# HB1035, HB0443, HB0251, respectively). (3S)-3-[[3-[[4-(Trifluoromethyl)benzoyl]amino]phenyl]methoxy]-L-aspartic acid (TFB-TBOA) was purchased from Tocris (Minneapolis, MN; Cat# 2532). Spironolactone, mifepristone and luzindole were purchased from Cayman Chemical Company (Ann Arbor, MI; Cat# 9000324, 10006317, 15598, respectively). All other chemicals were purchased from MilliporeSigma (Burlington, MA). All recordings were performed at RT.

### Blood collection and corticosterone radioimmunoassay

Trunk blood was collected from 18 mice kept under 12H:12H L:D conditions, and 18 mice kept under 12H:12H D:D conditions. Trunk blood was collected within 3 min of cage disturbance from 3 mice every 4 hours. Blood was centrifuged at 5500 RCF for 10 min, and supernatant was transferred to a new tube and stored at −80°C until used for radioimmunoassay for corticosterone. Plasma corticosterone was assessed using a commercial I-125 corticosterone radioimmunoassay kit according to the manufacturer’s instructions (Cat# 07120102, MP Biochemicals, LLC, Orangeburg, NY, USA). The intra-assay coefficient of variation was 3.4%. This assay kit shows a slight cross-reaction with desoxycorticosterone 0.34%, testosterone 0.10%, cortisol 0.05%, aldosterone 0.03%, and progesterone 0.02%.

### Biocytin filling and confocal imaging

Biocytin 0.2–0.4% (w/v) was added to the intracellular solution used to patch astrocytes and neurons. Each cell was patched and filled for at least 20 min. The slices were then fixed overnight at 4°C in 4% PFA/PBS, cryo-protected in 30% sucrose PBS, and incubated in 0.1% streptavidin-Alexa Fluor 488 conjugate and 0.1% Triton X-100 for 3 hr at RT. The slices were then mounted onto microscope slides using Fluoromount-G mounting medium (Cat# 0100-01; SouthernBiotech, Birmingham, AL). Confocal images were acquired using a Zeiss LSM710 inverted microscope equipped with 488 nm Ar laser. All images were acquired as z-stacks using a 40X/1.4 NA oil-immersion objective. To visualize full cells, we stitched together z-stacks (1024×1024 pixels) collected by averaging four frames for each focal plane (1 μm z-step). The image analysis to measure the astrocyte coverage area was performed using Fiji (https://fiji.sc/). Briefly, we generated a maximum intensity projection of each image stack and manually traced the contour of the outer boundaries of the area of the neuropil covered by each astrocyte. Confocal images that contained more than one filled astrocyte were discarded from the analysis, because they introduced inaccuracies in the derived contours. The polar plots were obtained by centering the soma of each astrocyte at the center of a polar plot. The main primary branch of each astrocyte was oriented along the 270° angle. Average coverage traces were obtained using IgorPro (Wavemetrics, Lake Oswego, OR).

### Protein-retention expansion microscopy and two-photon imaging

All experiments were performed according to the proExM protocol described by (Tillberg et al., 2016). Briefly, to measure the expansion factor (**Supplementary Fig. 8**) we fixed hippocampal slices with 4% PFA/PBS overnight at 4°C, cryoprotected them in 30% sucrose PBS and stored them in PBS. The day before starting the proExM protocol, we incubated them with DAPI Fluoromount G overnight at 4°C (Cat# 0100-20; SouthernBiotech, Birmingham, AL). Slices containing biocytin filled astrocytes were fixed, cryo-protected, stored in PBS and labelled with streptavidin-Alexa Fluor 488 as described in the previous section. In both cases, each slice was then incubated with 200 μl of anchoring solution overnight at RT. The following day, the slices were gelled and digested with Proteinase K overnight at RT. Subsequently, they were expanded using three consecutive incubations with distilled water for 15 min each. Slices stained with DAPI were imaged as they expanded during each incubation period, using a 2X/0.06 NA air objective on an EVOS FL Cell Imaging System equipped with DAPI filter set (λ_ex_: 357/44 nm, λ_em_: 447/60 nm; ThermoFisher Scientific, Waltham, MA). Images were collected at 0.1 fps and manually traced to measure slice perimeter and surface area using IMOD (https://bio3d.colorado.edu/imod/). The expanded gels containing biocytin-filled astrocytes were then covered with 2% agarose and submerged in distilled water before being imaged with a custom-made two-photon laser-scanning microscope. The two-photon laser scanning system (Scientifica, Clarksburg, NJ) was powered by a Ti:sapphire pulsed laser (Coherent, Santa Clara, CA) tuned to 760 nm and connected to an upright microscope with a 60X/1.0 NA objective (Olympus Corporation, Center Valley, PA). The green fluorescent signal was separated from the red using a 565 nm dichroic mirror and filtered using FITC filter sets (Olympus Corporation, Center Valley, PA). We averaged eight frames for each optical section (512×512 pixels) and collected z-stacks for a total distance of ~200 μm (in 1.2 μm steps). The composition of the anchoring, gelling and digestion solution is reported below.

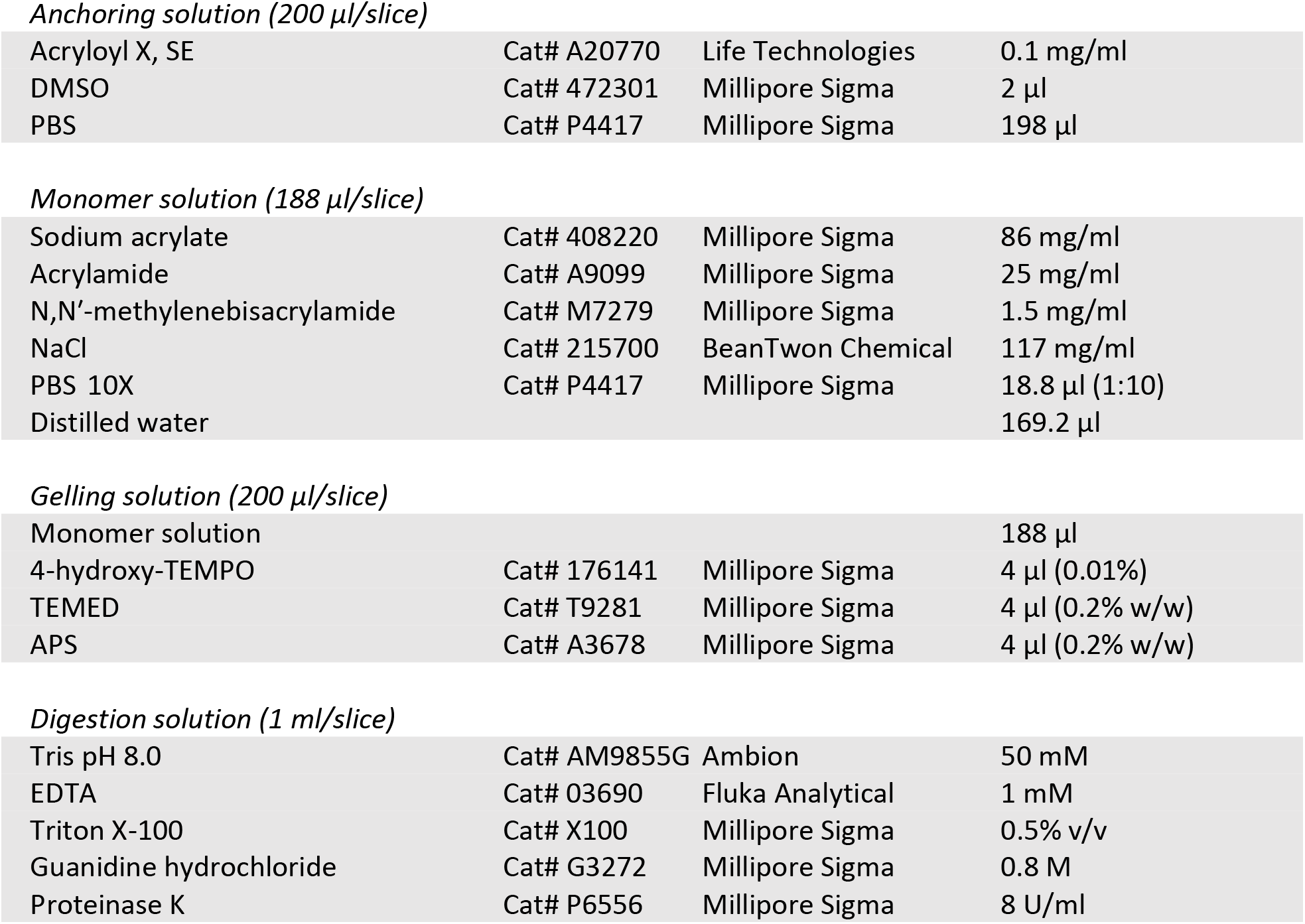

|  |  |  |  |
| --- | --- | --- | --- |
| Acryloyl X, SE | Cat# A20770 | Life Technologies | 0.1 mg/ml |
| DMSO | Cat# 472301 | Millipore Sigma | 2 μl |
| PBS | Cat# P4417 | Millipore Sigma | 198 μl |
| *Monomer solution (188 $\mu$ l/slice)* |  |  |  |
| Sodium acrylate | Cat# 408220 | Millipore Sigma | 86 mg/ml |
| Acrylamide | Cat# A9099 | Millipore Sigma | 25 mg/ml |
| N,N'-methylenebisacrylamide | Cat# M7279 | Millipore Sigma | 1.5 mg/ml |
| NaCl | Cat# 215700 | BeanTwon Chemical | 117 mg/ml |
| PBS 10X | Cat# P4417 | Millipore Sigma | 18.8 μl (1:10) |
| Distilled water | | | 169.2 μl |
| *Gelling solution (200 $\mu$ l/slice)* |  |  |  |
| Monomer solution | | | 188 μl |
| 4-hydroxy-TEMPO | Cat# 176141 | Millipore Sigma | 4 μl (0.01%) |
| TEMED | Cat# T9281 | Millipore Sigma | 4 μl (0.2% w/w) |
| APS | Cat# A3678 | Millipore Sigma | 4 μl (0.2% w/w) |
| *Digestion solution (1 ml/slice)* |  |  |  |
| Tris pH 8.0 | Cat# AM9855G | Ambion | 50 mM |
| EDTA | Cat# 03690 | Fluka Analytical | 1 mM |
| Triton X-100 | Cat# X100 | Millipore Sigma | 0.5% v/v |
| Guanidine hydrochloride | Cat# G3272 | Millipore Sigma | 0.8 M |
| Proteinase K | Cat# P6556 | Millipore Sigma | 8 U/ml |

### Western blot

Western blot experiments were performed on protein extracts from the hippocampus of mice of either sex aged P14–21, sacrificed at ZT3.5 or ZT15.5. Membrane proteins were extracted using the Mem-PER Plus Membrane Protein Extraction Kit (Cat# 89842; ThermoFisher Scientific, Waltham, MA) according to the manufacturer’s instructions using a mixture of protease and phosphatase inhibitors (10 μl/ml, Cat# 78441; ThermoFisher Scientific, Waltham, MA). The total membrane protein concentration was determined by spectrophotometry. Equal amounts of protein (100 μg) from each sample were resolved on 10 or 12% acrylamide gels. The proteins were transferred to PVDF membranes (Cat# P2563; MilliporeSigma, Burlington, MA) using a semidry blotting approach. The membranes were blocked with 5% nonfat milk in TBST pH 7.6, and probed with primary antibodies overnight at 4°C in 5% BSA in TBST, pH 7.6. Secondary antibody incubation was performed for 1–2 hr at RT in 5% nonfat milk in TBST, pH 7.6. Pre-adsorption experiments were performed using the control antigen provided by the primary antibody supplier according to the manufacturer’s instructions (1 μg/μg antibody). Membranes were probed with either rabbit anti-GLAST (1:1,000), anti-GLT-1 (1:1,000) or β-actin antibodies (1:1,000). Biotinylated horse anti-rabbit antibody was used as the secondary antibody for all blots (1:1,000 for GLAST and GLT-1, 1:5,000 for β-actin). We amplified the immuno-labeling reactions using the Vectastain ABC kit (1:2,000 for GLAST and GLT-1; 1:5,000 for β-actin) and the Clarity Western ECL system served as the substrate for the peroxidase enzyme (Cat# 1705060, Bio-Rad, Hercules, CA). For semi-quantitative analysis, protein band intensities were collected as 16-bit images using a digital chemiluminescence imaging system (c300, Azure Biosystems, Dublin, CA) at different exposure times (0.5–200 s). Each image was converted to an 8-bit image for image analysis using Fiji software (https://fiji.sc/). Only images collected at exposure times that did not lead to pixel saturation were included in the analysis. The intensity of each band was calculated as the mean gray value in a region of interest (ROI) surrounding each band in three images collected at different exposure times. All GLAST and GLT-1 band intensities were normalized to the band intensity of β-actin in the same lane. Only images collected at exposure times that did not lead to pixel saturation were used to quantify band intensities.

A list of antibodies and reagents, with their RRID, is reported below:

|  |  |  |
| --- | --- | --- |
| RRID: AB_10695722 | rabbit anti GLAST | Cell Signaling Technology, Danvers, MA |
| RRID: AB_2190743 | rabbit anti GLT-1 | Cell Signaling Technology, Danvers, MA |
| RRID: AB_2223172 | rabbit anti β-actin | Cell Signaling Technology, Danvers, MA |
| RRID: AB_2336201 | biotinylated horse anti-rabbit | Vector Laboratories, Burlingame, CA |
| RRID: AB_2336819 | Vectastain ABC kit | Vector Laboratories, Burlingame, CA |

### Surface protein biotinylation assay

We prepared acute hippocampal slices following the same experimental procedures used to prepare acute slices for electrophysiology recordings. Once prepared, the slices were stored in slicing solution in a submersion chamber at 36°C for 30 min and at RT for at least 30 min and up to 1 hr. Subsequently, the slices were processed following the biotinylation protocol developed and validated by (Thomas-Crusells et al., 2003). *First*, the acute hippocampal slices were washed twice for 5 min in ice-cold slicing solution, here referred to as modified artificial cerebrospinal fluid (mACSF), and then transferred to a 24-well plate containing mACSF added with 100 μM sulfo-NHS-SS-biotin (Cat# PI21328, VWR, Radnor, PA) for 45 min, on ice. The slices were then washed twice with cold mACSF containing 1 mM L-lysine (Cat# L5501, Millipore Sigma, Burlington, MA), to block all excess reactive sulfo-NHS-SS-biotin. We dissected the hippocampus out of the biotinylated slices, pooled the slices in an Eppendorf tube and removed the excess supernatant. The membrane proteins were isolated according to the Mem-Per Plus Membrane Protein Extraction Kit (Cat#89842, ThermoFisher Scientific, Waltham, MA). Subsequently, the membrane protein fraction was incubated with 40 μl of Ultra-link immobilized Streptavidin beads (Pierce Streptavidin UltraLink Resin, Cat# 53113, ThermoFisher Scientific, Waltham, MA) with gentle rotation overnight at 4°C. The following day, the beads were washed three times (5,000 × g at 4°C for 2 min) in solubilization buffer containing protease inhibitors and biotinylated proteins were released from the beads with 40 μl of solubilization buffer containing 5% β-mercaptoethanol and stored at −20°C until further use. The biotinylated proteins were separated by 10% SDS-PAGE and transferred to an Immobilon-P PVDF membrane (Cat# P2563, Millipore Sigma, Burlington, MA). The membranes were blocked at RT for 1 h with 5% BSA (Cat# A9418, Millipore Sigma, Burlington, MA) in Tris Buffered Saline, pH 7.5, containing 0.1% Tween-20 (TBS-T), and immunoblotted overnight at 4°C with rabbit anti-GLAST (1:1,000), GLT-1 (1:1,000) or β-actin (1:5,000) (Cat# listed above, Cell Signaling Technology, Danvers, MA) in 5% BSA/TBS-T. The secondary antibody, biotinylated horse anti-rabbit (Vector Laboratories, Burlingame, CA) was incubated for 1 h at RT in 5% BSA/TBS-T (1:1,000 for GLAST and GLT-1; 1:5,000 for β-actin). Immuno-labeling reactions were amplified using the Vectastain ABC kit (1:2,000 for GLAST and GLT-1; 1:5,000 for β-actin) and the Clarity Western ECL system served as the substrate for the peroxidase enzyme (Cat# 1705060, Bio-Rad, Hercules, CA). For semi-quantitative analysis, protein band intensities were collected as 16-bit images using a digital chemiluminescence imaging system (c300, Azure Biosystems, Dublin, CA) at different exposure times (0.5–200 s), as described for the Western blot analysis. Membranes were stripped using a mild stripping solution of 200 mM glycine, 0.1% SDS, 1% Tween-20, pH 2.2 with shaking at RT for 15 min, and re-probed with different antibodies.

### Electron microscopy and axial STEM tomography

Acute hippocampal slices processed for electron microscopy analysis were prepared as described for the electrophysiology experiments, from P17 mice. Slices were microwave-fixed for 13 s in 6% glutaraldehyde, 2% PFA, 2 mM CaCl_2_ in 0.1 N sodium cacodylate buffer and stored overnight at 4°C. After three washes in 0.1 N cacodylate buffer, we cut samples from the middle part of CA1 *stratum radiatum*, ~100 μm away from the pyramidal cell layer. These samples were treated with 1% osmium tetroxide for 1 hr on ice, en bloc mordanted with 0.25% uranyl acetate at 4°C overnight, washed and dehydrated with a graded series of ethanol, and embedded in epoxy resins. Thin sections (70–90 nm) were counterstained with lead citrate and uranyl acetate and examined on a JEOL 1200 EX transmission electron microscope. Images were collected with a CCD digital camera system (XR-100, AMT, Woburn, MA). To visualize the arrangement of pre-synaptic terminals, post-synaptic terminals, and astrocytic processes, thick sections (~1 μm) were cut from regions of CA1 *stratum radiatum* and electron tomograms were collected in a 300 kV electron microscope operated in the scanning transmission electron microscopy (STEM) mode, as described previously (Hohmann-Marriott et al., 2009; Sousa et al., 2011). A sample thickness of 1 μm – enabled by axial STEM tomography – provided sufficient sample depth to visualize features of interest in their entirety, such as synapses. In contrast to standard TEM tomography, conventional TEM tomography is limited to a specimen thickness of ~400 nm and cannot be applied to such thick sections because the transmitted electrons undergo multiple inelastic scattering processes, resulting in images that are blurred by chromatic aberration of the objective lens. Axial STEM tomography is not affected by chromatic aberration because the objective lens that forms the electron probe is in front of the specimen. Dual-axis tilt series of selected sections were recorded using an FEI Tecnai TF30 TEM/STEM operating at 300 kV (1.5° tilt increment, tilt range from 55° to −55°, pixel size = 5.6 nm). Image registration, tomogram generation, tracing, surface area and volume measures were performed using IMOD 4.7 (http://bio3d.colorado.edu/imod/). In the tomograms, we identified astrocytic processes based on their lack of synaptic vesicles and post-synaptic densities and because they did not give rise to pre- or post-synaptic terminals. Orthoslices through the STEM tomograms showed that astrocytic processes contained glycogen granules, intermediate filament bundles and a more electron-lucent cytoplasm with respect to that of neurons. The astrocytic processes were traced for the entire thickness of the reconstructed volume (1–1.5 μm). We reconstructed all astrocytic processes and all PSDs in each block. The reconstructed volumes were converted into object files and imported into the open-source software Blender 2.76 (https://www.blender.org/). Astrocyte-PSD distance was measured in Blender using custom-made analysis software written in Python (Sweeney et al., 2017).

### Tomographic holographic 3D microscopy

The measures of the cell membrane to cytoplasm ratio were obtained from acute hippocampal slices, prepared as described in the electrophysiology section of the methods, from animals aged P14–21 at ZT3.5 (L-phase) or ZT15.5 (D-phase). We used slices prepared at ZT3.5 for our osmolality test. Briefly, we prepared different batches of the slice recording solution, with different concentrations of NaCl (95, 107, 119, 131, 143 mM). We measured the osmolality of these solutions using a vapor pressure osmometer (Wescor 5600, Wescor, Logan, UT). Each slice was stored in one of these solutions for 1 hr, then it was fixed overnight with 4% PFA/PBS and cryo-protected in 30% sucrose PBS. The slices were then resectioned at 40 μm thickness using a vibrating blade microtome (VT1200S, Leica Microsystems, Buffalo Grove, IL) and mounted on a microscope slide using Fluoromount-G mounting medium (Cat# 0100-01; SouthernBiotech, Birmingham, AL). The slices were then imaged using the 3D Cell Explorer holographic tomographic 3D microscope (Nanolive, Ecublens, Switzerland) equipped with a 60X objective (NA=0.8), a class I laser source at λ=520 nm with a power output of 0.1 mW (sample exposure = 0.2 mW/mm^2^), and a USB 3.0 CMOS Sony IMX174 sensor providing 1024×1024 pixels at 165 fps (Cotte et al., 2013). The output data were acquired as image stacks of 512×512×96 voxels (0.186×0.186×0.372 μm) with a rate of tomogram acquisition of 0.5 s^−1^. This label-free non-phototoxic type of microscopy measures the refractive index (RI) in 3D samples, and allows users to visualize it using different levels of tolerance referred to as the refractive index gradient (IG). The acquired images were processed using Software for Tomographic Exploration of living cElls (STEVE) and Fiji (https://fiji.sc/). By using STEVE, we digitally stained the cells’ membrane (RI = 1.327–1.333; IG = 0.015–0.070) and cytoplasm (RI = 1.329–1.330; IG = 0.007–0.021). The RI and IG settings were varied across slices, but remained constant for our measures in the pyramidal cell layer and *stratum radiatum* of hippocampal area CA1. By using Fiji, we calculated the number of digitally stained pixels in three ROIs (200×100×40 pixels) positioned in the CA1 pyramidal cell layer and *stratum radiatum* of each slice. The center of the ROIs along the z-axis was positioned half way through the original z-stack acquired using the 3D Cell Explorer.

### NEURON modelling

3D z-stacks of biocytin filled CA1 pyramidal neurons were acquired using a Zeiss LSM710 confocal microscope and saved as czi files. The czi files were imported into Fiji (https://fiji.sc/) using the Simple Neurite tracer v3.1.3 plugin (http://imagej.net/Simple_Neurite_Tracer), first developed by (Longair et al., 2011). The radius of the soma was calculated as the radius of an equivalent circle with the same area of the soma obtained from the maximum intensity projection of the confocal z-stacks. When using this plugin, we first picked the location of the center of the soma and then manually traced the 3D trajectory of different neuronal processes. Each process was tagged as axon, basal or dendritic branch so that they could be assigned different conductances. The tracings were converted into .swc and .asc files to be imported in the NEURON simulation environment v7.6 (Hines and Carnevale, 1997). All model and simulation files will be uploaded to the ModelDB database (https://senselab.med.yale.edu/modeldb/, acc. n. 257027). Also, the morphology will be uploaded to the neuromorpho.org database (www.neuromorpho.org), and an interactive “live paper” page will be setup in the Brain Simulation Platform of the Human Brain Project (https://www.humanbrainproject.eu/en/brain-simulation/live-papers/). To implement a CA1 pyramidal neuron model, we used the 3D accurate morphology reconstructed within this work (**Fig.5D**) equipped with the typical somatodendritic distribution of channels observed in rodents hippocampal CA1 pyramidal neurons (Migliore and Shepherd, 2002). The channel kinetics were identical to those used in (Migliore et al., 2018) (ModelDB acc.n. 244688). The AMPA synapse kinetic model was from (Tsodyks et al., 1998) (ModelDB acc.n. 3815), with *tau_1*= 4.5 ms, a peak conductance of 0.2 nS, and a *tau_rec* of 10 ms or 30 ms to model light and dark condition, respectively. The NMDA kinetics was taken from (Gasparini et al., 2004) (ModelDB acc.n. 44050) with the reverse (unbinding) rate β=0.005 ms^−1^ and a peak conductance of 0.7 nS or 0.55 nS to reproduce L- and D-phase conditions, respectively. To mimic the train of synaptic activations caused by extracellular electrical pulses, we added 20 synapses randomly distributed in the apical dendrites between 50 and 300 μm from the soma, synchronously activated every 10-100 ms.

### CellBlender modeling

We performed 3D Monte Carlo reaction-diffusion simulations using the CellBlender bundle for Blender 2.79 (Windows 10), which includes Blender 2.79 and the CellBlender 1.1 add-on (http://mcell.org/). We built an *in silico* representation of the synaptic environment including an axon with a bouton (green), a spine with its parent dendritic branch (gold) and four neighboring astrocytic processes (blue; **Fig. 7L**). The axon had a radius r=125 nm and contained a spherical bouton with r=250 nm. The bouton was positioned 20 nm away from the spine, a distance that matches the typical height of the synaptic cleft. The spine radius was also r=250 nm. The spine head was connected to the dendritic shaft by a thin (r=50 nm), 500 nm long neck. The dendritic shaft was 2 μm long, with r=500 nm. The bouton-spine interface was flanked by four astrocytic processes with radius r=100 nm (Sweeney et al., 2017). Each astrocyte process expressed glutamate transporters with surface density d=10,800 μm^2^ (Lehre and Danbolt, 1998). Each transporter was modeled using a simplified 3-state kinetic scheme based on kinetic models for glutamate transporters developed by (Barbour, 2001) and (Bergles et al., 2002), also used in our own previous work (Sweeney et al., 2017). In this scheme, the glutamate binding, translocation and re-orientation rates were consistent with estimates from the model of (Bergles et al., 2002). The glutamate-unbinding rate was derived from the equation of the apparent steady state affinity for glutamate, with k_m_=13 μM (Barbour, 2001):

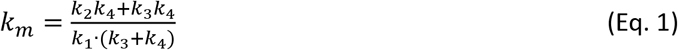

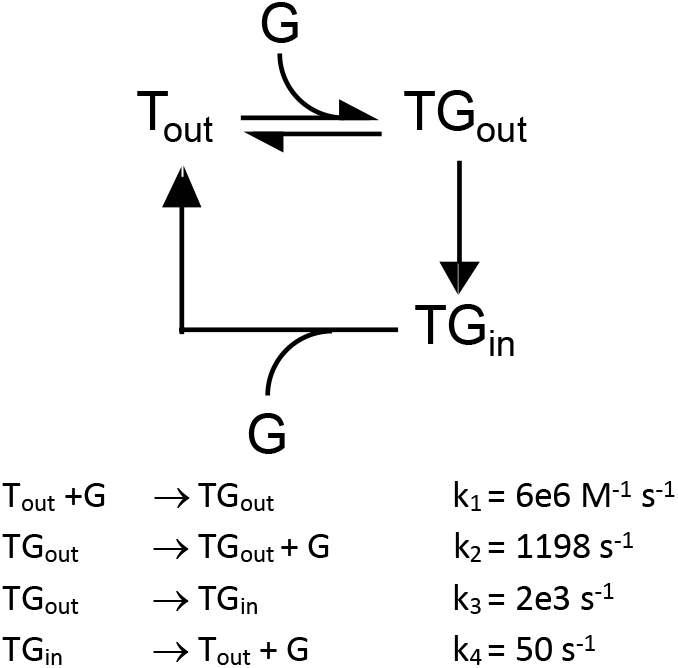

All kinetic rates were adjusted for Q_10_=3, to approximate the receptor and transporter kinetics at physiological temperature (36°C). At the beginning of each simulation, we released 2,000 glutamate molecules from a point source placed in the center of the synaptic cleft. The apparent glutamate diffusion coefficient in the synaptic cleft was 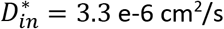 (Nielsen et al., 2004). The hindrance to diffusion within the cleft is mainly due to the interstitial viscosity, estimated as *λ_v_* = 1.2 (Nicholson and Hrabetova, 2017). Therefore *λ_in_* = *λ_v_* = 1.2. The extracellular tortuosity is typically *λ_out_* = 1.6. Because the apparent diffusion coefficient D* is inversely proportionate to *λ*^2^, it follows that:

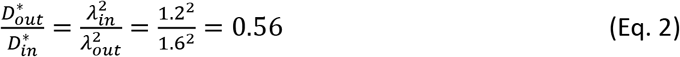

Since 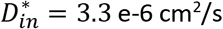, this means that 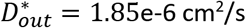. The apparent diffusion coefficient for glutamate changed from 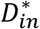 to 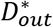 as glutamate diffused away from a 250 nm radius cleft centered at the top of the spine head. Each simulation was run within a 1.6 μm·2.2 μm·2.6 μm world with absorptive properties for glutamate. The results were averaged across 100 seeds (i.e. 100 simulations) using custom made code written in Python 3 (A.Sc.).

### Data analysis

Electrophysiological recordings were analyzed within the Igor Pro environment using custom made software (A.Sc.). Synaptically activated transporter currents (STCs) were isolated as described previously (Diamond, 2005; Scimemi et al., 2009; Sweeney et al., 2017). Briefly, single and pairs of stimuli, 100 ms apart, were delivered every 10 s. We averaged 10 single and 10 STC pairs and subtracted single from paired STCs to isolate the current evoked by the second pulse. The single STC was then shifted in time by 100 ms (the inter-pulse interval), so that its onset matched the onset of the second STC. The current obtained by subtracting the time-shifted single STC from the paired STCs represented the facilitated portion of the STC (fSTC). This analytical strategy allowed us to get rid of the sustained potassium current that is superimposed on the STC but does not facilitate with repeated stimulation. In the unlikely event that a component of the sustained potassium current was still present after this subtraction, we removed it by subtracting a mono-exponential function with an onset time of 4 ms and an amplitude that was scaled to match the amplitude of the remaining potassium current (Scimemi et al., 2009). We used the fSTCs in control conditions and in the presence of a sub-saturating concentration of TFB-TBOA (1 μM) to calculate the time course of glutamate clearance by performing a deconvolution analysis of the STCs (Diamond, 2005; Scimemi et al., 2009; Sweeney et al., 2017). *First*, the STCs recorded under control conditions and in TFB-TBOA (1 μM) were binomially smoothed and fitted with the following equation (Nielsen et al., 2004):

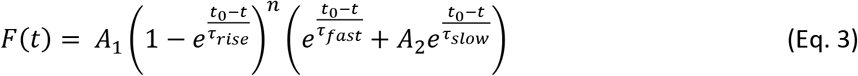

*Second*, we approximated glutamate clearance in TFB-TBOA (1 μM) with an instantaneously rising function decaying mono-exponentially, with the same time course of the STC. *Third*, we deconvolved this approximated glutamate clearance from the STC recorded in low TFB-TBOA to obtain the filter. *Fourth*, we deconvolved the filter from the STC recorded in control conditions to obtain the glutamate clearance waveform in control conditions.

Miniature events were detected using an optimally scaled template (Clements and Bekkers, 1997) adapted for Igor Pro (A.Sc.). Paired-pulse ratio (PPR) of EPSCs was calculated by subtracting single from paired EPSCs averaged across 20 traces, after first peak normalization. When delivering trains of stimuli, we interleaved single pulses with trains of four and five pulses. The 5^th^ EPSC was isolated by subtracting the average response to four pulses from the average response to five pulses.

### Image preprocessing for Sholl analysis

The Sholl analysis was performed on adaptively thresholded maximum intensity projections of filtered confocal z--stacks. Each plane of the z-stack was first filtered with a coherence-enhancing diffusion filter (Weickert and Scharr, 2002), and then we filtered the maximal projection intensity image as described below. Astrocytic processes are generally thin and elongated. To take advantage of this property, we used steerable oriented filters and stored in each pixel the maximal response to different filter sets. Any region near a round bright spot is “oriented” towards it, meaning that the maximal response is obtained with respect to the filter with similar orientation. We used a round isotropic Gaussian filter to find local intensity maxima and oriented filters to highlight the thin elongated structures. To suppress the algorithm from detecting false-positive filamentous structures in dark and noisy regions of interest we performed a randomization-based regularization consisting of many iterations in which we disturbed the input image with noise and then thresholded the responses. By averaging the results obtained over multiple iterations, we obtained an estimate of the probability that each pixel has an oriented structure brighter than its surroundings. The final mask is the product of the isotropic and anisotropic filter responses.

### Image analysis of complexity and entropy

The complexity-entropy analysis allows quantification of the spatial properties of representative images with respect to their balance between randomness and structural order, triviality and complexity as previously described (Brazhe, 2018). In mathematical terms, this is described as location in the complexity-entropy plane or their distribution along a complexity-entropy spectrum. Highly ordered structures (e.g., a regular grating) have near-zero entropy and near-zero complexity. In contrast, completely disordered structures (e.g., independent and identically distributed Gaussian samples) have maximal entropy and small statistical complexity. Intermediate values of entropy are associated with higher values of complexity if the underlying pattern contains features with preferred orientation. For example, signals generated by systems with deterministic chaos result in a complexity-entropy spectrum with near-0.5 peak complexity and near-0.7 entropy (Lamberti et al., 2004; Rosso et al., 2007). The implementation code is available at DOI:10.5281/zenodo.1217636. Briefly, complexity and entropy measures can be based on a feature distribution of the analyzed patterns, which can be compared to equally probable or singular feature cases. Accordingly, the relative entropy (H) was defined as the Shannon entropy:

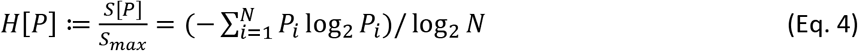

normalized by the entropy of an equally probable distribution (*S*_max_), thus giving values in the range [0 … 1]. Here *N* is the number of features analyzed. The complexity measures were based on the notion of disequilibrium (Lamberti et al., 2004) and defined in terms of normalized Jensen-Shannon divergence from an equally probable distribution (Rosso et al., 2007):

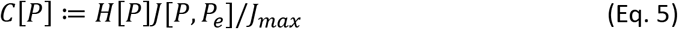

In this expression, *J*[*P*, *P_e_*] is Jensen-Shannon divergence of some distribution P from an equally probable distribution *P_e_*:

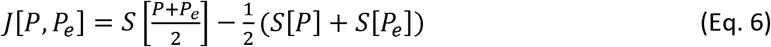

and J_max_ is obtained when the probability of just one feature is equal to 1, while the probability of all other features is zero. The defined probability-based definitions of entropy and complexity require building descriptive probability-like measures for spatial patterns, preferably allowing for multiscale resolution and local analysis. A spatial pattern can be described in terms of local edge orientations and scales, which can be achieved by discrete shearlet transform, which is analogous to wavelet transform with added orientation information. We treated normalized power of shearlet coefficients as probability densities of given spatial feature orientation and scale at a given location. Complexity and entropy calculations were only performed in regions containing non-zero pixels, while other areas were masked out.

### Novel object recognition test

We performed novel object recognition test on C56BL/6 mice of either sex, maintained on as 12H:12H L:D or D:D cycle. The mice were aged P16 on the first training session. The task was conducted either at ZT3.5 or ZT15.5 in a white Plexiglas box (L 46 cm × W 46 cm × H 38 cm). Each mouse was video monitored for 10 min at 30 fps using a Night Vision CMOS IR USB 2.0 webcam (Model #B07PNHJP8J from webcamera_usb). We analyzed the position of the mouse in four adjacent quadrants of the arena (top left, top right, bottom left, bottom right) using ezTrack (Pennington et al., 2019). The test including one habituation session, five training sessions and one testing session. Briefly, during the first habituation session, we positioned a mouse in the open field arena and allowed it to freely explore it for 10 min. During each training session, we re-introduced the mouse in the open field arena, after positioning a nonfamiliar object in its top right corner. The novel object was a metal parallelepiped. During the test session, we removed the object from the arena and allowed the mouse to explore the open field. We calculated the proportion of time spent in the top right corner of the arena during each of the seven sessions using ezTrack. The learning discrimination index was calculated as the difference in the proportion of time spent in the top right corner between the second and first session. The memory discrimination index was calculated as the difference in the proportion of time spent in the top right corner of the arena between the seventh and sixth session.

### Statistical analysis

Data are presented as mean±SEM unless otherwise stated. All experiments were performed on multiple mice of either sex. Statistical significance was determined by Student’s paired or unpaired t test or ANOVA, as appropriate (IgorPro 6.37). Differences were considered significant at p<0.05 (*p<0.05; **p<0.01; ***p<0.001).

## References

Antunes, M., and Biala, G. (2012). The novel object recognition memory: neurobiology, test procedure, and its modifications. Cogn Process 13, 93–110.

Attwell, D., and Laughlin, S.B. (2001). An energy budget for signaling in the grey matter of the brain. J Cereb Blood Flow Metab 21, 1133–1145.

Bae, K., Jin, X., Maywood, E.S., Hastings, M.H., Reppert, S.M., and Weaver, D.R. (2001). Differential functions of mPer1, mPer2, and mPer3 in the SCN circadian clock. Neuron 30, 525–536.

Balsalobre, A., Brown, S.A., Marcacci, L., Tronche, F., Kellendonk, C., Reichardt, H.M., Schutz, G., and Schibler, U. (2000). Resetting of circadian time in peripheral tissues by glucocorticoid signaling. Science 289, 2344–2347.

Barbour, B. (2001). An evaluation of synapse independence. J Neurosci 21, 7969–7984.

Barnes, C.A., McNaughton, B.L., Goddard, G.V., Douglas, R.M., and Adamec, R. (1977). Circadian rhythm of synaptic excitability in rat and monkey central nervous system. Science 197, 91–92.

Becquet, D., Girardet, C., Guillaumond, F., Francois-Bellan, A.M., and Bosler, O. (2008). Ultrastructural plasticity in the rat suprachiasmatic nucleus. Possible involvement in clock entrainment. Glia 56, 294–305.

Benitez-King, G., Hernandez, M.E., Tovar, R., and Ramirez, G. (2001). Melatonin activates PKC-alpha but not PKC-epsilon in N1E-115 cells. Neurochemistry International 39, 95–102.

BenitezKing, G., Rios, A., Martinez, A., and AntonTay, F. (1996). In vitro inhibition of Ca2+/calmodulin-dependent kinase II activity by melatonin. Bba-Gen Subjects 1290, 191–196.

Bergles, D.E., Tzingounis, A.V., and Jahr, C.E. (2002). Comparison of coupled and uncoupled currents during glutamate uptake by GLT-1 transporters. J Neurosci 22, 10153–10162.

Brancaccio, M., Edwards, M.D., Patton, A.P., Smyllie, N.J., Chesham, J.E., Maywood, E.S., and Hastings, M.H. (2019). Cell-autonomous clock of astrocytes drives circadian behavior in mammals. Science 363, 187-+.

Brancaccio, M., Patton, A.P., Chesham, J.E., Maywood, E.S., and Hastings, M.H. (2017). Astrocytes Control Circadian Timekeeping in the Suprachiasmatic Nucleus via Glutamatergic Signaling. Neuron 93, 1420–1435 e1425.

Brazhe, A. (2018). Shearlet-based measures of entropy and complexity for two-dimensional patterns. Phys Rev E 97.

Broadbent, N.J., Gaskin, S., Squire, L.R., and Clark, R.E. (2010). Object recognition memory and the rodent hippocampus. Learn Mem 17, 5–11.

Buzsaki, G. (2002). Theta oscillations in the hippocampus. Neuron 33, 325–340.

Carlberg, C., and Wiesenberg, I. (1995). The Orphan Receptor Family Rzr/Ror, Melatonin and 5-Lipoxygenase - an Unexpected Relationship. J Pineal Res 18, 171–178.

Chaudhury, D., and Colwell, C.S. (2002). Circadian modulation of learning and memory in fear-conditioned mice. Behav Brain Res 133, 95–108.

Chung, S., Son, G.H., and Kim, K. (2011). Circadian rhythm of adrenal glucocorticoid: its regulation and clinical implications. Biochim Biophys Acta 1812, 581–591.

Colgin, L.L., Denninger, T., Fyhn, M., Hafting, T., Bonnevie, T., Jensen, O., Moser, M.B., and Moser, E.I. (2009). Frequency of gamma oscillations routes flow of information in the hippocampus. Nature 462, 353–357.

Collins, D.R., and Davies, S.N. (1997). Melatonin blocks the induction of long-term potentiation in an N-methyl-D-aspartate independent manner. Brain Res 767, 162–165.

Combe, C.L., Canavier, C.C., and Gasparini, S. (2018). Intrinsic Mechanisms of Frequency Selectivity in the Proximal Dendrites of CA1 Pyramidal Neurons. J Neurosci 38, 8110–8127.

Cotte, Y., Toy, F., Jourdain, P., Pavillon, N., Boss, D., Magistretti, P., Marquet, P., and Depeursinge, C. (2013). Marker-free phase nanoscopy. Nat Photonics 7, 113–117.

Debski, K.J., Ceglia, N., Ghestem, A., Ivanov, A.I., Brancati, G.E., Bröer, S., Bot, A.M., Müller, J.A., Schoch, S., Becker, A., et al. (2017). The circadian hippocampus and its reprogramming in epilepsy: impact for chronotherapeutics. bioRxiv.

Diamond, J.S. (2005). Deriving the glutamate clearance time course from transporter currents in CA1 hippocampal astrocytes: transmitter uptake gets faster during development. J Neurosci 25, 2906–2916.

Dickmeis, T. (2009). Glucocorticoids and the circadian clock. J Endocrinol 200, 3–22.

Dodge, F.A., Jr., and Rahamimoff, R. (1967). On the relationship between calcium concentration and the amplitude of the end-plate potential. J Physiol 189, 90P–92P.

Dubocovich, M.L. (2007). Melatonin receptors: Role on sleep and circadian rhythm regulation. Sleep Med 8, S34–S42.

Dubocovich, M.L., Rivera-Bermudez, M.A., Gerdin, M.J., and Masana, M.I. (2003). Molecular pharmacology, regulation and function of mammalian melatonin receptors. Front Biosci 8, d1093–1108.

Ebihara, S., Marks, T., Hudson, D.J., and Menaker, M. (1986). Genetic-Control of Melatonin Synthesis in the Pineal-Gland of the Mouse. Science 231, 491–493.

Engert, F., and Bonhoeffer, T. (1999). Dendritic spine changes associated with hippocampal long-term synaptic plasticity. Nature 399, 66–70.

Ennaceur, A., and Delacour, J. (1988). A new one-trial test for neurobiological studies of memory in rats. 1: Behavioral data. Behav Brain Res 31, 47–59.

Gasparini, S., Migliore, M., and Magee, J.C. (2004). On the initiation and propagation of dendritic spikes in CA1 pyramidal neurons. J Neurosci 24, 11046–11056.

Gavrilov, N., Golyagina, I., Brazhe, A., Scimemi, A., Turlapov, V., and Semyanov, A. (2018). Astrocytic Coverage of Dendritic Spines, Dendritic Shafts, and Axonal Boutons in Hippocampal Neuropil. Front Cell Neurosci 12, 248.

Gerstner, J.R., Lyons, L.C., Wright, K.P., Jr., Loh, D.H., Rawashdeh, O., Eckel-Mahan, K.L., and Roman, G.W. (2009). Cycling behavior and memory formation. J Neurosci 29, 12824–12830.

Gilbert, C.D. (1998). Adult cortical dynamics. Physiological Reviews 78, 467–485.

Guilding, C., and Piggins, H.D. (2007). Challenging the omnipotence of the suprachiasmatic timekeeper: are circadian oscillators present throughout the mammalian brain? Eur J Neurosci 25, 3195–3216.

Guo, K., and Labate, D. (2007). Optimally sparse multidimensional representation using shearlets. Siam J Math Anal 39, 298–318.

Haber, M., Zhou, L., and Murai, K.K. (2006). Cooperative astrocyte and dendritic spine dynamics at hippocampal excitatory synapses. Journal of Neuroscience 26, 8881–8891.

Harris, K.M., and Teyler, T.J. (1983). Age differences in a circadian influence on hippocampal LTP. Brain Res 261, 69–73.

Hines, M.L., and Carnevale, N.T. (1997). The NEURON simulation environment. Neural Comput 9, 1179–1209.

Hirrlinger, J., Hulsmann, S., and Kirchhoff, F. (2004). Astroglial processes show spontaneous motility at active synaptic terminals in situ. European Journal of Neuroscience 20, 2235–2239.

Hohmann-Marriott, M.F., Sousa, A.A., Azari, A.A., Glushakova, S., Zhang, G., Zimmerberg, J., and Leapman, R.D. (2009). Nanoscale 3D cellular imaging by axial scanning transmission electron tomography. Nat Methods 6, 729–731.

Holtmaat, A.J.G.D., Trachtenberg, J.T., Wilbrecht, L., Shepherd, G.M., Zhang, X.Q., Knott, G.W., and Svoboda, K. (2005). Transient and persistent dendritic spines in the neocortex in vivo. Neuron 45, 279–291.

Huettner, J.E. (1989). Indole-2-Carboxylic Acid - a Competitive Antagonist of Potentiation by Glycine at the Nmda Receptor. Science 243, 1611–1613.

Hui, G.K., Figueroa, I.R., Poytress, B.S., Roozendaal, B., McGaugh, J.L., and Weinberger, N.M. (2004). Memory enhancement of classical fear conditioning by post-training injections of corticosterone in rats. Neurobiol Learn Mem 81, 67–74.

Ishida, A., Mutoh, T., Ueyama, T., Bando, H., Masubuchi, S., Nakahara, D., Tsujimoto, G., and Okamura, H. (2005). Light activates the adrenal gland: Timing of gene expression and glucocorticoid release. Cell Metabolism 2, 297–307.

Jiao, Y.Y., Lee, T.M., and Rusak, B. (1999). Photic responses of suprachiasmatic area neurons in diurnal degus (Octodon degus) and nocturnal rats (Rattus norvegicus). Brain Research 817, 93–103.

Jonas, P., Major, G., and Sakmann, B. (1993). Quantal Components of Unitary Epscs at the Mossy Fiber Synapse on Ca3 Pyramidal Cells of Rat Hippocampus. J Physiol-London 472, 615–663.

Kalsbeek, A., and Buijs, R.M. (2002). Output pathways of the mammalian suprachiasmatic nucleus: coding circadian time by transmitter selection and specific targeting. Cell Tissue Res 309, 109–118.

Kasahara, T., Abe, K., Mekada, K., Yoshiki, A., and Kato, T. (2010). Genetic variation of melatonin productivity in laboratory mice under domestication. Proc Natl Acad Sci U S A 107, 6412–6417.

Katagiri, H., Tanaka, K., and Manabe, T. (2001). Requirement of appropriate glutamate concentrations in the synaptic cleft for hippocampal LTP induction. European Journal of Neuroscience 14, 547–553.

Kim, J.J., and Yoon, K.S. (1998). Stress: metaplastic effects in the hippocampus. Trends Neurosci 21, 505–509.

Korf, H.W., and von Gall, C. (2006). Mice, melatonin and the circadian system. Mol Cell Endocrinol 252, 57–68.

Kwapis, J.L., Alaghband, Y., Kramar, E.A., Lopez, A.J., Vogel Ciernia, A., White, A.O., Shu, G., Rhee, D., Michael, C.M., Montellier, E., et al. (2018). Epigenetic regulation of the circadian gene Per1 contributes to age-related changes in hippocampal memory. Nat Commun 9, 3323.

Kwon, H.B., and Sabatini, B.L. (2011). Glutamate induces de novo growth of functional spines in developing cortex. Nature 474, 100–104.

Lamberti, P.W., Martin, M.T., Plastino, A., and Rosso, O.A. (2004). Intensive entropic non-triviality measure. Physica A 334, 119–131.

Lavialle, M., Aumann, G., Anlauf, E., Prols, F., Arpin, M., and Derouiche, A. (2011). Structural plasticity of perisynaptic astrocyte processes involves ezrin and metabotropic glutamate receptors. P Natl Acad Sci USA 108, 12915–12919.

Lehman, M.N., Silver, R., Gladstone, W.R., Kahn, R.M., Gibson, M., and Bittman, E.L. (1987). Circadian rhythmicity restored by neural transplant. Immunocytochemical characterization of the graft and its integration with the host brain. J Neurosci 7, 1626–1638.

Lehre, K.P., and Danbolt, N.C. (1998). The number of glutamate transporter subtype molecules at glutamatergic synapses: chemical and stereological quantification in young adult rat brain. J Neurosci 18, 8751–8757.

Lester, R.A.J., and Jahr, C.E. (1992). Nmda Channel Behavior Depends on Agonist Affinity. Journal of Neuroscience 12, 635–643.

Longair, M.H., Baker, D.A., and Armstrong, J.D. (2011). Simple Neurite Tracer: open source software for reconstruction, visualization and analysis of neuronal processes. Bioinformatics 27, 2453–2454.

Marr, D. (1971). Simple memory: a theory for archicortex. Philos Trans R Soc Lond B Biol Sci 262, 23–81.

McEown, K., and Treit, D. (2011). Mineralocorticoid receptors in the medial prefrontal cortex and hippocampus mediate rats’ unconditioned fear behaviour. Horm Behav 60, 581–588.

Meijer, J.H., Watanabe, K., Schaap, J., Albus, H., and Detari, L. (1998). Light responsiveness of the suprachiasmatic nucleus: long-term multiunit and single-unit recordings in freely moving rats. J Neurosci 18, 9078–9087.

Meyer-Bernstein, E.L., Jetton, A.E., Matsumoto, S.I., Markuns, J.F., Lehman, M.N., and Bittman, E.L. (1999). Effects of suprachiasmatic transplants on circadian rhythms of neuroendocrine function in golden hamsters. Endocrinology 140, 207–218.

Migliore, M., De Simone, G., and Migliore, R. (2015). Effect of the initial synaptic state on the probability to induce long-term potentiation and depression. Biophys J 108, 1038–1046.

Migliore, M., and Shepherd, G.M. (2002). Emerging rules for the distributions of active dendritic conductances. Nat Rev Neurosci 3, 362–370.

Migliore, R., Lupascu, C.A., Bologna, L.L., Romani, A., Courcol, J.D., Antonel, S., Van Geit, W.A.H., Thomson, A.M., Mercer, A., Lange, S., et al. (2018). The physiological variability of channel density in hippocampal CA1 pyramidal cells and interneurons explored using a unified data-driven modeling workflow. PLoS Comput Biol 14, e1006423.

Moore, R.Y. (1996). Neural control of the pineal gland. Behav Brain Res 73, 125–130.

Moore, R.Y., and Lenn, N.J. (1972). A retinohypothalamic projection in the rat. J Comp Neurol 146, 1–14.

Musshoff, U., Riewenherm, D., Berger, E., Fauteck, J.D., and Speckmann, E.J. (2000). Melatonin receptors in rat hippocampus: Molecular and functional investigations. European Journal of Neuroscience 12, 481–481.

Namihira, M., Honma, S., Abe, H., Tanahashi, Y., Ikeda, M., and Honma, K. (1999). Daily variation and light responsiveness of mammalian clock gene, Clock and BMAL1, transcripts in the pineal body and different areas of brain in rats. Neurosci Lett 267, 69–72.

Nicholson, C., and Hrabetova, S. (2017). Brain Extracellular Space: The Final Frontier of Neuroscience. Biophys J 113, 2133–2142.

Nielsen, T.A., DiGregorio, D.A., and Silver, R.A. (2004). Modulation of glutamate mobility reveals the mechanism underlying slow-rising AMPAR EPSCs and the diffusion coefficient in the synaptic cleft. Neuron 42, 757–771.

Nishida, H., and Okabe, S. (2007). Direct astrocytic contacts regulate local maturation of dendritic spines. Journal of Neuroscience 27, 331–340.

O’Donnell, C., and van Rossum, M.C. (2014). Systematic analysis of the contributions of stochastic voltage gated channels to neuronal noise. Front Comput Neurosci 8, 105.

Ozcan, M., Yilmaz, B., and Carpenter, D.O. (2006). Effects of melatonin on synaptic transmission and longterm potentiation in two areas of mouse hippocampus. Brain Research 1111, 90–94.

Papouin, T., Dunphy, J.M., Tolman, M., Dineley, K.T., and Haydon, P.G. (2017). Septal Cholinergic Neuromodulation Tunes the Astrocyte-Dependent Gating of Hippocampal NMDA Receptors to Wakefulness. Neuron 94, 840–854 e847.

Pavlides, C., Watanabe, Y., Magarinos, A.M., and McEwen, B.S. (1995). Opposing roles of type I and type II adrenal steroid receptors in hippocampal long-term potentiation. Neuroscience 68, 387–394.

Peirson, S.N., Brown, L.A., Pothecary, C.A., Benson, L.A., and Fisk, A.S. (2018). Light and the laboratory mouse. J Neurosci Meth 300, 26–36.

Pennington, Z.T., Dong, Z., Feng, Y., Vetere, L.M., Page-Harley, L., Shuman, T., and Cai, D.J. (2019). ezTrack: An open-source video analysis pipeline for the investigation of animal behavior. Sci Rep 9, 19979.

Prolo, L.M., Takahashi, J.S., and Herzog, E.D. (2005). Circadian rhythm generation and entrainment in astrocytes. Journal of Neuroscience 25, 404–408.

Qi, X.R., Kamphuis, W., Wang, S., Wang, Q., Lucassen, P.J., Zhou, J.N., and Swaab, D.F. (2013). Aberrant stress hormone receptor balance in the human prefrontal cortex and hypothalamic paraventricular nucleus of depressed patients. Psychoneuroendocrinology 38, 863–870.

Quirk, J.C., and Nisenbaum, E.S. (2002). LY404187: a novel positive allosteric modulator of AMPA receptors. CNS Drug Rev 8, 255–282.

Rafii-El-Idrissi, M., Calvo, J.R., Harmouch, A., Garcia-Maurino, S., and Guerrero, J.M. (1998). Specific binding of melatonin by purified cell nuclei from spleen and thymus of the rat. J Neuroimmunol 86, 190–197.

Raghavan, A.V., Horowitz, J.M., and Fuller, C.A. (1999). Diurnal modulation of long-term potentiation in the hamster hippocampal slice. Brain Res 833, 311–314.

Rakic, P., Bourgeois, J.P., Eckenhoff, M.F., Zecevic, N., and Goldmanrakic, P.S. (1986). Concurrent Overproduction of Synapses in Diverse Regions of the Primate Cerebral-Cortex. Science 232, 232–235.

Rawashdeh, O., de Borsetti, N.H., Roman, G., and Cahill, G.M. (2007). Melatonin suppresses nighttime memory formation in zebrafish. Science 318, 1144–1146.

Rawashdeh, O., Parsons, R., and Maronde, E. (2018). Clocking In Time to Gate Memory Processes: The Circadian Clock Is Part of the Ins and Outs of Memory. Neural Plast.

Reichenbach, A., Derouiche, A., and Kirchhoff, F. (2010). Morphology and dynamics of perisynaptic glia. Brain Research Reviews 63, 11–25.

Reick, M., Garcia, J.A., Dudley, C., and McKnight, S.L. (2001). NPAS2: an analog of clock operative in the mammalian forebrain. Science 293, 506–509.

Reppert, S.M., Weaver, D.R., and Godson, C. (1996). Melatonin receptors step into the light: Cloning and classification of subtypes. Trends Pharmacol Sci 17, 100–102.

Richards, D.A., Mateos, J.M., Hugel, S., de Paola, V., Caroni, P., Gahwiler, B.H., and McKinney, R.A. (2005). Glutamate induces the rapid formation of spine head protrusions in hippocampal slice cultures. P Natl Acad Sci USA 102, 6166–6171.

Rosenfeld, P., Van Eekelen, J.A., Levine, S., and De Kloet, E.R. (1988). Ontogeny of the type 2 glucocorticoid receptor in discrete rat brain regions: an immunocytochemical study. Brain Res 470, 119–127.

Rosso, O.A., Larrondo, H.A., Martin, M.T., Plastino, A., and Fuentes, M.A. (2007). Distinguishing noise from chaos. Phys Rev Lett 99.

Ruby, N.F., Hwang, C.E., Wessells, C., Fernandez, F., Zhang, P., Sapolsky, R., and Heller, H.C. (2008). Hippocampal-dependent learning requires a functional circadian system. P Natl Acad Sci USA 105, 15593–15598.

Salituro, F.G., Harrison, B.L., Baron, B.M., Nyce, P.L., Stewart, K.T., Kehne, J.H., White, H.S., and Mcdonald, I.A. (1992). 3-(2-Carboxyindol-3-Yl)Propionic Acid-Based Antagonists of the N-Methyl-D-Aspartic Acid Receptor Associated Glycine Binding-Site. J Med Chem 35, 1791–1799.

Salituro, F.G., Harrison, B.L., Baron, B.M., Nyce, P.L., Stewart, K.T., and Mcdonald, I.A. (1990). 3-(2-Carboxyindol-3-Yl)Propionic Acid-Derivatives - Antagonists of the Strychnine-Insensitive Glycine Receptor Associated with the N-Methyl-D-Aspartate Receptor Complex. J Med Chem 33, 2944–2946.

Saunders, A., Macosko, E.Z., Wysoker, A., Goldman, M., Krienen, F.M., de Rivera, H., Bien, E., Baum, M., Bortolin, L., Wang, S.Y., et al. (2018). Molecular Diversity and Specializations among the Cells of the Adult Mouse Brain. Cell 174, 1015-+.

Scimemi, A., and Diamond, J.S. (2013). Deriving the time course of glutamate clearance with a deconvolution analysis of astrocytic transporter currents. J Vis Exp.

Scimemi, A., Fine, A., Kullmann, D.M., and Rusakov, D.A. (2004). NR2B-containing receptors mediate cross talk among hippocampal synapses. J Neurosci 24, 4767–4777.

Scimemi, A., Meabon, J.S., Woltjer, R.L., Sullivan, J.M., Diamond, J.S., and Cook, D.G. (2013). Amyloid-beta1-42 slows clearance of synaptically released glutamate by mislocalizing astrocytic GLT-1. J Neurosci 33, 5312–5318.

Scimemi, A., Tian, H., and Diamond, J.S. (2009). Neuronal transporters regulate glutamate clearance, NMDA receptor activation, and synaptic plasticity in the hippocampus. J Neurosci 29, 14581–14595.

Shimizu, K., Kobayashi, Y., Nakatsuji, E., Yamazaki, M., Shimba, S., Sakimura, K., and Fukada, Y. (2016). SCOP/PHLPP1beta mediates circadian regulation of long-term recognition memory. Nat Commun 7, 12926.

Shiromani, P.J., Xu, M., Winston, E.M., Shiromani, S.N., Gerashchenko, D., and Weaver, D.R. (2004). Sleep rhythmicity and homeostasis in mice with targeted disruption of mPeriod genes. Am J Physiol Regul Integr Comp Physiol 287, R47–57.

Smale, L., Lee, T., and Nunez, A.A. (2003). Mammalian diurnality: Some facts and gaps. J Biol Rhythm 18, 356–366.

Smarr, B.L., Jennings, K.J., Driscoll, J.R., and Kriegsfeld, L.J. (2014). A time to remember: the role of circadian clocks in learning and memory. Behav Neurosci 128, 283–303.

Snider, K.H., Dziema, H., Aten, S., Loeser, J., Norona, F.E., Hoyt, K., and Obrietan, K. (2016). Modulation of learning and memory by the targeted deletion of the circadian clock gene Bmal1 in forebrain circuits. Behav Brain Res 308, 222–235.

Sousa, A.A., Azari, A.A., Zhang, G., and Leapman, R.D. (2011). Dual-axis electron tomography of biological specimens: Extending the limits of specimen thickness with bright-field STEM imaging. J Struct Biol 174, 107–114.

Stephan, F.K., and Zucker, I. (1972). Circadian rhythms in drinking behavior and locomotor activity of rats are eliminated by hypothalamic lesions. Proc Natl Acad Sci U S A 69, 1583–1586.

Sumova, A., Bendova, Z., Sladek, M., El-Hennamy, R., Mateju, K., Polidarova, L., Sosniyenko, S., and Illnerova, H. (2008). Circadian Molecular Clocks Tick along Ontogenesis. Physiol Res 57, S139–S148.

Sumova, A., Sladek, M., Polidarova, L., Novakova, M., and Houdek, P. (2012). Circadian system from conception till adulthood. Prog Brain Res 199, 83–103.

Sweeney, A.M., Fleming, K.E., McCauley, J.P., Rodriguez, M.F., Martin, E.T., Sousa, A.A., Leapman, R.D., and Scimemi, A. (2017). PAR1 activation induces rapid changes in glutamate uptake and astrocyte morphology. Sci Rep 7, 43606.

Thomas-Crusells, J., Vieira, A., Saarma, M., and Rivera, C. (2003). A novel method for monitoring surface membrane trafficking on hippocampal acute slice preparation. J Neurosci Meth 125, 159–166.

Tillberg, P.W., Chen, F., Piatkevich, K.D., Zhao, Y., Yu, C.C., English, B.P., Gao, L., Martorell, A., Suk, H.J., Yoshida, F., et al. (2016). Protein-retention expansion microscopy of cells and tissues labeled using standard fluorescent proteins and antibodies. Nat Biotechnol 34, 987–992.

Tsodyks, M., Pawelzik, K., and Markram, H. (1998). Neural networks with dynamic synapses. Neural Comput 10, 821–835.

Vanecek, J. (1998). Cellular mechanisms of melatonin action. Physiol Rev 78, 687–721.

Venkova, K., Foreman, R.D., and Greenwood-Van Meerveld, B. (2009). Mineralocorticoid and glucocorticoid receptors in the amygdala regulate distinct responses to colorectal distension. Neuropharmacology 56, 514–521.

Wakamatsu, H., Yoshinobu, Y., Aida, R., Moriya, T., Akiyama, M., and Shibata, S. (2001). Restricted-feeding-induced anticipatory activity rhythm is associated with a phase-shift of the expression of mPer1 and mPer2 mRNA in the cerebral cortex and hippocampus but not in the suprachiasmatic nucleus of mice. European Journal of Neuroscience 13, 1190–1196.

Wang, L.M., Suthana, N.A., Chaudhury, D., Weaver, D.R., and Colwell, C.S. (2005). Melatonin inhibits hippocampal long-term potentiation. Eur J Neurosci 22, 2231–2237.

Wang, L.M.C., Dragich, J.M., Kudo, T., Odom, I.H., Welsh, D.K., O’Dell, T.J., and Colwell, C.S. (2009). Expression of the circadian clock gene Period2 in the hippocampus: possible implications for synaptic plasticity and learned behaviour. Asn Neuro 1.

Weickert, J., and Scharr, H. (2002). A scheme for coherence-enhancing diffusion filtering with optimized rotation invariance. J Vis Commun Image R 13, 103–118.

Weinert, D. (2005). Ontogenetic development of the mammalian circadian system. Chronobiol Int 22, 179–205.

Whitehead, G., Jo, J., Hogg, E.L., Piers, T., Kim, D.H., Seaton, G., Seok, H., Bru-Mercier, G., Son, G.H., Regan, P., et al. (2013). Acute stress causes rapid synaptic insertion of Ca2+-permeable AMPA receptors to facilitate long-term potentiation in the hippocampus. Brain 136, 3753–3765.

Willshaw, D.J., Dayan, P., and Morris, R.G. (2015). Memory, modelling and Marr: a commentary on Marr (1971) ‘Simple memory: a theory of archicortex’. Philos Trans R Soc Lond B Biol Sci 370.

Winson, J., and Abzug, C. (1977). Gating of neuronal transmission in the hippocampus: efficacy of transmission varies with behavioral state. Science 196, 1223–1225.

Winson, J., and Abzug, C. (1978). Neuronal transmission through hippocampal pathways dependent on behavior. J Neurophysiol 41, 716–732.

Yuen, E.Y., Liu, W., Karatsoreos, I.N., Ren, Y., Feng, J., McEwen, B.S., and Yan, Z. (2011). Mechanisms for acute stress-induced enhancement of glutamatergic transmission and working memory. Mol Psychiatr 16, 156–170.

Zerangue, N., and Kavanaugh, M.P. (1996). Flux coupling in a neuronal glutamate transporter. Nature 383, 634–637.

Zheng, B., Albrecht, U., Kaasik, K., Sage, M., Lu, W., Vaishnav, S., Li, Q., Sun, Z.S., Eichele, G., Bradley, A., et al. (2001). Nonredundant roles of the mPer1 and mPer2 genes in the mammalian circadian clock. Cell 105, 683–694.

Zheng, C., Bieri, K.W., Hwaun, E., and Colgin, L.L. (2016). Fast Gamma Rhythms in the Hippocampus Promote Encoding of Novel Object-Place Pairings. eNeuro 3.

